# Low-Rank Full Matrix Factorization for dropout imputation in single cell RNA-seq and benchmarking with imputation algorithms for downstream applications

**DOI:** 10.1101/2024.10.21.619343

**Authors:** Jinghan Huang, Anson C.M. Chow, Nelson L.S. Tang, Sheung Chi Phillip Yam

## Abstract

**Background:** While single cell RNA sequencing becomes a powerful technology, the presence of the large number of zero counts represents a challenge for both wet-lab processing and data analysis. Imputation of these dropouts can now be performed by three categories of algorithms: Model or smoothing, Matrix theory or Deep learning. However, two fundamental questions remain unsettled: (1) whether imputation should be performed; (2) which imputation algorithm to use with various downstream applications. Notably, imputation is not commonly used in real scRNA-seq applications because of their uncertain benefits, concerns about false inferences in downstream applications, and the lack of in-depth benchmark.

**Methods:** Here, we performed two tasks. First, we developed an algorithm using adaptive low-rank full matrix factorization (afMF) based on a previous limited implementation confined to using low rank matrix decomposition (ALRA). Second, to evaluate the impact of various imputation algorithms on downstream analyses, a new benchmark framework incorporating commonly used downstream applications was developed. This benchmark framework put emphasis on real datasets which had ground truth or matched bulk data such that algorithm performance was compared to more convinced data rather than less realistic simulated parameters.

**Results:** Our results indicated that afMF and ALRA (matrix based) provided good imputation and outperformed raw log-normalization in various downstream applications. afMF outperformed ALRA in several evaluations (cell-level differential expression analysis, GSEA, classification, biomarker prediction, clustering, SC-bulk profiling similarity). Besides, afMF ranked among the top levels in automatic cell type annotation, trajectory inference by DPT, and AUCell & SCENIC. Both showed acceptable scalability, while afMF had longer running time. MAGIC (smoothing based) and AutoClass (deep learning based) also performed well but may produce false positives. In contrast, more complicated methods (other deep learning or model based) were prone to overfitting and data distortion. We also found that certain downstream algorithms are not compatible with imputation, including trajectory inference with Slingshot and cell-cell communication. Prior imputation either showed no improvement or generated false positive findings with these downstream applications.

**Conclusions:** We hope this in-depth evaluation and the algorithm developed in this study can enhance the selection of appropriate imputation algorithm for specific scRNA-seq downstream analyses.

The algorithm and the benchmark framework are available at GitHub: https://github.com/GO3295/SCImputation

## Background

Single cell RNA-seq has been widely used in biomedical research in the last ten years. However, the ‘dropout’ problem in the data, also known as inflated zeros as it results in many zero counts in the data matrix, is still an un-resolved issue. Dropouts (or zero counts) could be due to both biological and technical reasons. As previously discussed(1,2), some researchers believed that the inflated zeros should be imputed so that downstream analyses could benefit from it(3–5), while others insisted that imputations may introduce false-positive signals and those zeros may contain important information as well(6,7). No definitive conclusion has been reached for whether and when imputation should be performed so far, and addressing this question would be beneficial to the biological community using scRNA-seq. From a biological perspective, we may expect no expression of many genes in particular cell types or conditions (biological zero); on the other hand, it is so common to see well-known housekeeping (HK) genes, e.g., GAPDH or UBC, have zero counts in cells caused by technical reasons (technical zero). The technical zeros which do not represent the ground truth, are the prime targets to be handled by various imputation methods.

There are several origins of technical zeros. Technical zero could be due to sampling problem in a finite total number of reads in the setting of scRNA-seq. It could also be a wet-lab technical or measurement error. It is not uncommon to see zero counts for many HK genes in scRNA-seq datasets. Specifically, HK genes account for a large proportion of transcripts inside a cell and they are found universally among various cell types. In fact, HK genes also represent the majority of reads in any single cell inside the scRNA-seq data matrix. Besides, cell type specific marker genes are also present in a high abundance but are only confined in specific cell types. Both kinds of highly expressed genes represent a high proportion of transcripts in a single cell. When random dropout (technical zero) happens to these genes, the downstream normalization and subsequent analysis will be adversely affected. For example, there is a heavy zero tail for B cells, CD4+ T cells, and NK cells in the log count distribution of GAPDH in our study. For cell type specific marker genes, CD8A and CD19, they are also missing in significant proportion of T cells and B cells, respectively (**Figure 2A**, **first row**). These technical dropouts need to be taken care of to get more representative results in downstream analysis. Furthermore, the presence of zeros prevents log-transformation of the data and therefore investigators need to compromise by using pseudocount addition rather than using the observed counts in scRNA-seq data analysis. Under these circumstances, imputation would be a more preferred alternative. Nonetheless, imputation is still not a popular approach in real-world applications since investigators have concerns about its effects on downstream applications and its use may probably result in incorrect or false positive inference. A detailed and in-depth benchmark incorporating popular downstream applications is also lack.

A number of imputation algorithms have been developed to recover those technical zeros while keeping the biological signals intact. These algorithms can be classified into three classes: (1) smoothing or model based, where various statistical distributions are assumed; (2) matrix decomposition/factorization; (3) deep learning. For instance, MAGIC(3) is a popular algorithm that is based on data smoothing. ALRA(4) applies matrix decomposition with thresholding and shows a conserved but stable performance. DCA(8) is a deep-learning-based method and has demonstrated some advantages(9). Representative algorithms from each of the three classes are included in this evaluation and benchmarking exercise, so that we may understand performance characteristics of the three classes. In addition, we developed a new algorithm ‘afMF’ (**a**daptively threshold **f**ull **M**atrix **F**actorization) which is an improved version of ALRA as it takes into account of iterative low-rank full matrix factorization and is also evaluated here.

Meanwhile, several comparative studies for dropout imputation have been conducted(1,9–12) but had several limitations: (1) lack of in-depth analysis, i.e., to see the effects with popular downstream applications; (2) limited number of datasets, dataset types and tested algorithms; (3) using biased, unreasonable performance metrics or confined to basic summary statistics only; (4) confined to using many simulated datasets rather than real data. These limitations are also complicated by lack of real datasets with given ground truth. We are using a collection of real datasets with ground truth or near ground truth to address this deficiency. The lack of understanding of imputation in scRNA-seq made investigators to believe that imputation is meaningless as good evidence supporting that it can be beneficial to real applications is still lacking. At the moment, most of the algorithms are not used in any real-world applications or only confined to be used in a limited number of downstream applications (e.g., cell type clustering). A more thorough evaluation of the compatibility between imputation of missing data and all key downstream application of scRNA-seq is required.

In order to have ground truth or near ground truth in real dataset analysis, we preferentially selected datasets from three kinds of experiments, including (1) cell mixture of known cell types and gene expression profile of component cell types that have been well understood by bulk RNA-seq; (2) cell-cycle/time experiment with well-known checkpoint description. By this approach, downstream analyses (e.g., cell clustering) of cell mixture samples processed by scRNA-seq can be confirmed and compared according to characteristics of component cell lines (near ground-truth is available). (3) Datasets with individual cells measured for additional ground truth data required, e.g., protein expression to verify mRNA expression.

We evaluated the compatibility between imputation algorithms and downstream analyses, that is whether the particular downstream application / algorithm is compatible with prior imputation of the count data matrix. The issue of compatibility is obvious when applying downstream algorithms or tools that have in-situ imputation steps, as prior imputation may be unnecessary or make the results worse. There are some downstream tools specially designed for sparse data to account for the zero-inflation problems and thus imputation will be again redundant. As such, many researchers have chosen zero-inflated-model-based tools instead of imputing them in advance, but there are also studies declared that zero-inflated models were not fit for scRNA-seq data(7). And thus, it is required to find out what kind of downstream algorithms are not compatible with prior data imputation. An in-depth understanding of compatibility between prior imputation algorithms and downstream applications is urgently needed to better standardize scRNA-seq routine.

Therefore, motivated by the recent published benchmark reviews(2), we develop an improved benchmark pipeline to address these issues by including the following features (**Figure 1**), including: (1) 20 top or new algorithms with acceptable scalability; (2) more than 20 real (mixture/purified cell type/time-course) or simulated datasets; (3) a pre-screening test to select algorithms for further evaluations; (4) Visualizations (Gene Expression Violin plots; PCA/UMAP plots; Cell-Cell Correlations); (5) Differential Expression (DE) Analysis: Wilcoxon Rank Sum test, MAST(13) and Pseudobulk DE analysis(14); (6) Enrichment Analysis (GSEA)(15,16); (7) Biomarker Prediction and Classification; (8) Automatic Cell Type Annotation: SCINA(17) and scType(18); (9) Dimension Reduction and Clustering; (10) Cell Cycle Dynamics; (11) Pseudotime Trajectory Analysis: Monocle3(19), Slingshot(20) and DPT(21); (12) AUCell & SCENIC(22); (13) Cell-Cell Communication: CellPhoneDB(23) & CellChat(24); (14) Supporting Analysis: Single Cell-Bulk Profiling Similarity & Surface Protein-mRNA Correlation; (15) Running Time, Memory Usage and Recommendation. We used some of the datasets and evaluation metrics developed from previous works(9,10) as their properties have been well described. To our knowledge, this is the first study to perform in-depth benchmark for imputation algorithms incorporating various advanced downstream applications (i.e., pseudobulk DE analysis, GSEA, automatic cell type annotation, Monocle3, Slingshot, DPT, AUCell & SCENIC and cell-cell communication) of scRNA-seq.

**Figure 1.**
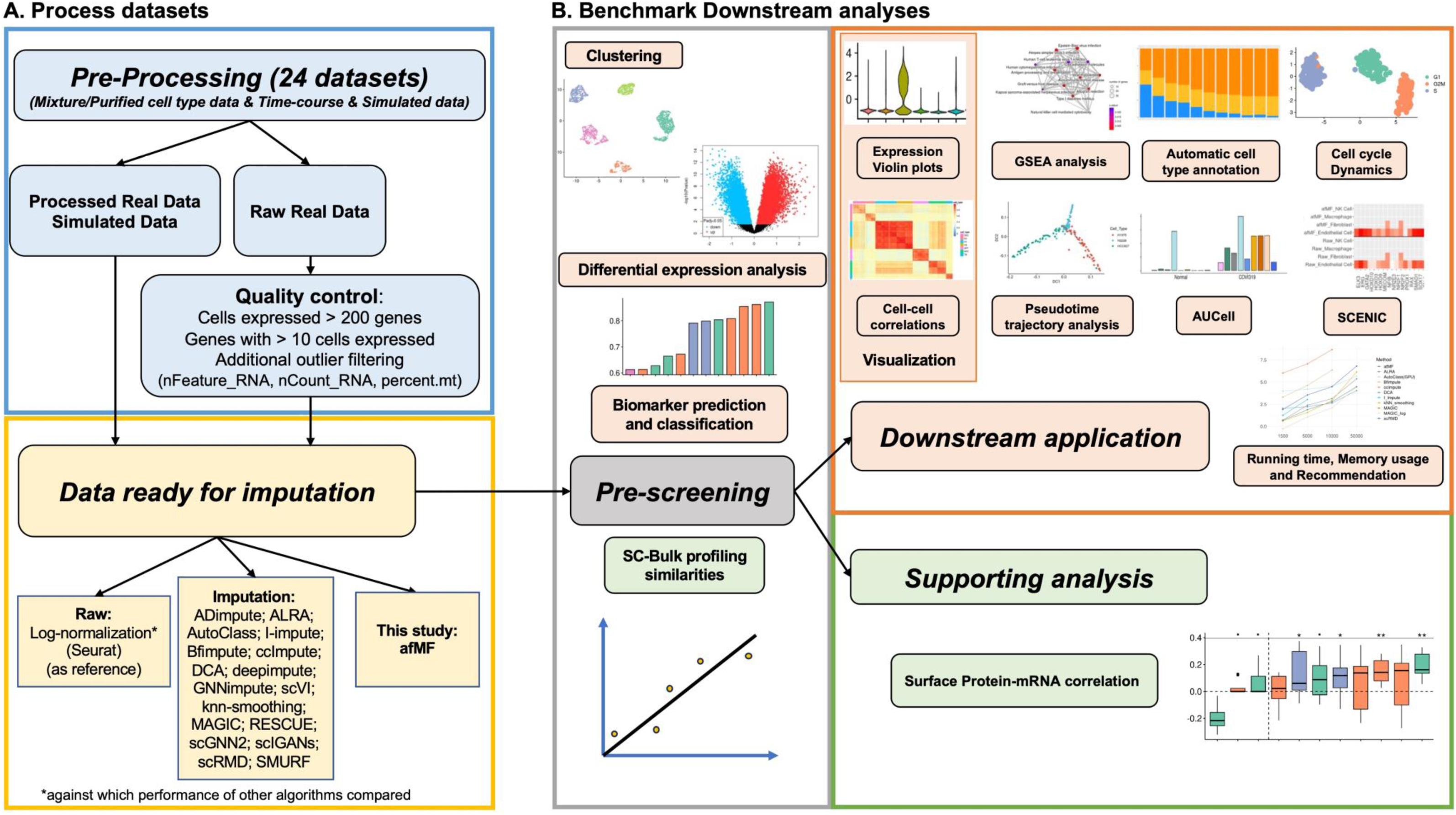
Workflow in this study. Twenty-two datasets including bulk, mixture/purified cell type, time-course and simulated mixture data were pre-processed with quality control, normalization, and imputation. Performances of raw and various imputation methods were evaluated by eleven downstream applications and three types of supporting analyses. A pre-screening test, including Differential Expression Analysis, Biomarker Prediction and Classification, Clustering and Single Cell-Bulk Profiling similarities on two datasets GSE75748 and GSE81861, was performed to initially select the imputation algorithms for in-depth evaluations.

## Methods

### Datasets and Algorithms

Datasets including scRNA-seq, bulk RNA-seq and other related datasets used in this study were collected from multiple public resources(4),(25–41) (**Table 1)**. High-quality datasets that have been widely used in various evaluations and algorithm tests were collected. The simulated datasets with given ground truth were generated by Splatter(42) and SplatPop(43). In some evaluations, datasets were divided into mixture/purified cell type and time-course data as they may have different levels of complexity for imputing.

**Table 1.** Datasets used in this study.

| **Datasets** | **Type** | **Description** | **Evaluation** |
| --- | --- | --- | --- |
| **CellBench 10x 5cl (GSE126906)** | Single cell RNA-seq | Cell mixture sample of 5 cell lines H2228, H1975, A549, H838 and HCC827. | Differential Expression Analysis; GSEA Analysis; Biomarker Prediction & Classification; Clustering; SC-Bulk Similarity. |
| **GSE86337** | Bulk RNA-seq | Bulk samples of 5 cell lines H2228, H1975, HCC827, H838 and A549. | Differential Expression Analysis; GSEA Analysis; Biomarker Prediction; SC-Bulk Similarity. |
| **GSE81861** | Single cell RNA-seq | Cell mixture sample of 5 cell lines A549, GM12878, H1-hESC, IMR90, and K562. | Differential Expression Analysis; GSEA Analysis; Biomarker Prediction & Classification; Clustering; SC-Bulk Similarity. |
| **ENCODE bulk RNA-seq** | Bulk RNA-seq | Bulk samples of 5 cell lines A549, GM12878, H1-hESC, IMR90, and K562. | Differential Expression Analysis; GSEA Analysis; Biomarker Prediction; SC-Bulk Similarity. |
| **GSE75748** | Single cell RNA-seq & Bulk RNA-seq | Cell mixture sample and bulk sample of 7 cell types DEC, EC, H1, H9, HFF, NPC and TB; Time course profiling single cell/bulk sample using H1. | Differential Expression Analysis; GSEA Analysis; Biomarker Prediction & Classification; Clustering; Pseudotime Trajectory Analysis; SC-Bulk Similarity. |
| **GSE124872** | Single cell RNA-seq & Bulk RNA-seq | Alveolar macrophages and type-2 pneumocytes single cell/bulk samples from 24 months and 3 months old mice. | Differential Expression Analysis; GSEA Analysis; Biomarker Prediction & Classification; SC-Bulk Similarity. |
| **EGAS00001003215 and EGAS00001003823** | Single cell RNA-seq & Bulk RNA-seq | Naive or memory T cells (Th0; Th2; Th17; iTreg) single cell/bulk samples: stimulated for 5 d with anti-CD3/anti-CD28 coated beads or resting. | Differential Expression Analysis; GSEA Analysis; Biomarker Prediction & Classification; SC-Bulk Similarity. |
| **GSE79578** | Single cell RNA-seq & Bulk RNA-seq | Mouse embryonic stem cells after different periods of continuous exposure to retinoic acid. | Differential Expression Analysis, Pseudotime Trajectory Analysis; Classification; GSEA analysis. |
| **GSE90047** | Single cell RNA-seq & Bulk RNA-seq | Sorted hepatoblasts, hepatocytes and cholangiocytes from E10.5-E17.5 mouse fetal livers. | Differential Expression Analysis, Pseudotime Trajectory Analysis; Classification; GSEA analysis. |
| **GSE155673** | Single cell RNA-seq | Cells from PBMC with labeled cell types and disease status (COVID-19 and healthy controls). | Differential Expression Analysis & GSEA analysis (real application for COVID-19); Automatic Cell Type Annotation; AUCell. |
| **brain vasculature**<br>http://compbio.mit.edu/scADbbb/ | Single cell RNA-seq | Human brain tissues from the ROSMAP with cell types labeled. | Automatic Cell Type Annotation. |
| **GSE64016** | Single cell RNA-seq | H1 and H1-Fucci single cells | Cell Cycle Dynamics. |
| **CellBench cellmix1-4 (GSE118704)** | Single cell RNA-seq | 9 cell mixtures with differentiation information from three cell lines H2228, H1975 and HCC827. | Pseudotime Trajectory Analysis. |
| **GSE118068** | Single cell RNA-seq | Nine mouse hindbrains from different developmental stages (embryonic days and postnatal days). | Pseudotime Trajectory Analysis. |
| **GSE115978** | Single cell RNA-seq | Cells from fresh tumor resections, isolated immune and non-immune cells by FACS based on CD45 staining. Cell types were pre-labeled. | SCENIC |
| **E-MTAB-6701** | Single cell RNA-seq | Cells from matched first trimester samples of maternal blood and decidua, as well as fetal cells from the placenta itself. | CellPhoneDB |
| **E-MTAB-13384** | Single cell RNA-seq | Cardiac muscle cell from mice with WT/DNMT3AR882H knock-in | CellChat |
| **GSE100866** | Single cell RNA-seq | Cord blood mononuclear cells were profiled by CITE-seq using a panel of 13 antibodies. | Surface Protein-mRNA Correlation |
| **Simulated dataset1 (Mock90)** | Single cell RNA-seq (Simulated) | Splatter: nGenes=10000, batchCells=1500, group.prob=c(0.3,0.25,0.2,0.15,0.1); set dropout.mid to let dropout rate=90%. Others are default. | Differential Expression Analysis; Biomarker Prediction; Classification; Automatic Cell Type Annotation; Clustering; SC-Ground Truth Similarity. |
| **Simulated dataset2 (SplatPop90)** | Single cell RNA-seq (Simulated) | SplatPop: nGenes=8000, batchCells=200, group.prob=c(0.4,0.3,0.3), condition.prob=c(0.5,0.5), similarity.scale=15, de.facLoc=0.5, de.facScale=0.5, cde.facLoc=0.38, cde.facScale=0.38, set dropout.mid to let dropout rate=90%. Others are default. | Differential Expression Analysis; Biomarker Prediction & Classification; Automatic Cell Type Annotation; Clustering. |

Twenty-two real datasets and two simulated datasets (i.e., Splatter and SplatPop) were used in various downstream analyses. In some evaluations, datasets were divided into mixture, purified cell type data and time-course data as they may have different levels of complexity.

To select imputation algorithms for evaluation, we reviewed literatures(1,9,10) and listed twenty of them(3–5,8,44–57) (**Table 2**). Basically, these algorithms were either demonstrated to have top performance in previous evaluations or were new algorithms with good design and documentation. Imputed data was generated by each software following the instructions with default parameters. For algorithms required number of clusters as input (e.g., Bfimpute and AutoClass), the number of labeled cell types were used. As a baseline for comparison, Seurat(58) log-normalized data without imputation was used.

**Table 2.** Methods used in this study.

| Class of method | Methods | Platform | DOI and Reference | Post-logarithm | Further Evaluation* |
| --- | --- | --- | --- | --- | --- |
| Baseline: no imputation<br>(labeled as raw) | **Log-normalization (Seurat)** | R | https://doi.org/10.1016/j.cell.2019.05.031 (58) | No | Yes |
| Matrix decomposition /<br>factorization | **afMF** | Python | Developed in this study | No | Yes |
| Matrix decomposition /<br>factorization | **ALRA** | R | https://doi.org/10.1038/s41467-021-27729-z (4) | No | Yes |
| Matrix decomposition /<br>factorization | **Bfimpute** | R | https://doi.org/10.1016/j.crmeth.2021.100133 (44) | No | Yes |
| Matrix decomposition /<br>factorization | **scRMD (raw and log)** | R | https://doi.org/10.1093/bioinformatics/btaa139 (56) | Yes/No | Yes |
| Matrix decomposition /<br>factorization | SMURF | Python | https://doi.org/10.1093/bib/bbad026 (57) | Yes | No |
| Model and smoothing based | ADimpute | R | https://doi.org/10.1371/journal.pcbi.1009849 (52) | No | No |
| Model and smoothing based | **ccImpute** | R | https://doi.org/10.1186/s12859-022-04814-8 (46) | No | Yes |
| Model and smoothing based | **I-impute** | Python | https://doi.org/10.1186/s12864-020-07007-w (49) | Yes | Yes |
| Model and smoothing based | **knn-smoothing (KS)** | Python | https://doi.org/10.1101/217737 (51) | Yes | Yes |
| Model and smoothing based | **MAGIC (square root and log)** | Python | https://doi.org/10.1016/j.cell.2018.05.061 (3) | No | Yes |
| Model and smoothing based | RESCUE | R | https://doi.org/10.1186/s12859-019-2977-0 (53) | No | No |
| Deep learning based | **AutoClass** | Python | https://doi.org/10.1038/s41467-022-29576-y (5) | No | Yes |
| Deep learning based | scGNN2 | Python | https://doi.org/10.1093/bioinformatics/btac684 (54) | No | No |
| Deep learning based | scIGANs | Python | https://doi.org/10.1093/nar/gkaa506 (55) | Yes | No |
| Deep learning based | **DCA** | Python | https://doi.org/10.1038/s41467-018-07931-2 (8) | No | Yes |
| Deep learning based | scVI | Python | https://doi.org/10.1038/s41592-018-0229-2 (47) | Yes | No |
| Deep learning based | deepimpute | Python | https://doi.org/10.1186/s13059-019-1837-6 (48) | Yes | No |
| Deep learning based | GNNimpute | Python | https://doi.org/10.1186/s12859-021-04493-x (45) | Yes | No |

Nineteen methods including unimputed log-normalization (Seurat) and eighteen imputation algorithms based on R or Python were evaluated in this study.
\* Ten algorithms showed in Bold Font, which passed the pre-screening test and were selected for further in-depth evaluations.

### Data pre-processing

Processed datasets were collected from previous studies. For raw datasets without quality control, we applied the following filtering criteria to remove: (1) cells expressed < 200 genes; (2) outlier cells based on number of genes expressed (nFeature_RNA), number of total counts (nCount_RNA), and percentage of mitochondrial gene expression (percent.mt); (3) Genes expressed in less than 10 cells. For raw data as comparison, Seurat log-normalization was performed. All the imputed data was log-transformed (i.e., either before or after imputing, depending on the software). Simulated datasets were generated using the parameters shown in **Table 1**.

### New imputation algorithm using Low-Rank Full Matrix Factorization: afMF and its implementation

afMF (**a**daptive **f**ull **M**atrix **F**actorization) is an imputation algorithm that builds upon the methodology of another algorithm called ALRA. The algorithm begins by following ALRA’s data preprocessing steps, which include normalization, log-transformation, and rank selection. While ALRA employs randomized SVD for imputation, afMF takes a different approach by utilizing full matrix factorization as the imputation method. Once the rank (k) has been chosen, afMF assumes that the target single-cell gene matrix (X) consists of two matrices (P and Q). The first matrix contains the latent information of genes, while the second matrix contains the relative cell trait information of each cell with respect to the hidden gene information.

To solve the two matrices, afMF employs an iterative gradient descent algorithm. The algorithm fixes one matrix and solves for the optimal values of the other matrix based on its first-order derivatives. This iterative process, which is the key feature of this algorithm, allows afMF to refine the matrices and progressively improve the imputation accuracy for the task of matrix completion. Overall, afMF presents an adaptive approach to scRNA-seq imputation, combining the preprocessing steps of ALRA with its own full matrix factorization method. By leveraging iterative gradient descent, afMF iteratively updates the matrices to effectively reconstruct the missing values and enhance the quality of the imputed single-cell gene expression data. The iteration will be completed when the difference in norm between imputed matrix at time t and t-1 is smaller than 10^−4^.

After the iteration, the entire single cell gene expression matrix is reconstructed by matrix multiplication. Subsequently, it applies ALRA’s approach to rescale the completed matrix product. As definition of biological zero is not clear-cut, only outliers lower than 3 sigma were assigned as biological zero.

The algorithm is available at GitHub: https://github.com/GO3295/SCImputation

The details are as follows:

**<u>Regularization (alpha = 0)</u>**

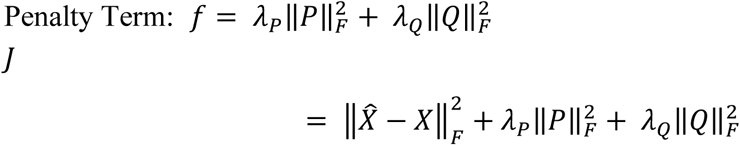

Considering *PQ*^*T*^ = *X̂*

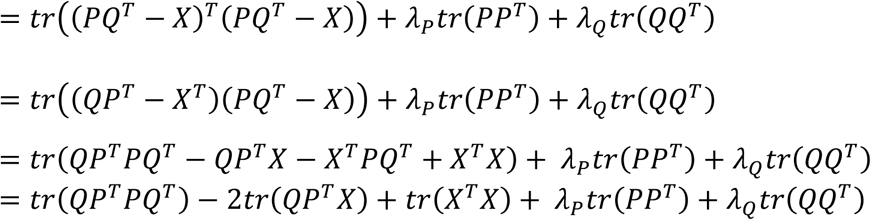

Initializing *P*(^0^), *Q*(^0^) as random matrix

For *P*^(*i*)^

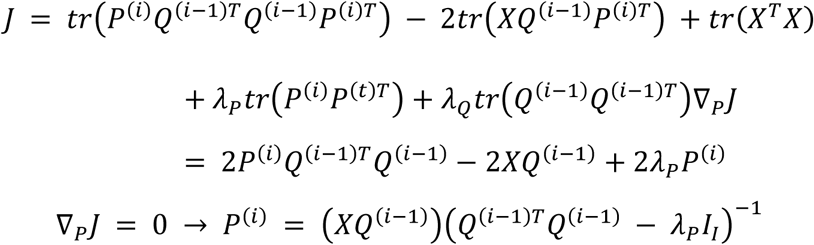

For *Q*^(*i*)^

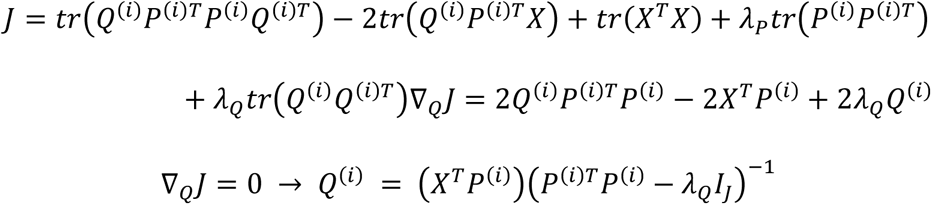

### Selection of other short-listed imputation algorithms for detailed benchmarking with afMF

Good imputation algorithms should have good robustness to any types of datasets. A pre-screening test including two datasets (GSE75748 and GSE81861) to select algorithms for further evaluations were performed. The pre-screening evaluations included: (1) Differential Expression (DE) Analysis using Rank Sum Test in pairwise cell type comparisons; (2) Biomarker Prediction (Area Under Curve) and Classification (Random Forest); (3) Clustering (Louvain Algorithms); (4) Single Cell-Bulk Profiling Similarity. Details were described below. Based on the results (**Additional file 1, Figure S1**), eleven methods were used for further in-depth evaluations (**Table 2**).

### Downstream applications

#### Gene Expression Violin plots and Cell-Cell Correlation visualizations

Initially, we explored how the gene expression visualization changed when using various imputation methods. We used GSE155673 and plotted GAPDH, CD8A and CD19 for comparing both housekeeping gene and cell type specific markers across different cell types. The Seurat function VlnPlot() was applied.

We next calculated the cell-cell correlations in CellBench-10X5CL to study if imputation can increase cell-cell similarity within same cell types and improve visualizations. Spearman correlation coefficients were calculated between pairwise cells and results were visualized through heatmaps using R pheatmap function.

#### Differential Expression (DE) Analysis

We evaluated the performance of different imputations methods on DE analysis using seven real scRNA-seq datasets with their matched bulk (GSE75748 (single cell and time-course), GSE81861, CellBench-10X5CL, GSE124872, EGAS00001003215, GSE79578 and GSE90047), where the matched bulk DE results was used as ‘gold standard’. Datasets were divided into three types: mixture cell types, purified cell types with two conditions, and time-course. The gene expression between pairwise cell types, conditions and time were compared respectively. In brief, bulk DE analysis was performed by limma-voom(59). DE analysis for scRNA-seq was performed by Wilcoxon Rank Sum test, MAST(13) and pseudobulk-limma-trend(14). The popular pseudobulk DE analysis aggregates the information from same cell types and individuals and has been demonstrated to be the preferred method for DE analysis(14,60). Note that the pseudobulk DE analysis was only performed on datasets with multiple individuals (GSE124872 and EGAS00001003215). Results were then ranked by sign (i.e., the sign of the fold change) -log_10_(P-value) and the absolute values of fold changes (FC). Next, the Spearman correlation coefficients were calculated for both gene ranks and logFC between the bulk results and the single cell results. All genes and the top 1000 DEGs (in bulk) were compared. Additionally, the significances of the top 500 DEGs and tailed-20%-percentiles non-DEGs in bulk data were compared among different methods. The false positive rates, defined as the percentages of tailed 10%, 20%, 25%, 33%, and 50% percentiles genes in bulk presented in the top 500 DEGs in single cell results, were also compared. These metric values were subtracted by the results of the unimputed log-normalized data. Extreme values were limited to a cutoff value for better visualization. All the analyses were performed with the default parameters unless other specified. Note that in time-course data the median values of the three datasets were used for bar plots since only three data points presented.

We additionally performed the DE analysis in a real application (GSE155673), where we used pseudobulk-limma-trend to study the interferon (IFN) response between COVID-19 patients and normal controls. Five IFN genes IFI27, IFI44L, IFIT1, IFIT2, IFITM3 and four housekeeping genes ACTB, UBC, GAPDH and SDHA were selected and their P-values and logFC were compared among different imputations. Next, we collected all IFN-related genes from IFN-related Gene Ontology (GO) terms and calculated the percentages of genes that reached statistical significance under different cutoffs (P.adjusted<0.05/0.1).

#### Gene Set Enrichment Analysis (GSEA)

The GSEA was performed for each DE result using the R package ClusterProfiler(16), based on either sign -log_10_(P-value) or logFC (i.e., as input). The Gene Ontology (GO) terms were used as the gene sets. The bulk GSEA results were used as the gold standard. The Spearman correlation coefficients were calculated for ‘sign (Enrichment Score) × -log_10_(P-value)’ between the bulk GSEA results and the single cell GSEA results. All terms and ‘P<0.05’ terms were compared. The metric values were subtracted by the results of the unimputed log-normalized data. Extreme values were limited to a cutoff value for better visualization. All the analyses were performed with the default parameters unless other specified. Note that in time-course data the median values of the three datasets were used for bar plots since only three data points presented.

#### Classification and Biomarker Predictive Performance

Classifications for different cell types, conditions and time point in mixture, purified and time-course data were performed using Random Forest (RF) model from R RandomForest package. The top 10 DEGs for each comparison (i.e., one vs. others) in bulk data were used as input features. For each dataset, 70% samples were used as training set and the other 30% were for testing. The classification accuracy and RF predict probability for true label were compared.

The biomarker predictive performances of different methods were evaluated. Top 10 DEGs for pairwise cell types/conditions comparisons in bulk data were used as biomarkers for evaluation. The predictive performance was evaluated by calculating the area under the receiver operating characteristic curve (AUROC) for each of the selected genes in corresponding comparisons. The metric values were subtracted by the results of the unimputed log-normalized data. All the analyses were performed with the default parameters unless other specified.

#### Automatic cell type annotation

Two well-established R packages, SCINA(17) and ScType(18), were applied to evaluate the performance of different imputations on automatic cell type annotation. Two well-labeled data (GSE155673 and ROSMAP brain) were used. The cell-type-specific marker genes used as input were collected from Azimuth (https://azimuth.hubmapconsortium.org/) and PanglaoDB(61). The annotation accuracy, F1 score, true-label-annotation probability and unknown rate were calculated and compared. Extreme values have been limited to a cutoff value for better visualization. All the analyses were performed with the default parameters unless other specified.

#### Dimension Reduction, Clustering, and Cell Cycle Dynamics

To quantitatively evaluate the clustering performance of different methods, we adopted four evaluation metrics, Entropy of accuracy (*H*_*acc*_), Entropy of purity (*H*_*pur*_), Adjusted Rand Index (ARI) and Normalized mutual information (NMI) as previously suggested(9,10) using K-means Clustering and Louvain Clustering algorithms. Three simply labelled (i.e., with ground truth cell type label) datasets GSE75748, GSE81861, and CellBench 10X 5CL were used. Note that we did not include datasets with complicated design (i.e., with multiple conditions, tissues, cell types, batches, diseases, individuals, etc.) as there will be multiple labels for testing. In brief, the number of clusters was set to the number of known cell type labels in each dataset. The Seurat FindVariableFeatures() function with ‘vst’ methods were used to select high variable genes (top 3000 genes), followed by running PCA. The top 10 PCs were used for the two clustering algorithms. The detailed implementation of the algorithms was described previously(9).

*H*_*acc*_ evaluates the difference of the true groups within each predicted cluster, defined as:

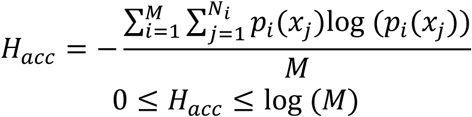

where *M* is the number of predicted clusters; *N*_*i*_ is the number of true groups in the *i*^*th*^ predicted cluster; *x*_*j*_ are cells in the *j*^*th*^ true group; and *p*_*i*_ (*x*_*j*_) are the proportions of cells in the *j*^*th*^ true group relative to the total number of cells in the *i*^*th*^ predicted cluster.

*H*_*pur*_ evaluates the difference of the predicted clusters within each true group, defined as:

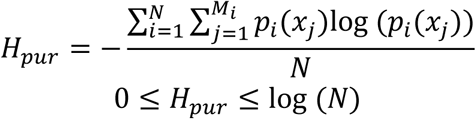

where *N* is the number of true group; *M*_*i*_ is the number of predicted clusters in the *i*^*th*^ true group; *x*_*j*_ are cells in the *j*^*th*^ predicted cluster; and *p*_*i*_ (*x*_*j*_) are the proportions of cells in the *j*^*th*^ predicted cluster relative to the total number of cells in the *i*^*th*^ true group. Smaller *H*_*acc*_ and *H*_*pur*_ indicates better clustering performance(9).

Adjusted Rand index (ARI) that evaluates the similarities between two data distributions [cite benchmark] was calculated using adjustedRandIndex() function in mclust package(62), defined as:

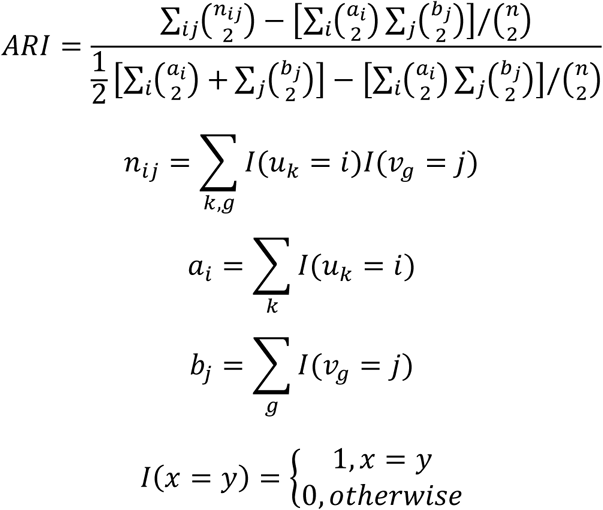

Where *i* and *j* enumerate the *k* clusters; 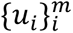 is the predicted cluster label; 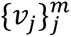 is the true group label.

Normalized mutual information (NMI) that measures the correlation between two random variables(10) is calculated using NMI() function from aricode package, defined as:

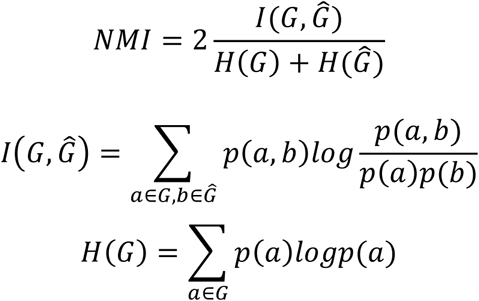

Where *G* is the true group, *Ĝ* is the predicted cluster. *p*(*a*), *p*(*b*) and *p*(*a*, *b*) are the probabilities that the cell belongs to cluster a, cluster b and both, respectively. For both ARI and NMI, higher values indicate better clustering performance.

For cell cycle dynamics, the dataset GSE64016 (with known cell cycle label G1, S, and G2M) and the Seurat function CellCycleScoring() was applied. Briefly, the cell cycle status and score for each cell was predicted by the Seurat algorithm. To compare the performance of the predictions in each imputation, Wilcoxon rank sum tests were performed on the predicted cell cycle scores between different known cell cycles (i.e., S vs. Others, G2M vs. Others) and the p-values were compared. The prediction accuracy (predicted cell cycle vs. true cell cycle) and the F1 score were calculated for each imputation.

To visualize the cell type/cell cycle clustering, PCA was performed followed by running UMAP with first 50 PCs. The two-dimension UMAP plots were generated for clusters/cell types/cell cycles.

To compare the clustering metrics among different methods, we first subtracted all the ARI and NMI values by the results of the unimputed data (vice versa for *H*_*acc*_ and *H*_*pur*_ for better visualization) and then took the median values of the three datasets. Extreme values were limited to a cutoff value for better visualization in plots. UMAP plots were generated for CellBench 10X 5CL and GSE64016 to evaluate the performance of imputations on dimension reductions. All the analyses were performed with the default parameters unless other specified.

#### Pseudotime trajectory analysis

The impact of imputation on pseudotime trajectory analysis was evaluated by Monocle3(19), DPT(21) and Slingshot(20) on eight datasets CellBench cellmix1-4 (four datasets with two lineages), GSE118068 (embryonic and postnatal), GSE75748 time-course, GSE79578 and GSE90047 that with real time/lineage labels.

In Monocle3, the starting point of the trajectory was set based on known information and other parameters were set to default. Monocle3-normalization was also applied for comparison. Next, Spearman correlation coefficients were calculated between the predicted pseudotime and the real lineage/time labels. The Pseudo-temporal ordering score (POS) which measures cell orders were calculated using orderscore() in TSCAN package, defined as:

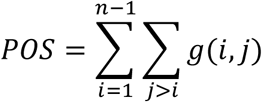

Where n is the number of cells and *g*(*i*, *j*) is the score that evaluates how well the predicted order of the i^th^ and j^th^ cells in the ordered path matches their known order(10). 2-D PCA and UMAP plots were generated for visualizing trajectories.

In DPT, the starting point and end point were set as ‘tips’ based on known information with other parameters set to default. Next, Spearman correlation coefficients and POS were calculated between the predicted DPT values and the real lineage/time labels. Note that due to the limitations of the software (i.e., always result in two branches), only CellBench data with two branches were used in DPT. The predicted branches by DPT were further evaluated with clustering metrics *H*_*acc*_, *H*_*pur*_ ARI, and NMI (as described in Clustering). 2-D diffusion map plots were generated for visualizing trajectories.

In Slingshot, the start and end point of the trajectory was set based on known information and other parameters were set to default. Next, Spearman correlation coefficients and POS were calculated between the predicted pseudotime and the real lineage/time labels. For trajectory with multiple branches, the predicted branches and the real branches were matched by their largest overlaps of cells. For instance, for datasets of three cell types (A, B, and C where A will differentiate to B and C) with even number, ideally predicted branch 1 would have 100% overlap rate with A-to-B known branch while only have 50% overlap rate with A-to-C known branch (vice versa for predicted branch 2). Then the true overlap rates (i.e., largest overlaps between predicted branches and real branches, as matched branches) were calculated and compared. 2-D PCA plots were generated for visualizing Slingshot results. Slingshot-normalization was applied for comparison as well.

The metric values were subtracted by the results of the unimputed log-normalized data. Extreme values were limited to a cutoff value for better visualization.

#### AUCell for pathway activation and SCENIC for gene regulatory analysis

We evaluated the AUCell application among different imputations on GSE155673, where the activations of the interferon (IFN) related pathways between COVID-19 and healthy controls were compared within monocytes. Briefly, the IFN-related pathways were collected from both Hallmark(15) gene sets (2 terms) and GO(63) gene sets (31 terms). Each cell was then assigned a binary status (activated or not) for each pathway by the AUCell pipeline and the percentages of monocytes with activated pathway were compared between COVID-19 and healthy controls. All the analyses were performed with the default parameters unless other specified.

We next investigated the compatibility of SCENIC application with different imputations. We followed the pipeline as previously suggested(64) using the same dataset GSE115978 and the various normalized/imputed data were used as input. Regulons with Z-score≥3 for each of the method were collected and visualized through heatmap and Venn diagram for comparison. Next, seven well-established regulons (i.e., TCF7, EOMES, TBX21, MITF, MYC, MAFB, PAX5) as previously demonstrated(64) were picked out. AUCell (implemented in SCENIC) were used to determine whether the selected regulon was activated in a selected cell. Ideally, a cell-type-specific regulon was expected to be activated only in the specific cell type, but not the other cell types. The percentages of cells that with activated regulons within expected cell types and other cell types were calculated and compared. The metric values were subtracted by the results of the unimputed log-normalized data. Extreme values were limited to a cutoff value for better visualization. All the analyses were performed with the default parameters unless other specified.

#### CellPhoneDB and CellChat for Cell-Cell Communication

We evaluated the potential impacts of imputation on cell-cell communications using the two packages CellPhoneDB(23) and CellChat(24) on datasets E-MTAB-6701 and E-MTAB-13384. For CellPhoneDB, we collected the interaction pairs of ligand-receptors-cell type as used in previous protocol(23) and followed the pipeline with all the parameters set to default. Bubble plots showing active interactions with adjusted p-values and mean expressions were generated for comparisons. The P-values of top interactions (in raw) and the number of significant interactions under different significant levels were compared. In CellChat analysis, we aimed to evaluate the communications from monocytes/macrophages to fibroblasts or other cell types as demonstrated in previous paper(30). Note that in the original research this communication was not replicated in this mice dataset and therefore we aim to verify if imputation could make validations. We followed the pipeline with all the parameters set to default and normalized/imputed data as input. The numbers/percentages of significant interactions (all or from-monocyte, stratified by CHIP status) were compared. Additionally, network plots for source-Monocyte and the bubble plots for selected interaction pairs from monocytes to fibroblasts(30) were generated for comparisons.

### Supporting Analysis

*Correlations between bulk RNA-seq profiles and single cell RNA-seq profiles* In this part, we adopted Hou’s method(9), made some revisions and extended to different data (GSE75748, GSE81861 and CellBench-10X5CL).

Specifically, Spearman correlation coefficients were calculated for each single cell profile and the bulk profile (within same group):

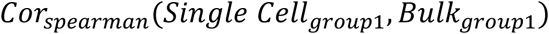

We noticed some methods can equally improve the correlations of different groups. Therefore, to reduce this influence we subtracted the results by the max different-group correlation coefficients:

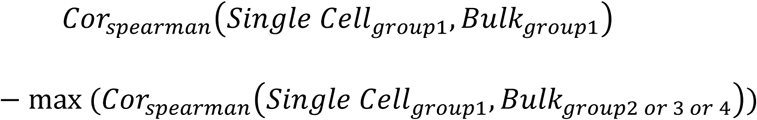

The median value of each result was calculated for comparison.

We performed the same analysis on pseudobulk profiles. Spearman correlation coefficients were calculated for each pseudobulk profile (i.e., median-aggregate of the same group) and the bulk profile (same group):

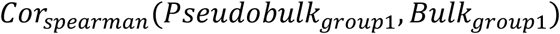

Then subtracted the results by the max different-group correlation coefficients:

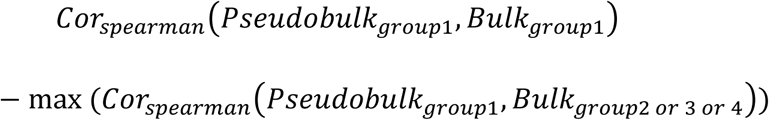

Next, we conducted pair-wise calculations of log fold changes between different groups in both pseudobulk profiles and bulk profiles and performed the Spearman correlation analysis:

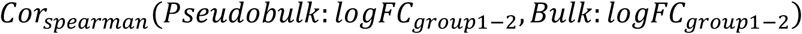

The metric values were subtracted by the results of the unimputed log-normalized data. Extreme values were limited to a cutoff value for better visualization.

#### Correlations between surface protein profiles and mRNA profiles in single cell

We used the data GSE100866 to study the impact of imputation on the correlations between measured mRNA and surface protein(4). Six marker genes in two tissues (CD4, CD2, CD19, CD14, CD34, CCR7 in PBMC and CBMC) with corresponding surface protein measurement were selected and compared. Spearman correlation coefficients were calculated between the two measurements for each gene. The metric values were subtracted by the results of the unimputed log-normalized data.

### Simulated data analysis

The simulated datasets with ground truth were generated using Splatter(42) (Mock90) and SplatPop(43) (SplatPop90) with parameters shown in **Table 1**. The evaluations for simulated data included Differential Expression Analysis, Classification, Biomarker Prediction, Automatic Cell Type Annotation, Dimension reduction & Clustering, and Imputed SC-Ground Truth Similarity as described above. Note that the ground truth (e.g., known DEGs, groups, true matrix) instead of bulk data were used as gold standard in these evaluations.

### Running time, memory usage and recommendation

We used four datasets (10000×1500, 10000×5000, 10000×10000 and 10000×50000 matrix) to evaluate the time spent and memory usage of different imputation methods. The time spent and memory usage were plotted in log scale.

A summary heatmap was generated based on the evaluations. The performances were classified into five levels ‘Generally Worse’, ‘Slightly Worse’, ‘No Obvious Difference’, ‘Slightly Better’ and ‘Generally Better’ and were highlighted in different colors. Note that algorithm-evaluation were labelled as ‘No Obvious Difference’ if that algorithm had both obvious advantages and disadvantages based on different metrics in that evaluation. 1-8 cores were used depending on datasets and algorithms, and the maximum memory usage for afMF (i.e., 50,000 cells) is about 22.3 GB.

### Other statistical analysis

The coding work including imputation algorithms and evaluations was performed using R 4.2.1 or Python 3.8. For each of the evaluations with enough data, Wilcoxon Rank Sum test was performed between the results of the imputation algorithms and the unimputed log-normalization. The significances were indicated with ‘*’ or ‘.’, where one, two and three ‘*’ represented ‘P<0.05’, ‘P<0.01’ and ‘P<0.005’ respectively and ‘.’ represented ‘P<0.1’. Note that some imputation algorithms were not assessed in some of the evaluations due to the processing failures/errors, large memory usage or long running time.

## Results

Based on the pre-screening results (**Additional file 1, Figure S1**), ten algorithms with stable performance (i.e., generally not worse than no-imputation) were used for further in-depth evaluations (**Table 2**).

### Imputation improved on the dropout problem and basic analysis: restoration of HK gene expression violin plots, 2-D PCA, and cell-cell correlation

We firstly focused on the impact of prior data imputations on the dropout problem and basic data analysis. We did a survey to show the extent of dropout affecting the two kinds of genes: housekeeping genes and cell type specific genes (**Figure 2 and Additional file 2, Figure S2-4**). In fact, they represented technical zeros here as the expected counts of these highly expressed genes should not be zero in the single cell samples and these were regarded as false zeros. We observed a heavy zero tail for B cells, CD4+ T cells, and NK cells in the violin plots of log count of GAPDH; in contrast, after afMF, ALRA and DCA imputation, these dropouts were largely ameliorated (**Figure 2A, GAPDH**). For cell type specific marker genes CD8A and CD19, afMF and DCA provided decent distributions in which the marker genes were uniquely higher expressed in CD8+ T cells and B cells respectively with no heavy tail toward zero (**Figure 2A, CD8A and CD19**). ALRA worked to avoid the majority of dropouts in B cells but not for CD8+ T cells. Compared to the other methods, DCA is likely to overfit and distort the data especially for cell type specific marker genes. Results for other algorithms were shown in **Additional file 2 Figure S2-4.**

**Figure 2.**
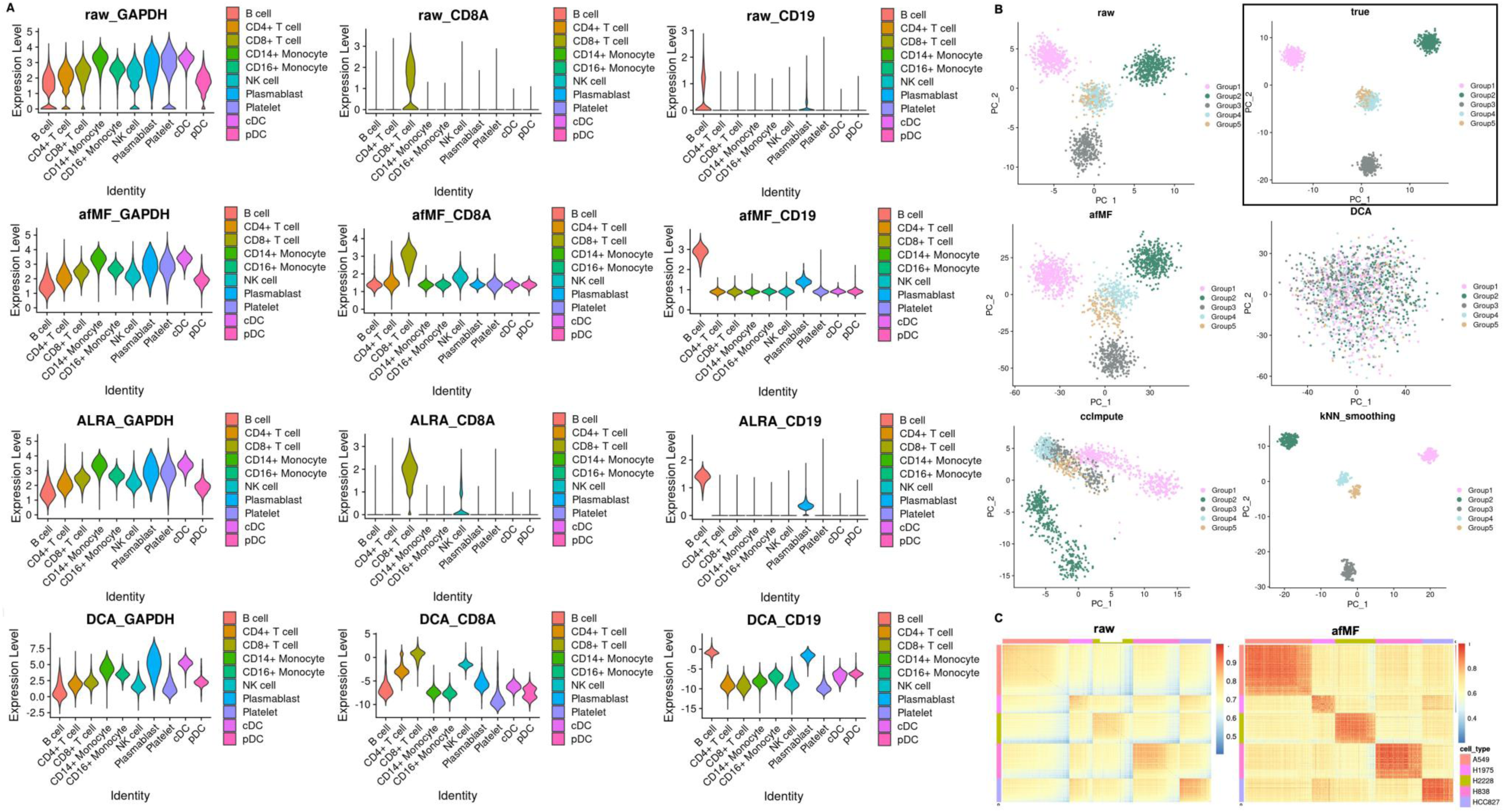
The impact of imputations on the dropouts: gene expression violin plots, 2-D PCA, and cell-cell correlation heatmaps. The impact of imputations on the dropouts and related visualizations. (**A**) The impact of different imputation algorithms on gene expression were visualized through violin plots across different cell types in dataset GSE155673 using housekeeping gene GAPDH and cell type specific gene CD8A and CD19. (**B**) The impact of different imputation algorithms on 2-D PCA were visualized using simulated dataset Mock90 (dropout rate=90%). Results from the raw (not imputed) and the true (ground truth) data were also shown. (**C**) The impact of different imputation algorithms on cell-cell correlations were visualized through heatmaps by calculating the Spearman correlation coefficients (SCCs) between pairwise cells in dataset CellBench-10X5CL.

Next, we examined the potential consequence of imputation in generating artefactual gene co-expression or structure in covariance matrix. A simulated dataset Mock90 without co-expression structure was used to detect any potential fabrication of structure in the eigenvector space after imputations, with visualizations of the first 2 PCs in PCA (**Figure 2B** and **Additional file 2, Figure S5-6**). Compared to the ground truth (true in **Figure 2B**), unimputed data (raw), afMF and ALRA, Bfimpute, kNN-smoothing, and scRMD (shown in **Additional file 2, Figure S5**) could separate Group4 and Group5 without fabricating additional false structure in PCA. However, three Model-based imputation algorithms, ccImpute, I-impute and MAGIC fabricated fortuitous co-expression structure on the PCA embeddings. On the other hand, both Bfimpute and kNN-smoothing provided the best separation of the two groups and concentrated clusters and even outperformed the ground truth. This is likely to be the overfitting problem, and the variances of the same cell types in the 2-D PCA plots were made extremely low which was an artefact of kNN-smoothing imputation and was not present in the ground truth. This is again demonstrated in subsequent real data analysis in which the cells showed abnormal clustering patterns in kNN-smoothing imputation. Fabrication features were also seen in a dataset generated by SplatPop (**Additional file 2, Figure S6**). False co-expression structures were fabricated by AutoClass (Deep learning based) and Bfimpute (Matrix decomposition), while afMF and ALRA better replicated the patterns in ground truth. Combining results of these two simulated datasets, only afMF, ALRA, scRMD (three Matrix-based) and DCA (Deep learning based) did not fabricate artefactual gene co-expression structure in PCA. Those artefacts adversely affected downstream analyses as we are going to show in subsequent sections.

We next visualized cell-cell correlations through heatmaps (**Figure 2C** and **Additional file 2, Figure S7**). afMF, MAGIC/MAGIC-log, ALRA, AutoClass, DCA and kNN-smoothing showed better separations and patterns for different cell types and provided much clearer visualizations. Specifically, similarities between cells within same cell types were enhanced, or cell similarities between different cell types decreased in these imputations, i.e., zeros were no longer zeros and became totally different values in different cell types.

### Imputation improved Differential Expression (DE) Analysis and Gene Set Enrichment Analysis (GSEA) with compatible statistical metrics

DE analysis is the essential part of downstream applications. For scRNA-seq data, DE can be analyzed by one of these methods: (1) MAST; (2) Wilcoxon rank sum test; or (3) pseudobulk analysis. Our results showed that performance of imputation in DE analysis was dependent on the methods used. Enhancement was found in (1) MAST and (2) Wilcoxon rank sum test, while setback was found when using imputed data with (3) pseudobulk analysis for DE analysis and GSEA.

Majority of the previous evaluations used p-value cutoffs, overlap rates, or simulated data that may have drawbacks: heavily affected by sample size, did not make full use of gene rank information, or unable to reflect the real data distribution. One example is that an algorithm may increase overlap DEGs, but the top rankings may be totally different from ground truth, which is not desired. Therefore, we conducted the evaluation based on gene rank information. Using MAST and Wilcoxon rank sum test, higher p-value-based rank concordance between bulk and afMF-imputed DE results were observed for all types of data (i.e., mixture/purified/time-course) with statistical significance (**Figure 3A-C** and **Additional file 3, Figure S8-9**). These conclusions still held when limiting the genes to only top 1000 DEGs (determined in bulk data) (**Additional file 3, Figure S10**). The top 500 DEGs showed greater statistical significance in afMF, ALRA, MAGIC and AutoClass (**Figure 3D-F** and **Additional file 3, Figure S8E-F**), while the tail-ranked genes showed no difference in statistical significance compared to no imputation (data not shown). Next, false positive rates were calculated based on ranks in single cell and bulk (see Methods) and only afMF and I_Impute showed generally lower false positive rates in all types of data (**Figure 3G-I** and **Additional file 3, Figure S11**). Additionally, higher logFC Spearman correlations were observed between bulk and afMF, MAGIC, and DCA imputed results for all types of data (**Additional file 3, Figure S12**).

**Figure 3.**
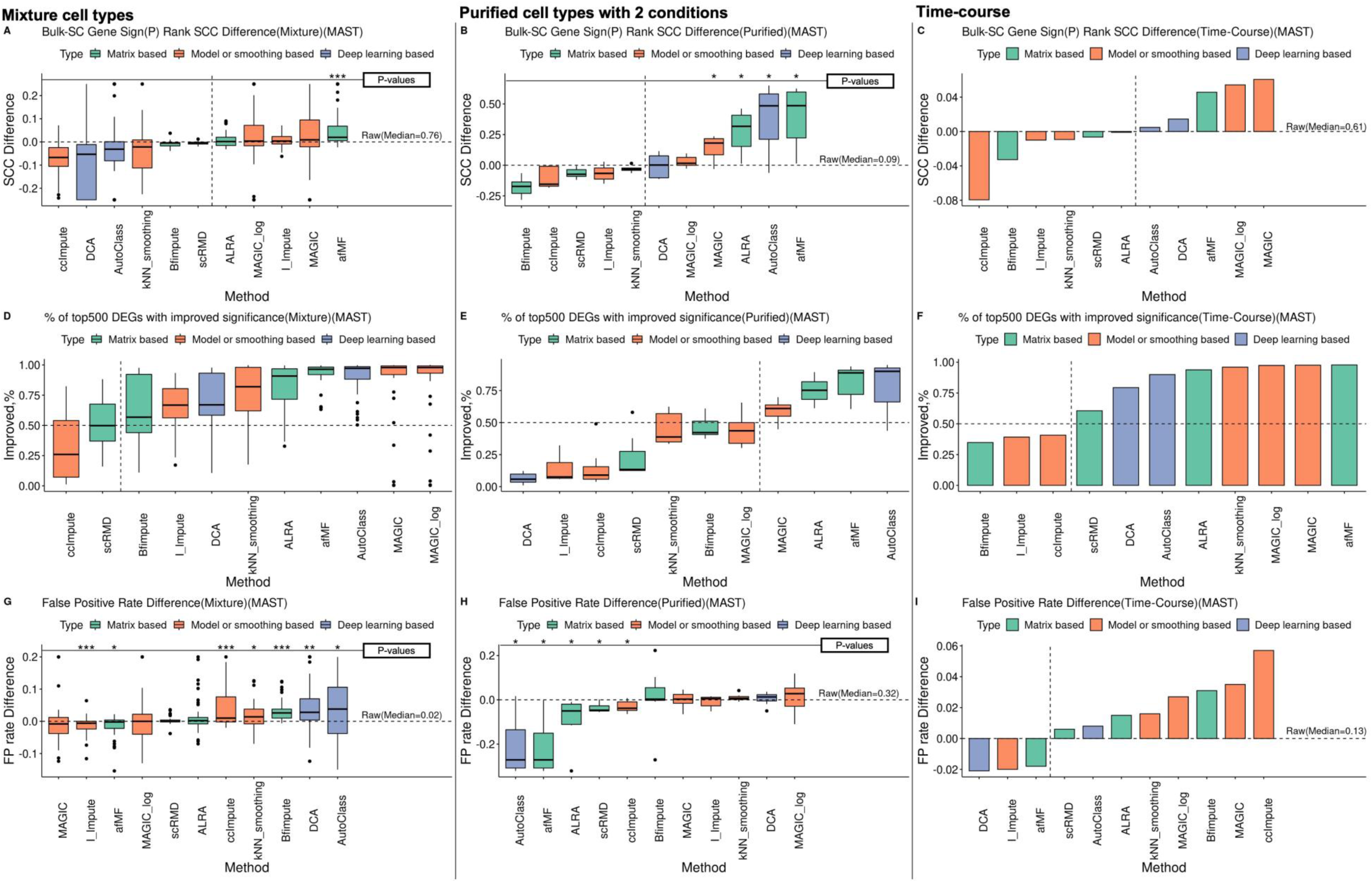
Performance of imputations in differential expression analysis (single cell level with MAST) The performance of different imputation methods on DE analysis were evaluated using seven real scRNA-seq datasets with matched bulk results as ‘gold standard’. Datasets were divided into three types: mixture cell types, purified cell types with conditions, and time-course. The gene expression between pairwise cell types, conditions and time were compared for each type respectively. DE analysis for bulk and scRNA-seq were performed by limma-voom and MAST respectively. Results were ranked by sign (i.e., the sign of the fold change) -log10(P-value) and the absolute values of fold changes (FC). The metrics were subtracted by the values of the raw results. Extreme values have been limited to a cutoff for better visualization. The median values were used for the time-course data since there are only three data points. One, two and three ‘*’ represented ‘P<0.05’, ‘P<0.01’ and ‘P<0.005’ respectively and ‘.’ represented ‘P<0.1’, compared to raw log-normalization using Wilcoxon rank sum test. (**A-C**) The Spearman correlation coefficients (SCC) were calculated for the gene ranks between the bulk results and the single cell results for the three data types. (**D-F**) the significances of the top 500 DEGs (in bulk) were compared among different imputation methods. (**G-I**) The false positive rates (defined as the third tertile genes in bulk results presented in the top 500 DEGs in single cell results) were compared.

Imputation is incompatible with pseudobulk DE analysis using limma-trend. Nearly all the imputation algorithms performed worse than no imputation in pseudobulk DE analysis (**Additional file 3, Figure S13**). Only the logFCs of kNN-smoothing and DCA had higher correlations with bulk results (**Additional file 3, Figure S13B and D**). Though most algorithms increased the significance of the top 500 DEGs, they had higher false positive rates as well (**Additional file 3, Figure S13E-F**). For example, 6 out of 10 imputation algorithms significantly increase the number of false positive DEGs in pseudobulk analysis (**Additional file 3, Figure S13F**). These results suggested that pseudobulk processing may serve as a smoothing step and thus heavily decreased the influence of dropouts in DE analysis.

GSEA is an extension use of DE results to study the enrichments of DEGs in specific biological processes or molecular functions. Whether imputation can improve GSEA was not studied yet. Our results showed that performance with GSEA followed that of DE analysis and it depended on the DE statistics used. Using either MAST DE sign - log_10_P or logFC as input, higher Spearman correlations between bulk GSEA and afMF-imputed GSEA results were observed for all types of data (**Figure 4** and **Additional file 3, Figure S14**). In contrast, MAGIC / MAGIC-log and AutoClass performed generally better as well but showed no improvement when using logFC or sign -log_10_P as input respectively in mixture data. These conclusions held when limiting to enrichment terms with P<0.05 (i.e., in bulk results) (**Additional file 3, Figure S15**). When using Wilcoxon Rank Sum DE results, MAGIC/MAGIC-log, afMF and ALRA showed steady improvements (**Additional file 3, Figure S16**). When using pseudobulk DE results, all imputations with sign -log_10_P as input showed no improvements, while kNN-smoothing, ALRA, I-Impute, DCA and afMF increased the correlations using logFC as input (**Additional file 3, Figure S17**). These results suggested that imputation could improve the discoveries of gene programs under specific conditions.

**Figure 4.**
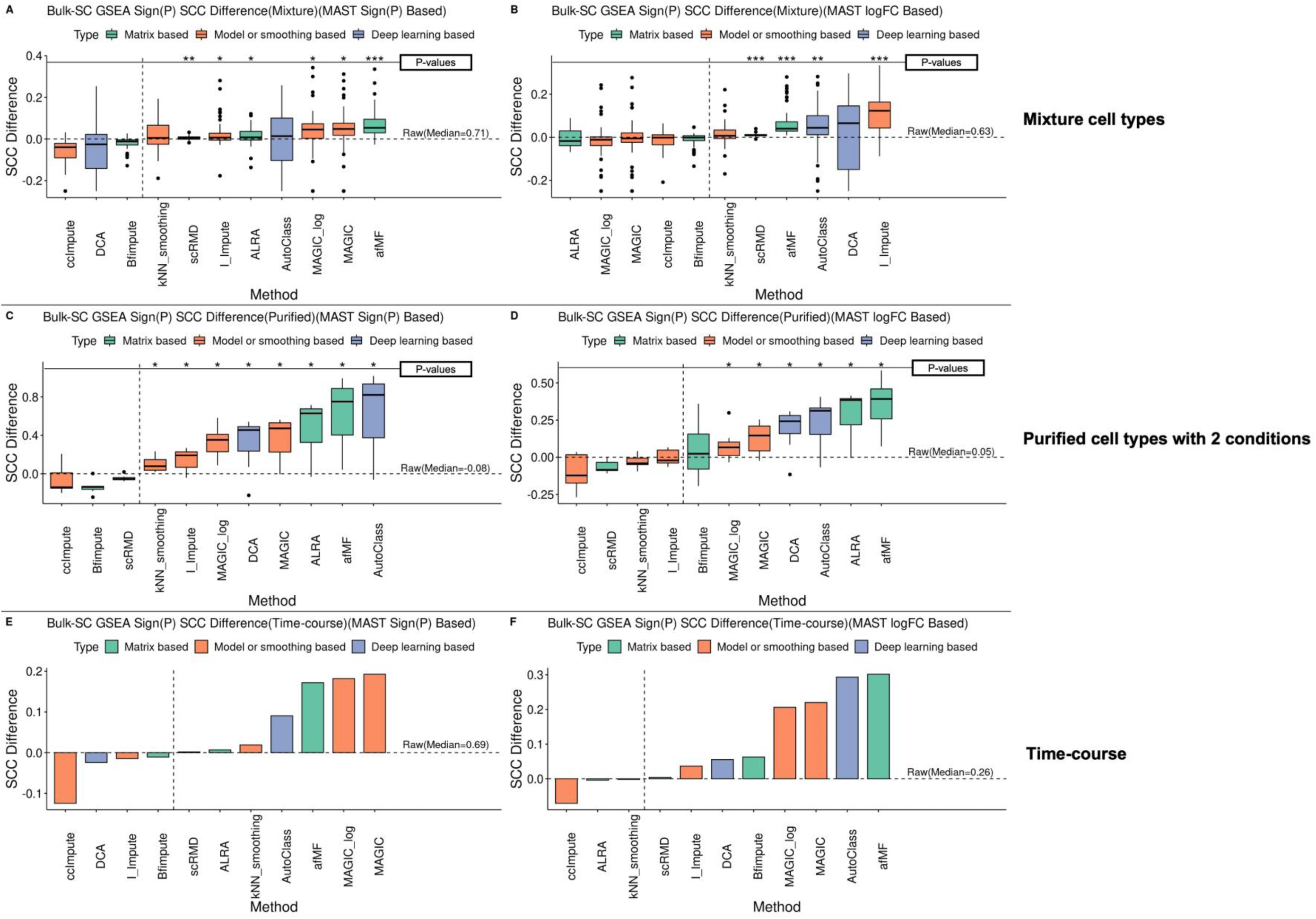
Performance of imputations in GSEA. The GSEA was performed for each of the DE results based on either sign -log10(P-value) or logFC (i.e., ranks as input). The Gene Ontology (GO) terms were used as the gene sets. The bulk GSEA results were used as the gold standard. The Spearman correlation coefficients were calculated for term ‘sign (Enrichment Score) × - log10(P-value)’ between the bulk GSEA results and the single cell GSEA results. All terms were used. The metrics were subtracted by the values of the raw results. Extreme values have been limited to a cutoff value for better visualization. The median values were used for the three time-course datasets. One, two and three ‘*’ represented ‘P<0.05’, ‘P<0.01’ and ‘P<0.005’ respectively and ‘.’ represented ‘P<0.1’, compared to raw log-normalization using Wilcoxon rank sum test. (**A-B**) Mixture Data; (**C-D**) Purified data; (**E-F**) Time-course data.

### Imputation improved Classification, Biomarker Prediction and Automatic Cell Type Annotation

Classification would be useful when identifying cell subtypes or predicting cells with unknown labels. Using Random Forest model, we observed higher classification accuracy in nearly all the imputation methods for all types of data, except for DCA & kNN-smoothing (**Figure 5A-C**). Most imputation methods showed improvements in terms of correct cell type prediction probabilities except I-Impute and Bfimpute (**Figure 5D** and **Additional file 4, Figure S18A-B**).

**Figure 5.**
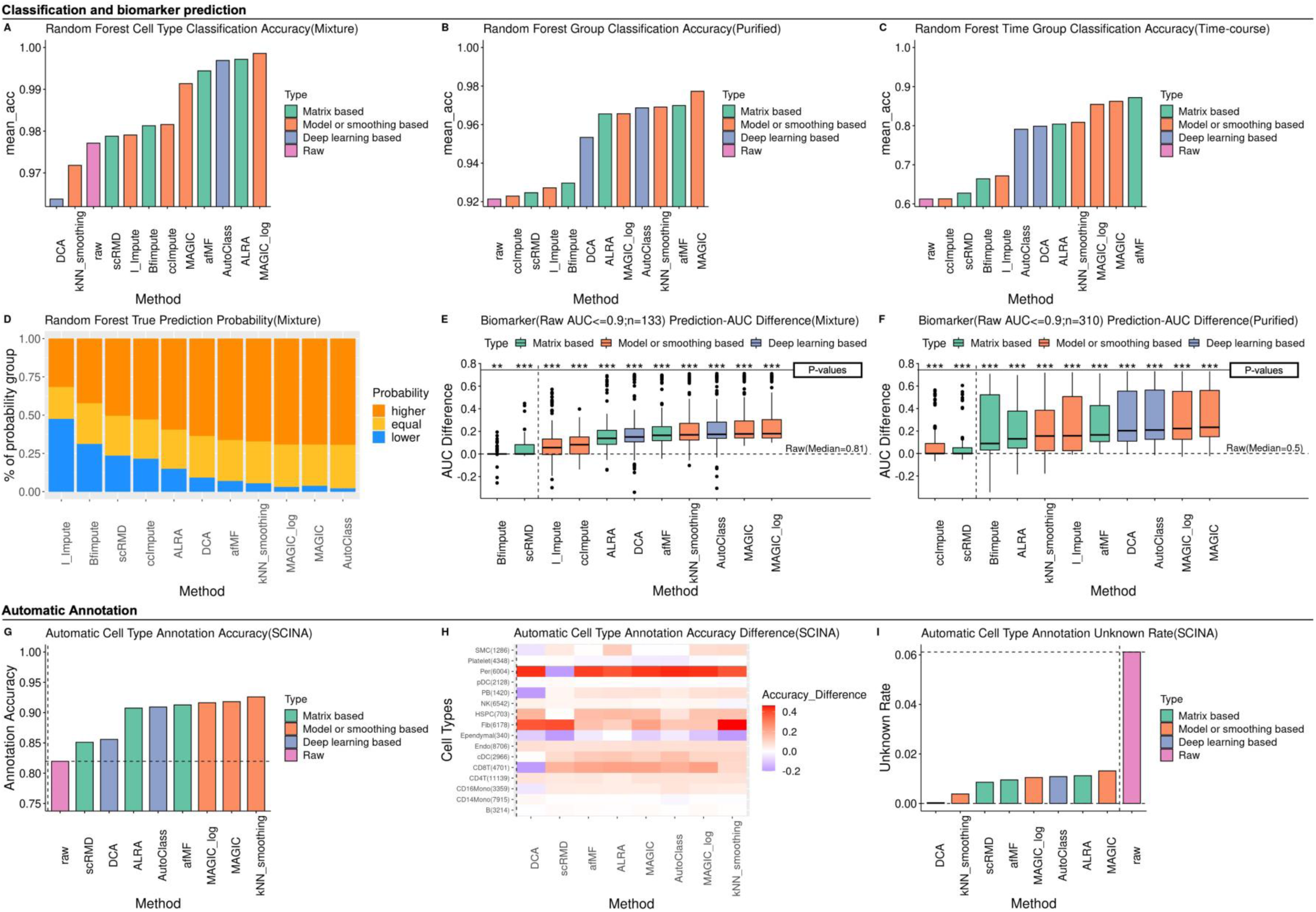
Performance of imputations in Classification, Biomarker Prediction and Automatic Cell Type Annotation. Classifications for different cell types, conditions and time point in mixture, purified and time-course data were performed using Random Forest (RF). The top 10 DEGs for each comparison in bulk data were used as input features. For each dataset, 70% samples were used as training set and the other 30% were for testing. The biomarker predictive performance of different methods was evaluated. Top 10 DEGs between pairwise cell types/conditions in bulk data were used as biomarkers in mixture/purified data. SCINA was applied to evaluate the performance of different imputations on automatic cell type annotation. Two well-labeled data (GSE155673 and ROSMAP brain) were used. The cell-type-specific marker genes used as input were collected from Azimuth and PanglaoDB. Some metrics were subtracted by the values of the raw results. One, two and three ‘*’ represented ‘P<0.05’, ‘P<0.01’ and ‘P<0.005’ respectively and ‘.’ represented ‘P<0.1’, compared to raw log-normalization using Wilcoxon rank sum test. (**A-C**) The classification accuracy for mixture, purified, and time-course datasets, and (**D**) RF prediction probability for true label were calculated. (**E-F**) The predictive performance was evaluated by calculating the AUROC for each of the biomarkers in each comparison. (**G-I**) The annotation accuracy and unknown rate were calculated and compared.

Cell-type-specific marker genes can be used to identify different cell types. Using Area Under Curve (AUC), higher AUC values for biomarker genes to discriminate cell types/conditions in mixture/purified data were discovered for MAGIC/MAGIC-log, AutoClass, kNN-smoothing, afMF, ALRA and DCA (**Figure 5E-F** and **Additional file 4, Figure S18C-F**). We also analyzed the extent of potential false positives and the non-marker genes were selected (based on bulk DE). Much higher percentage of non-markers with AUC>0.9 were discovered in AutoClass, MAGIC/MAGIC-log and DCA for both mixture and purified data (**Additional file 4, Figure S18D** and **F**). Therefore, only kNN-smoothing, afMF, and ALRA enhanced detection of cell type marker genes without increasing the false positive rate, indicating that they are better imputation algorithms for the purpose of identify cell-type-specific markers.

Cell type annotation(65) is the key step in scRNA-seq analysis. The use of imputation for cell type annotation may be underestimated. Since it usually takes much time and effort to do manual annotation, researchers have developed automatic tools to easily annotate cell types, such as SingleR(66), scmap(67) and Azimuth that require reference data, or other tools only require cell type marker genes as input. As imputation showed some advantages in classifications and biomarker predictions, we further investigated whether imputation was compatible with automatic cell type annotations. Using two well-established packages SCINA(17) and ScType(18), higher annotation accuracy, F1 scores and true prediction probabilities were observed for nearly all the selected imputation methods (**Figure 5G-H** and **Additional file 4, Figure S19**), except for DCA that performed worse in some cell types. As a result, all the selected imputation methods resulted in lower unknown rates (**Figure 5I**).

### Imputation moderately improved Clustering, Dimension Reduction, and Project Visualization

Clustering is the essential step for exploring subtypes. In general, various imputation enhanced cell clustering to various extents. Using Louvain and K-means algorithms on three mixture datasets with the Entropy of Accuracy (*H*_*acc*_), Entropy of Purity (*H*_*pur*_), Adjusted Rand Index (ARI) and Normalized Mutual Information (NMI), afMF and MAGIC-log showed improvements across all metrics compared to no-imputation (**Figure 6A-B**). AutoClass and ccImpute performed well only with Louvain algorithms, while MAGIC and ALRA performed better with K-means algorithms. In UMAP visualization for CellBench-10X5CL, afMF, ALRA, Bfimpute, ccImpute and scRMD remained similar and consistent structure of cell clusters, while others (e.g., kNN_smoothing, DCA, I_impute) showed artifacts by producing strange shapes and generated unexpected cell cluster patterns that might lead to false discoveries (**Additional file 5, Figure S20-21**).

**Figure 6.**
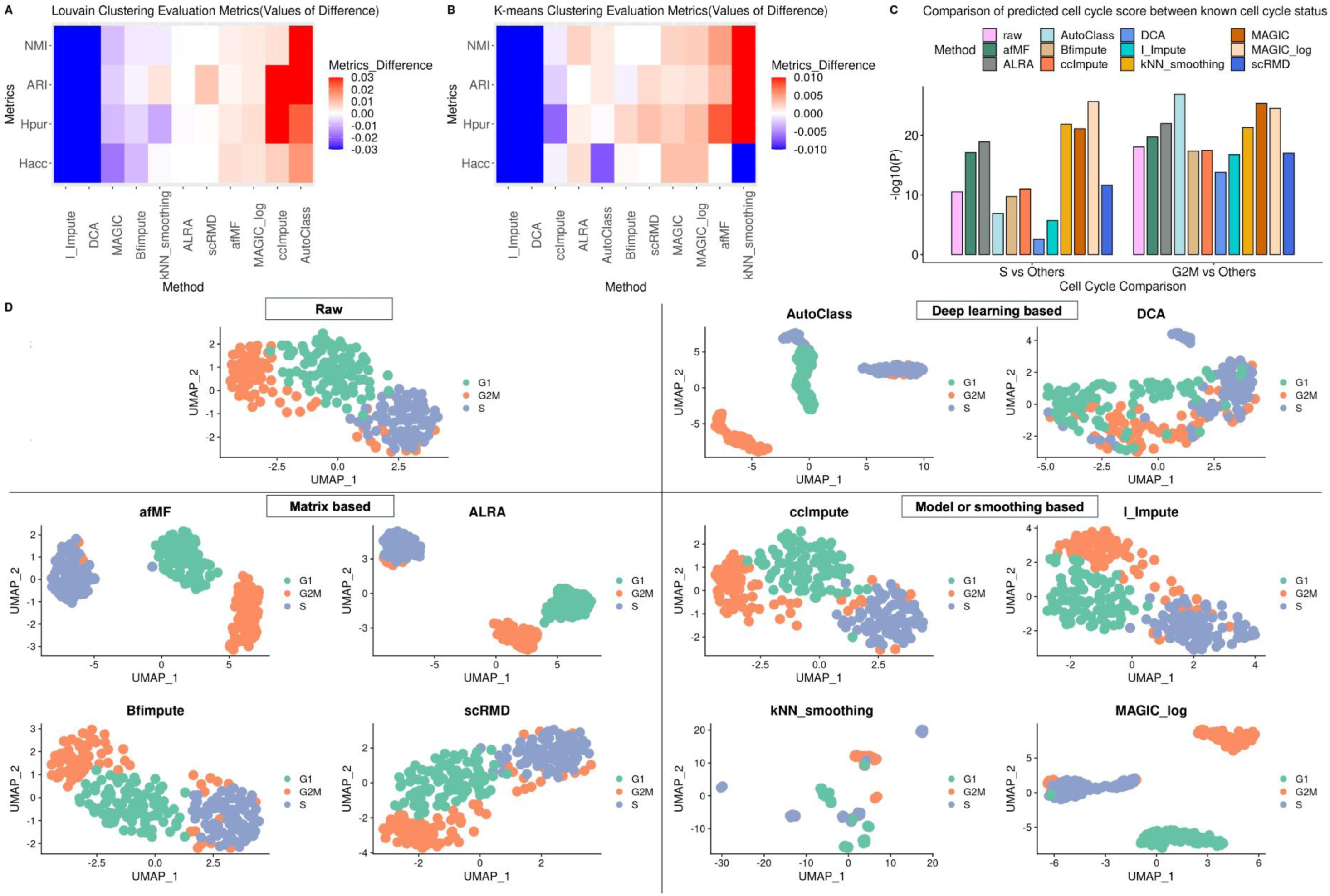
Performance of imputations in Clustering, Dimension Reduction and Cell Cycle Dynamics. Four evaluation metrics, Entropy of accuracy (*H*_*acc*_), Entropy of purity (*H*_*pur*_), Adjusted Rand Index (ARI) and Normalized mutual information (NMI) were used to quantitatively evaluate the clustering performance with Louvain Clustering and K-means Clustering algorithms. Three labelled (i.e., cell type) datasets were used. The number of clusters was set to the number of known cell type labels in each dataset. Top 3000 high variable genes were picked out followed by running PCA. The top 10 PCs were used for clustering. ARI and NMI were subtracted by the results of the raw data (vice versa for *H*_*acc*_ and *H*_*pur*_ for better visualization) and the medians were taken. In cell cycle dynamics, cell cycle status and score of each cell was predicted by the Seurat algorithm. We further investigated whether the suspicious trans-differentiation from plasmablasts to developing neutrophils could be avoided with imputations. Data was pre-processed with adjustment for the percentage of mitochondrial gene expression, followed by running PCA and UMAP in Seurat. All UMAP plots were generated with first 50 PCs. Extreme values in heatmaps have been limited to a cutoff for better visualization. (**A-B**) Median values of the four metrics in Louvain and K-means Clustering. (**C**) Rank Sum tests were performed on the predicted cell cycle scores between cells of different known cell cycles (i.e., S vs. Others, G2M vs. Others) and the p-values were compared. (**D**) UMAP plots for ground truth cell cycles.

Cell cycle dynamics have been well studied at single cell level. Using Seurat Cell Cycle function on a cell cycle dataset with ground truth labels, we observed some improvements of the predicted cell cycle scores after imputed by afMF, ALRA, kNN-smoothing and MAGIC/MAGIC-log. The statistical significance of the comparisons of predicted cell cycle scores between different known cell cycles were improved in these imputations (**Figure 6C**). The cell cycle prediction accuracy and F1 scores were higher in afMF, ALRA and AutoClass as well (**Additional file 5, Figure S22**). As a results, more distinct and clearer separations between different cell cycles were also observed with data after imputation by MAGIC, afMF and ALRA in UMAP (**Figure 6D**).

### The impact of imputation on Pseudotime Trajectory Analysis may depend on the tools used

Pseudotime trajectory analysis enables the study of cell differentiation and development. Though it has been evaluated previously, here we used imputed data with three more well-established algorithms: DPT, monocle3, and Slingshot with additional datasets.

When using DPT, ALRA, afMF, MAGIC/MAGIC-log and ccImpute improved the pseudotime analysis (**Figure 7A-B** and **Additional file 6, Figure S23**). The predicted branches were further evaluated by the clustering metrics with true branch labels as ground truth. ALRA, afMF, ccImpute and AutoClass showed improvements in branch predictions (**Figure 7C**). In diffusion map visualization, afMF, ALRA and AutoClass showed better continuum trajectories and branches, while others either showed less improvements or heavily distorted the trajectory patterns (**Figure 7D** and **Additional file 6, Figure S24**).

**Figure 7.**
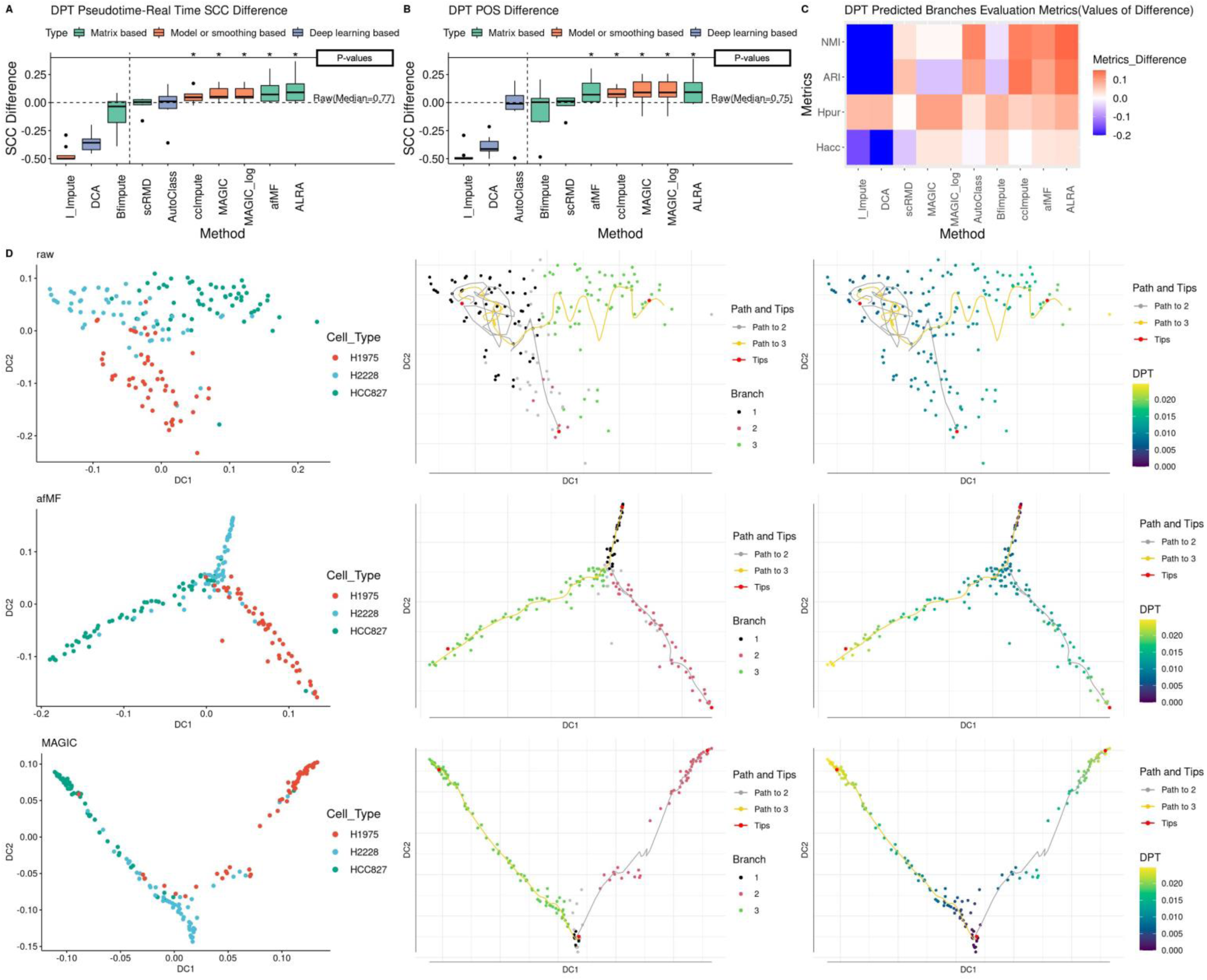
Performance of imputations in Pseudotime Trajectory Analysis with DPT. The performances of imputations on pseudotime trajectory analysis were evaluated with DPT using CellBench cellmix1-4 with real time & lineage labels. The starting point and end point were set as ‘tips’ based on known information with other parameters set to default. The metrics were subtracted by the values of the raw results. Extreme values have been limited to a cutoff value for better visualization. One, two and three ‘*’ represented ‘P<0.05’, ‘P<0.01’ and ‘P<0.005’ respectively and ‘.’ represented ‘P<0.1’, compared to raw log-normalization using Wilcoxon rank sum test. (**A-B**) Spearman correlation coefficients (SCC) and Pseudo-temporal ordering score (POS) were calculated between the predicted DPT values and the real lineage/time labels. (**C**) The predicted branches by DPT were further evaluated with clustering metrics *H*_*acc*_, *H*_*pur*_ ARI, and NMI and the true branches were used as ground truth. (**D**) 2-D diffusion map plots were generated for visualizing trajectories.

As shown in **Figure S25** for monocle3, higher Spearman correlations and pseudo-temporal ordering scores (POS) between predicted pseudotime and real time were observed in MAGIC/MAGIC-log with statistical significance. While afMF and AutoClass only showed slight improvements, the rest of the algorithms had no or negative influence. In PCA visualizations for cell types and pseudotime (**Additional file 6, Figure S25C-D**), afMF and ALRA showed relatively consistent patterns as no-imputation, while MAGIC-log only performed well in PCA but showed abnormal projections in UMAP as other algorithms.

On the other hand, Slingshot were found to be incompatible with most imputation algorithms as imputed data gave inferior or no improvement in trajectory analysis compared to raw data (**Additional file 6, Figure S26**). Therefore, Slingshot should not be used with imputed data.

### Imputation was compatible with advanced pathway analyses AUCell and SCENIC but not Cell-Cell Communication algorithms

AUCell is an advanced application for investigating the activities of gene sets (e.g., pathways) in each cell. As there is no ground truth, a well-studied pathway (i.e., interferon (IFN)) between COVID-19 and healthy controls were used. As shown in **Figure 8A-C**, in afMF and ALRA-imputed data, increased percentages of monocytes with Hallmark IFN-Alpha/Gamma response activated were only observed within COVID-19 subjects (true positive); in contrast, cells remained low levels of activation within healthy controls (true negative) as expected. On the other hand, other imputation algorithms showed IFN activation in both normal (false positive) and COVID-19 subjects (true positive) at the same time. When projected by UMAP, it is clear that after imputation by AutoClass activation of both IFN-Alpha and IFN-Gamma were found among healthy controls (normal cells), which were false positives activation. Similar conclusions were obtained for GO IFN-related pathways (**Figure 8D**). These results suggested that only Matrix based algorithms (ALRA and afMF) better revealed activation of IFN pathways in monocytes in COVID-19 patients while avoiding having excessive false positive activation among healthy controls.

**Figure 8.**
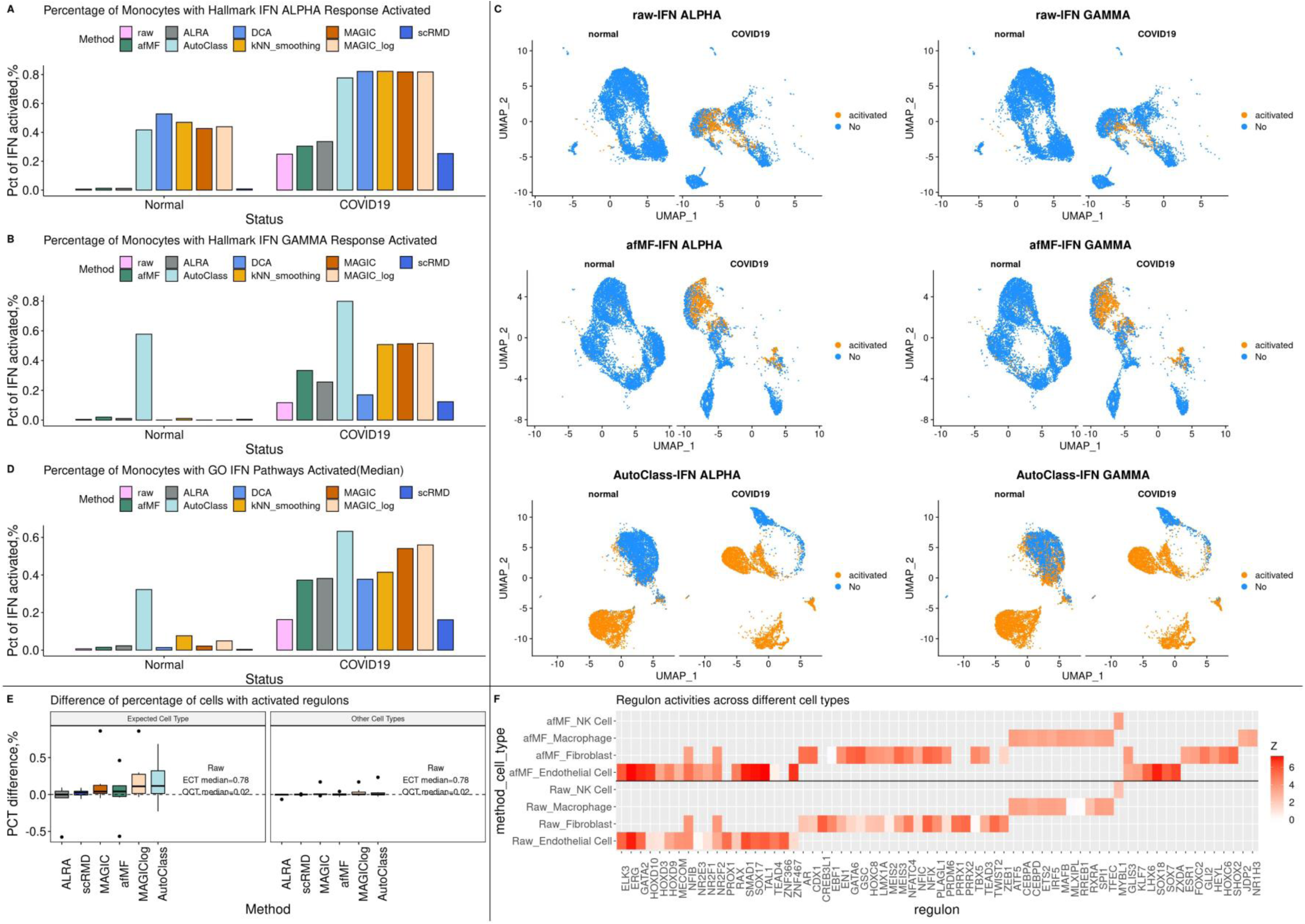
Performance of imputations in advanced applications: AUCell and SCENIC. The compatibility between AUCell and different imputations were evaluated using GSE155673, where the activations of the interferon (IFN) related pathways between COVID-19 and healthy controls were compared among monocytes. The IFN-related pathways were collected from Hallmark gene sets (2 terms) and GO gene sets (31 terms). The compatibility between SCENIC and different imputations were evaluated using GSE115978. Some metrics were subtracted by the values of the raw results. (**A-D**) Each cell was assigned a binary status (activated or not) for each pathway by the AUCell pipeline and the percentages of monocytes with activated IFN pathway were compared between COVID-19 and healthy controls. (**E**) seven well-established regulons (i.e., TCF7, EOMES, TBX21, MITF, MYC, MAFB, PAX5) as demonstrated previously were used to determine the activation status in a selected cell. A cell-type-specific regulon was expected to be activated only in the specific cell type, but not the other cell types. The percentages of cells that with activated regulons within expected cell type (ECT) and other cell types (OCT) were calculated and compared. (**F**) The resulted regulons with Z-score≥3 for raw and afMF were collected and compared.

SCENIC is an advanced application that incorporates AUCell for exploring gene regulatory networks and transcriptional factors(22). Similar as for IFN in AUCell, seven well-established cell-type-specific ‘regulons’ were selected as previously demonstrated(64). afMF, MAGIC/MAGIC-log and AutoClass performed well as they increased the percentages of cells with activated regulons within expected cell types (correct calls) while remained consistent levels as no-imputation within other unrelated cell types (false positive calls) (**Figure 8E**). Next, we compared all the identified regulons (Z-score>3) across all the cell types and observed that most selected imputation methods could recover the patterns in raw data but also added some unique significant regulons (**Figure 8F** and **Additional file 7, Figure S27**). Of note, these newly generated significant regulons should be further validated through other experiments.

CellPhoneDB(23) and CellChat(24) are two popular tools that enable researchers to study cell-cell communication networks through ligand-receptor databases. However, as shown in **Additional file 7, Figure S28-29**, huge increments of significant interactions were discovered after imputations. Though no ground truth is available for demonstration, they were believed to be the false positives as most of the pairwise cell-cell interaction were detected only after imputation (e.g., MAGIC, AutoClass, afMF). Similar as for CellChat, we also found large increase of significant interactions after imputation (**Additional file 7, Figure S30**); on the other hand, we discovered that CHIP status had more interactions than NO-CHIP status after imputation, which was expected.

### Imputed data showed better consistent with additional wet-lab data such as bulk RNA-sequencing and protein expression

Additional support of using imputation can be gathered from datasets with additional data after cell sorting or protein assay to validate transcriptome results. For one cell type, gene expression measured from bulk RNA-seq are more accurate than single cell RNA-seq. Bulk RNA-seq measures gene expression of groups of cells and thus better reflects the ground truth in some circumstances, i.e., when no need to consider cell-cell variability. Using three mixture datasets with matched bulk data, higher relative (same minus others) and absolute (same) Spearman correlations between same-cell-type single cell/pseudobulk (median-aggregated) and bulk profiling were observed for most algorithms with statistical significance, except for scRMD, Bfimpute and DCA (**Figure 9A-B** and **Additional file 8, Figure S31A-D**). Correlations of the pairwise cell type logFC between pseudobulk and bulk were higher in most imputations as well, except for Bfimpute and ccImpute (**Figure 9C** and **Additional file 8, Figure S31E**).

**Figure 9.**
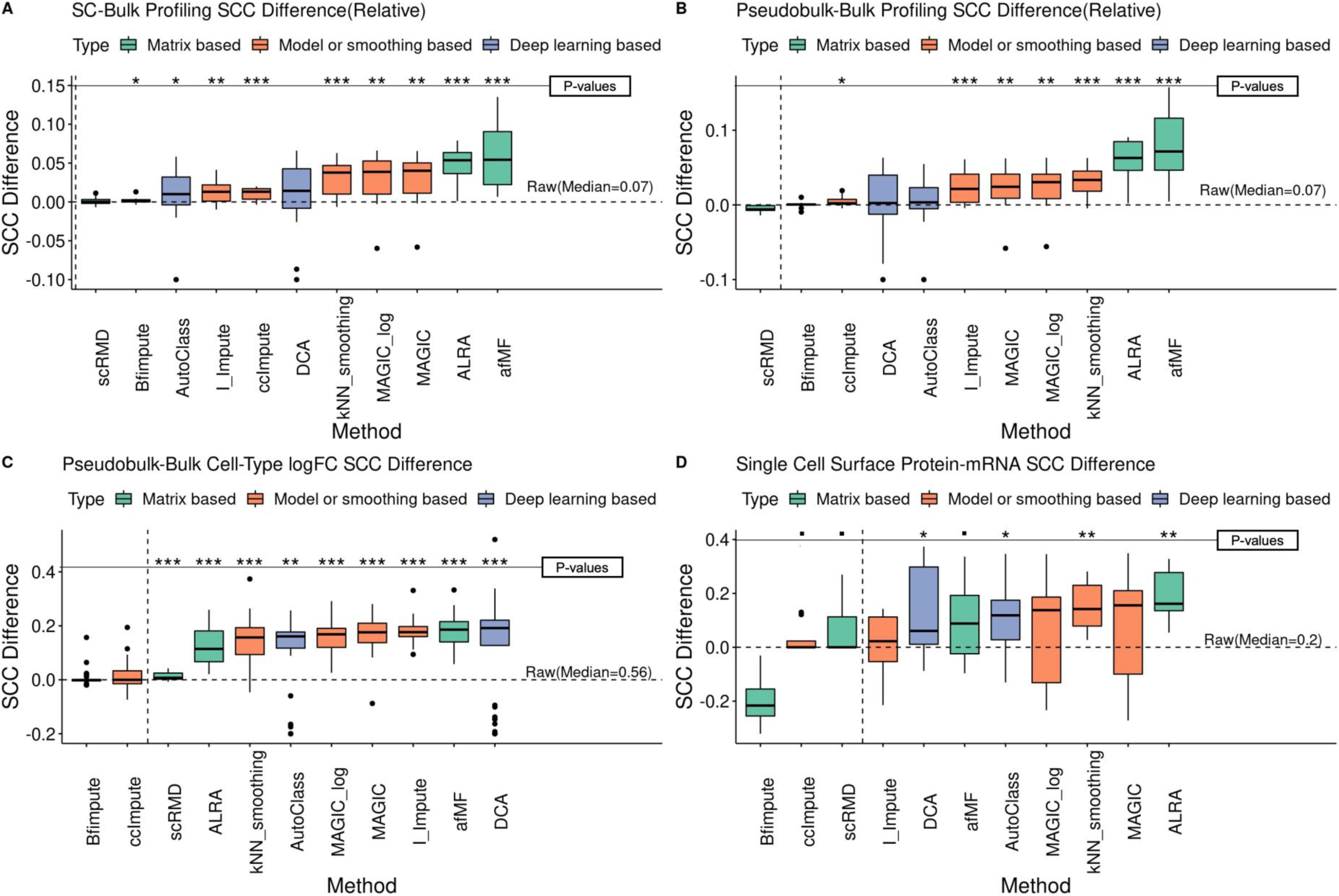
Performance of imputations in other supporting analysis. Supporting analysis provided other supportive evidence for imputations in addition to downstream applications, including SC-Bulk Profiling Similarity, mRNA-Surface Protein Correlation, Distribution similarity between scRNA-seq and FISH, and Cell-Cell Correlation. The metrics were subtracted by the values of the raw results. Extreme values have been limited to a cutoff value for better visualization. One, two and three ‘*’ represented ‘P<0.05’, ‘P<0.01’ and ‘P<0.005’ respectively and ‘.’ represented ‘P<0.1’, compared to raw log-normalization using Wilcoxon rank sum test. (**A**) Spearman correlation coefficients (SCC) were calculated for each single cell profile and the bulk profile (within same group/condition). Next, the same-group SCCs were subtracted by the max different-group SCC and the median value of each group was used for comparison. (**B**) The same analysis was performed for pseudobulk profile (i.e., median-aggregated of the same cell type). (**C**) Pair-wise calculations of log fold changes (logFC) between different groups in both pseudobulk profiles and bulk profiles were conducted and their SCCs were calculated. (**D**) Six marker gene mRNAs in two tissues (CD4, CD2, CD19, CD14, CD34, CCR7 in PBMC and CBMC in GSE100866) with corresponding surface protein measurement were compared. SCCs were calculated between the two measurements for each gene.

Though not all mRNA measured in scRNA-seq will translate to protein, some correlations between mRNA and protein are still expected. Using a CITE-seq dataset (GSE100866 PBMC and CBMC) with both mRNA and surface protein measurement, higher Spearman correlations between selected mRNA and surface protein were observed in ALRA, MAGIC/MAGIC-log, kNN-smoothing, AutoClass, afMF and DCA (**Figure 9D** and **Additional file 8, Figure S31F**).

### Simulated data analysis further supported the benefits of imputation

Real-life scRNA-seq data are complicated and difficult to simulate. Nonetheless, simulated datasets have the advantage that they come with the ground truth. Using simulated datasets generated from Splatter(42) and SplatPop(43), we found afMF, ALRA, MAGIC/MAGIC-log, AutoClass and kNN-smoothing performed generally better in DE analysis (MAST), Classification, Biomarker Prediction, Automatic Cell Type Annotation, and Cell-Ground Truth Profiling Similarity (**Additional file 9, Figure S32-33**). Only MAGIC/MAGIC-log, ALRA and kNN-smoothing showed improvements in simulated data K-means clustering (**Additional file 9, Figure S33J**), while all methods performed worse with Louvain clustering (**Additional file 9, Figure S33K**). In cell-cell correlation heatmaps for Mock90 dataset, afMF, ALRA and MAGIC / MAGIC-log improved the visualizations (**Additional file 9, Figure S34**).

### Running time, memory usage and recommendation

Good imputation algorithms should have acceptable running time and memory usage. Using four datasets with 1500, 5000, 10000 and 50000 cells, most of the algorithms showed acceptable running time (i.e., within 10 hours), except for I-Impute (**Figure 10A**). Meanwhile, only ccImpute and Bfimpute showed unacceptable memory usage (i.e., >512 GB) on large datasets (**Figure 10B**). Next, performance of different imputation algorithms on different downstream applications & supporting analysis were rated by comparison with no imputation (raw). Better performance was defined if enhancement was seen compared with raw data and it was further divided into two rating: slightly better and generally better. Generally, afMF, ALRA and MAGIC/MAGIC-log had relatively stable performance while others showed less or no improvements or were incompatible with various downstream tools (**Figure 10C** and **Additional file 10, Table S1**).

**Figure 10.**
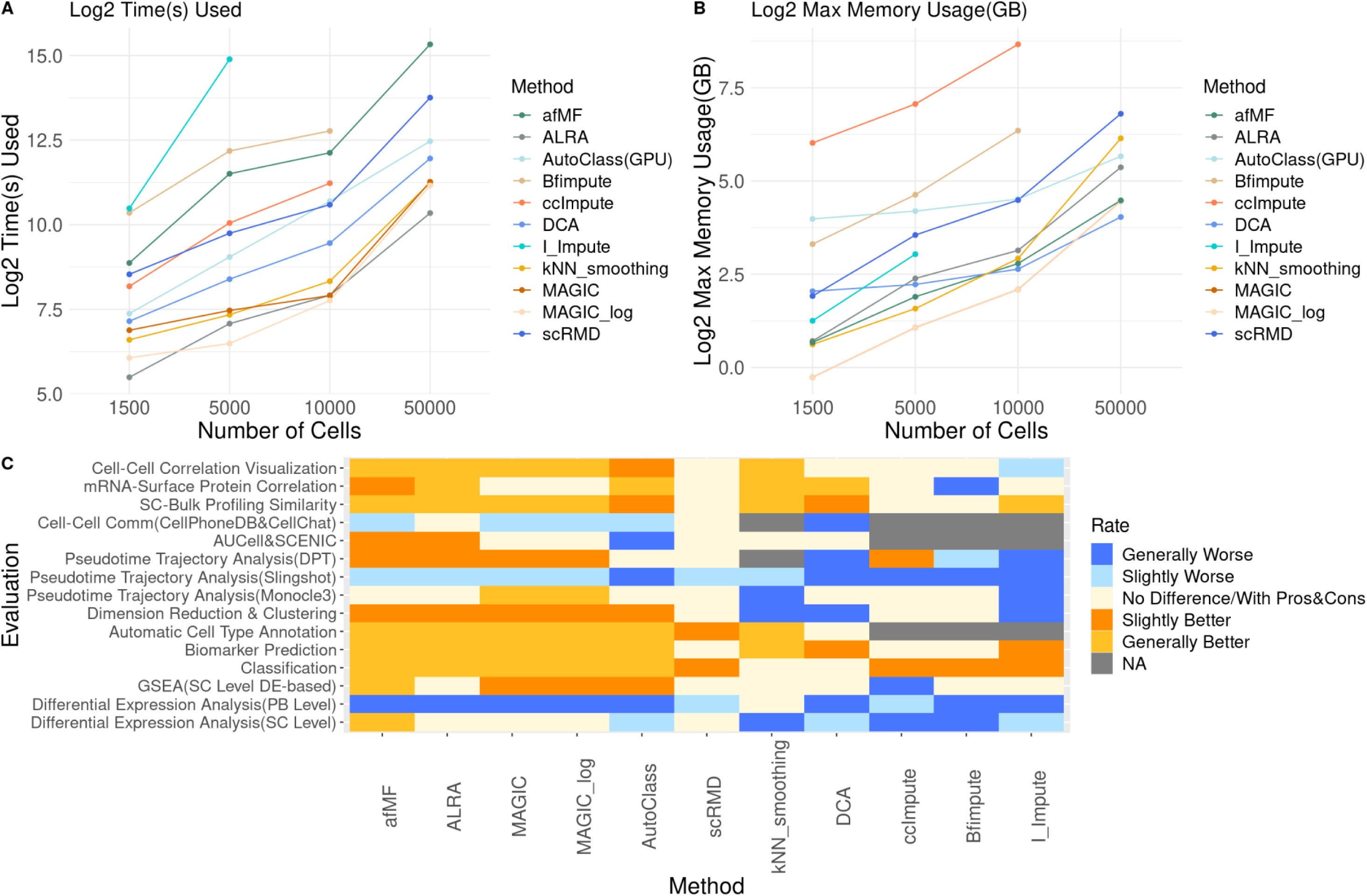
Running time, memory usage and recommendations for imputation algorithms. Four sizes of datasets (10000×1500, 10000×5000, 10000×10000 and 10000×50000 matrix) were applied to evaluate the (**A**) time spent and (**B**) memory usage of different imputation methods. (**C**) Based on the evaluations, a summary heatmap was created. The performances were classified into five levels ‘Generally Worse’, ‘Slightly Worse’, ‘No Obvious Difference’, ‘Slightly Better’ and ‘Generally Better’. Note that algorithm-evaluation were labelled as ‘No Obvious Difference’ if that algorithm had both obvious advantages and disadvantages based on different metrics in that evaluation.

We further compared the ALRA and the improved afMF performance in various scRNA-seq analysis. **Additional file 10 Table S2** detailly showed the performance of the two algorithms. While performances of many downstream analyses were comparable, afMF gave better results on DE analysis, GSEA, classification and biomarker prediction, and cell clustering.

## Discussion

In this study, we performed matrix completion by implementing an iterative full matrix factorization algorithm (afMF) to handle dropouts in scRNA-seq datasets. We demonstrated in detail that which imputation algorithm could be beneficial to which type of downstream applications (**Additional file 10, Table S1**). We showed that matrix-based methods such as afMF and ALRA had relatively stable performance and improved various downstream analyses. They also had good scalability which could be applied to data matrix of different size. It is encouraging to see that afMF ranked among the top imputation algorithms in evaluations of downstream analyses, such as DE analysis, GSEA, Classification, Biomarker Prediction, Automatic Cell Type Annotation, Dimension Reduction & Clustering, Pseudotime Trajectory Analysis with DPT, AUCell & SCENIC, SC-Bulk Profiling Similarity and mRNA-Surface Protein Correlation. Besides, MAGIC (smoothing based algorithm) and AutoClass (deep learning algorithm) also showed some enhanced output in selected applications but produced false positives in other applications.

There are several algorithms based on matrix theory technique to perform the task of matrix completion to impute zero counts in scRNA-seq matrix. scRMD(56) targeted to determine the matrices for dropout and noise. Although the concept is good, it needs solve two more additional latent matrices in addition to the underlying latent expression matrix. Bfimpute(44) is different from most other low-rank MF algorithms in that it subdivided cells into different cell-groups before performing Bayesian based MF. There are also other matrix theory algorithms using different optimization methods. afMF is different from ALRA in that an iterative process is used to optimize two low-rank matrices which may account for the added benefits shown in these evaluations.

In addition to the previous established evaluation metrics, we provided more in-depth downstream analyses incorporating current well-known single cell tools to benchmark imputation methods and explored their compatibilities: (1) Pseudobulk DE Analysis; (2) GSEA; (3) Automatic Cell Type Annotation; (4) Pseudotime Trajectory Analysis with three popular tools; (5) AUCell; (6) SCENIC; (7) Cell-Cell Communication. While most analyses are compatible with prior imputed data, we revealed one new but important feature in the evaluation of pseudobulk DE analysis. Most imputation methods gave inferior results in pseudobulk DE analysis. We reasoned that pseudobulk processing may serve as a smoothing step. Therefore, it may be already less affected by the influence of dropouts and prior imputation of data may make the data less compatible with pseudobulk DE analysis. Besides, the compatibility between Pseudotime Trajectory Analysis and prior data imputation is heavily depended on the algorithms used. In contrast, imputation was likely to produce false positives in Cell-Cell Communication.

Cell-Cell Communication such as CellPhoneDB used mean values throughout the analysis and should be sensitive to dropout zero counts. The exact reason why it gave inferior results when using imputed data is not clear. We reasoned that imputation may somehow add noise to the mean values of ligands and receptors and thus produced suspicious high level interaction results. Additionally, imputation showed limited or no improvement in 2 out of 3 popular pseudotime trajectory analysis (i.e., Monocle3 and Slingshot), which was also reported by Hou et al.(9) when they applied different trajectory tools to imputed data. Note that we had used other trajectory tools such as Monocle2 with DDRTree and TSCAN, but they showed worse compatibilities with most imputation methods as well (data not shown). It is suggested that imputation may not be helpful for describing cell-to-cell or gene-to-gene interaction and thus downstream analyses that utilized such information gave inferior results with imputation. One example is the inferior performance using imputed data with Slingshot that probably due to the incompatibility between imputation and its internal trajectory algorithm.

Our evaluations revealed that matrix decomposition / factorization-based methods such as afMF and ALRA did not adversely distort the data matrix while handling dropouts. As a class, Matrix-theory algorithms was demonstrated to have the top performance with most downstream analyses; in contrast, many deep-learning-based algorithms or model-based algorithms were found to overfit the data or generate artefact data structure leading to false positive findings. It is believed that the simpler imputation algorithms such as matrix factorization may be preferred for various scRNA-seq design.

As imputation algorithms are emerging these years, little attention has been put on the compatibility issue. From biologist perspective, it is essential to have an imputation method that is not only to impute the dropout, but also to improve various downstream analyses. Indeed, researchers do not know if they should do prior data imputation in real scRNA-seq analysis so far. Imputation is more widely used in eQTL study(68) and, ironically, in pseudotime trajectory analysis(69). According to our evaluations (i.e., DE analysis), imputation is a great option for conducting cell-type-specific eQTL study. Interestingly, imputation was not only applied in scRNA-seq data, but also in qPCR data and genomic data. For instance, genomic variant imputation(70) has now been widely used in current research. Our study provided a comprehensive illustration of imputation in scRNA-seq, and the methods and pipeline may be extended to other techniques such as spatial single cell transcriptomics.

Unimputed raw data was proposed to have useful information as well(6), and therefore comparing the results between raw and imputed data is necessary. Imputation can provide higher power to detect biological signals but can still introduce false positive results to some extent. Similar as current single cell analysis pipeline, multiple slots such as ‘raw’ and ‘imputed’ in current scRNA-seq object could be used to provide more convincing results.

The size of the scRNA-seq data is getting larger and larger and thus imputation could take much time and memory usage. Therefore, algorithms with insufficient scalability were not evaluated in this study. According to our results, imputation could be applied on purified cell type or subset data for downstream analysis and thus dramatically reduce time spent and memory usage. Additionally, GPU could be used to accelerate the process as well.

Those downstream tools that are designed for analysis of spare data matrix, can reduce the effect of the dropout problem in various ways. This raised another question: which is better, imputation in advance vs. algorithms designed for handling sparse data. For example, many model-based algorithms including a zero-inflated model will not be benefited by using prior imputed data. This may somehow explain why imputation does not work well when using some downstream tools.

It is worth to note that benchmarking various imputation algorithms with various downstream analyses on various datasets is a time-consuming and space-consuming work. Therefore, the data and tools used here are still limited. Our study is served as a comprehensive demonstration for the popular downstream applications, and it is encouraged that researchers can conduct the imputation benchmark for each downstream application individually(71) with more datasets, metrics, and tools in the future. Additionally, it would be of great interest to develop downstream applications with internal implementation of imputation algorithms.

## Conclusions

In this study, we implemented an improved algorithm afMF to handle dropouts and demonstrated in detail with benchmark framework. The matrix-based methods such as afMF and ALRA had relatively stable performance while kept acceptable scalability. We hope these detailed evaluations and the algorithm developed in this study can enhance the use of imputation in various scRNA-seq downstream analyses as an addition to the raw data analysis, and further promote new discoveries.

## List of abbreviations

scRNA-seq: single cell RNA sequencing
HK gene: housekeeping gene
PCA: principal component analysis
UMAP: uniform manifold approximation and projection
DE analysis: differential expression analysis
GSEA: gene set enrichment analysis
AUC: area under curve
FC: fold change
IFN: interferon
GO: gene ontology
RF: random forest
DEG: differential expressed gene
AUROC: area under the receiver operating characteristic curve
ARI: adjusted rand index
NMI: normalized mutual information
POS: pseudo-temporal ordering score
PBMC: peripheral blood mononuclear cells
CBMC: cord blood mononuclear cells
SC: single cell
SCC: Spearman correlation coefficients.

## Declarations

### Ethics approval and consent to participate

Not applicable.

### Consent for publication

Not applicable.

### Availability of data and materials

The experimental data that support the findings of this study are available in public data domains (e.g., GEO and EGA). All public data analyzed during this study are shown in **Table 1**. The code for the benchmark evaluations and the new imputation algorithm ‘afMF’ is available at: https://github.com/GO3295/SCImputation.

### Competing interests

NT is founding Director and shareholder of the biotech startup company, Cytomics Ltd, in Hong Kong Science Park. AC was an intern in Cytomics Ltd during his final undergraduate year when he developed the afMF algorithm.

### Funding

Not applicable.

### Authors’ contributions

JH contributed to the design of the work, performed data curation, analysis, interpretation of data, and the creation of new software used in the work, and drafted and revised the work. ACMC contributed to the design of the algorithm and the creation of new software used in the work. NLST conceived the research, contributed to the design of the work, interpretation of data, and drafted and revised the work. SCPY contributed to the conception and design of the algorithm. All authors read and approved the final manuscript.

## Acknowledgements

Not applicable.

