## Supplementary figures and images for "Low-Rank Full Matrix Factorization for dropout imputation in single cell RNA-seq and benchmarking with imputation algorithms for downstream applications"

### Additional file 1

#### Figure S1. Pre-screening for twenty imputation algorithms

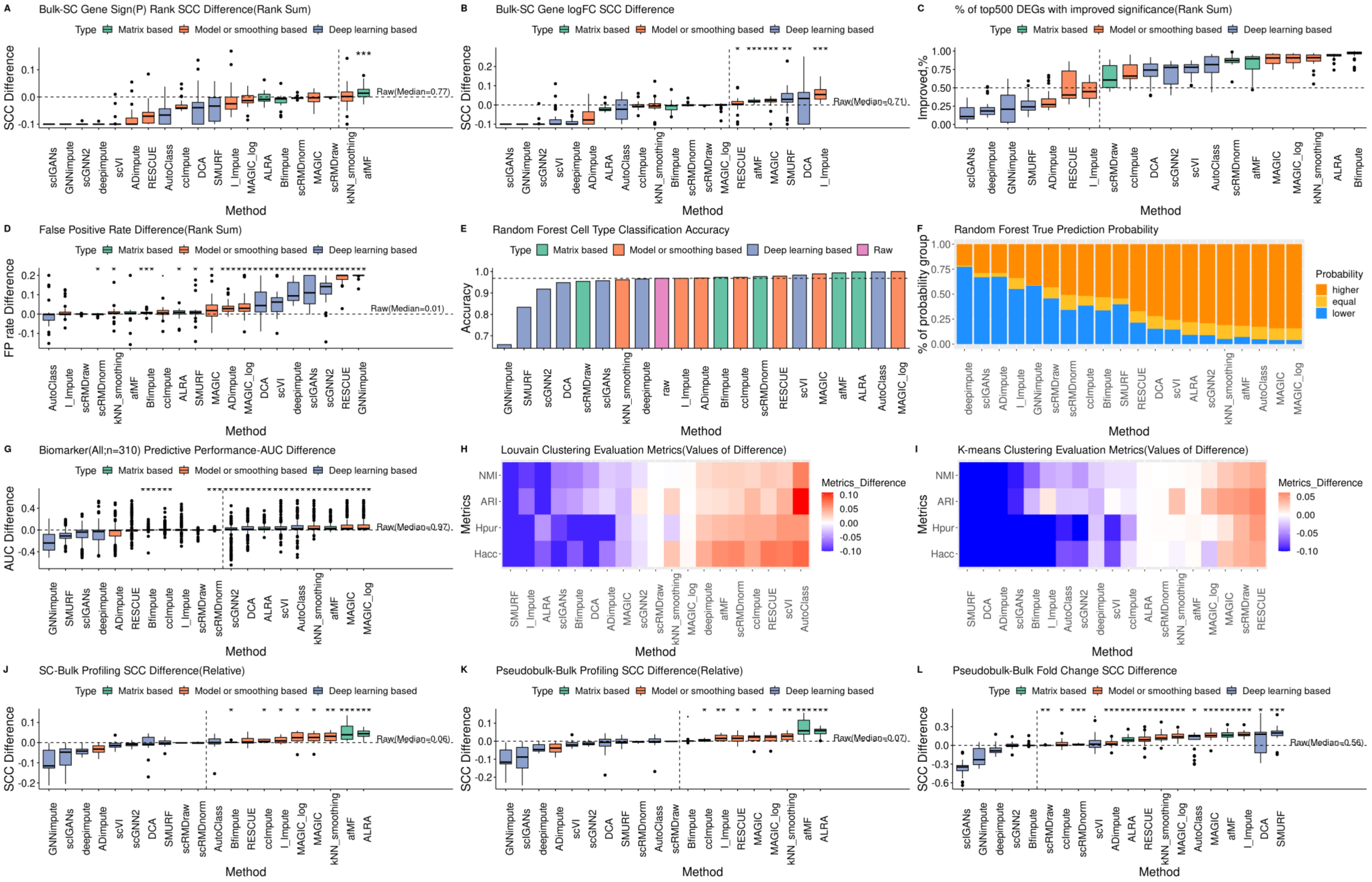
