## Additional file 2 for "Low-Rank Full Matrix Factorization for dropout imputation in single cell RNA-seq and benchmarking with imputation algorithms for downstream applications"

**Figure S2. Gene expression violin plots for GAPDH**

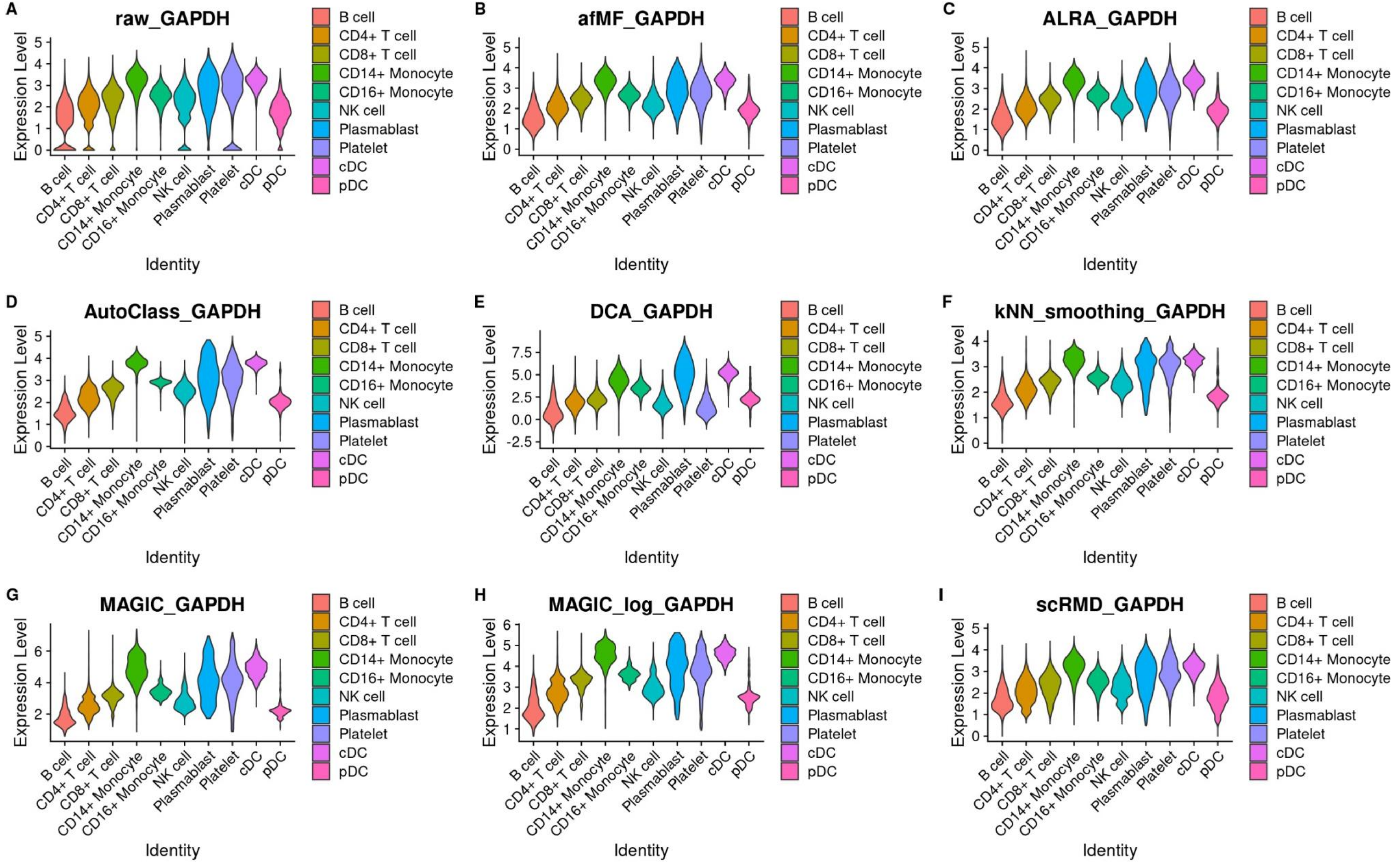

**Figure S3. Gene expression violin plots for CD8A**

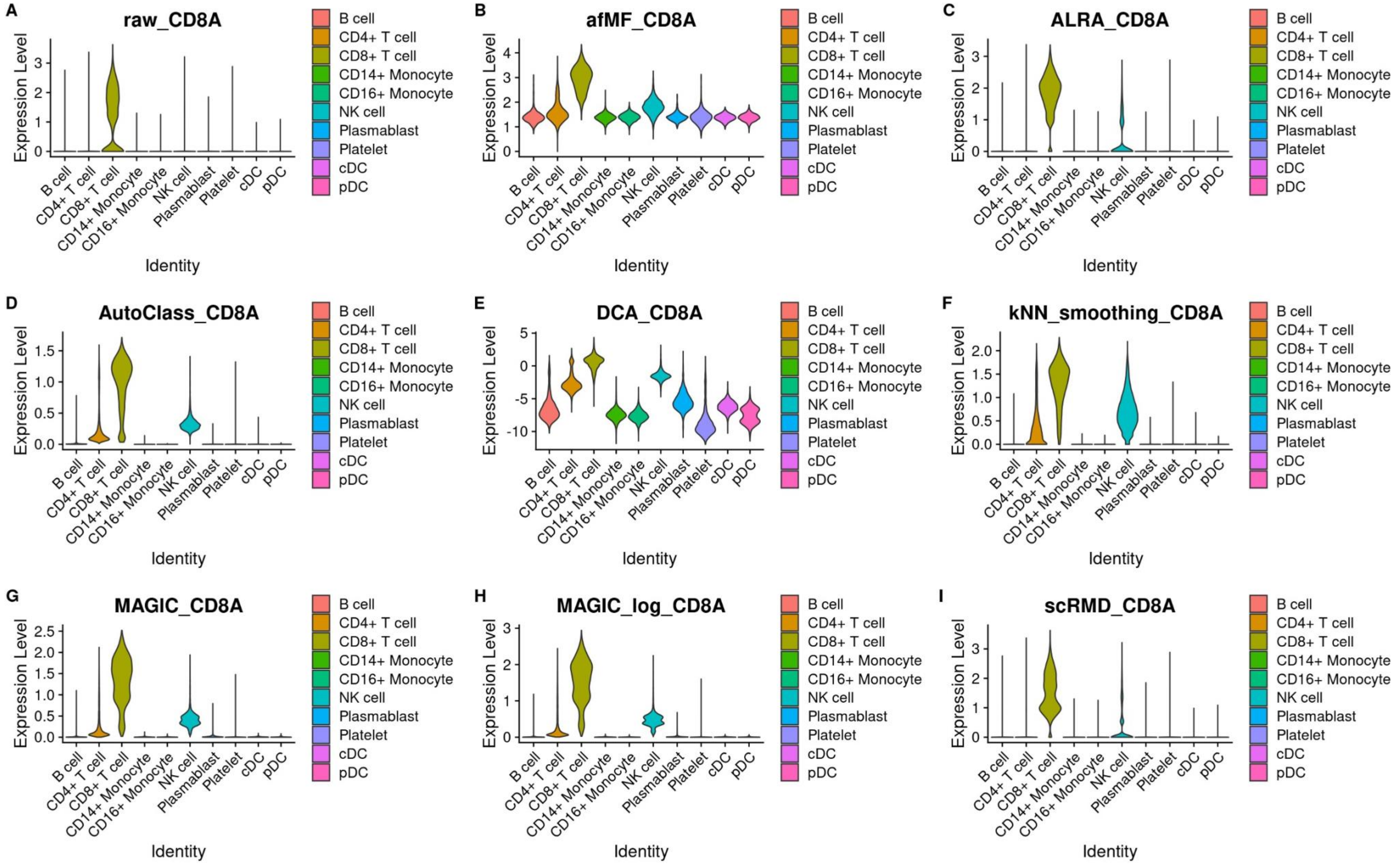

**Figure S4. Gene expression violin plots for CD19**

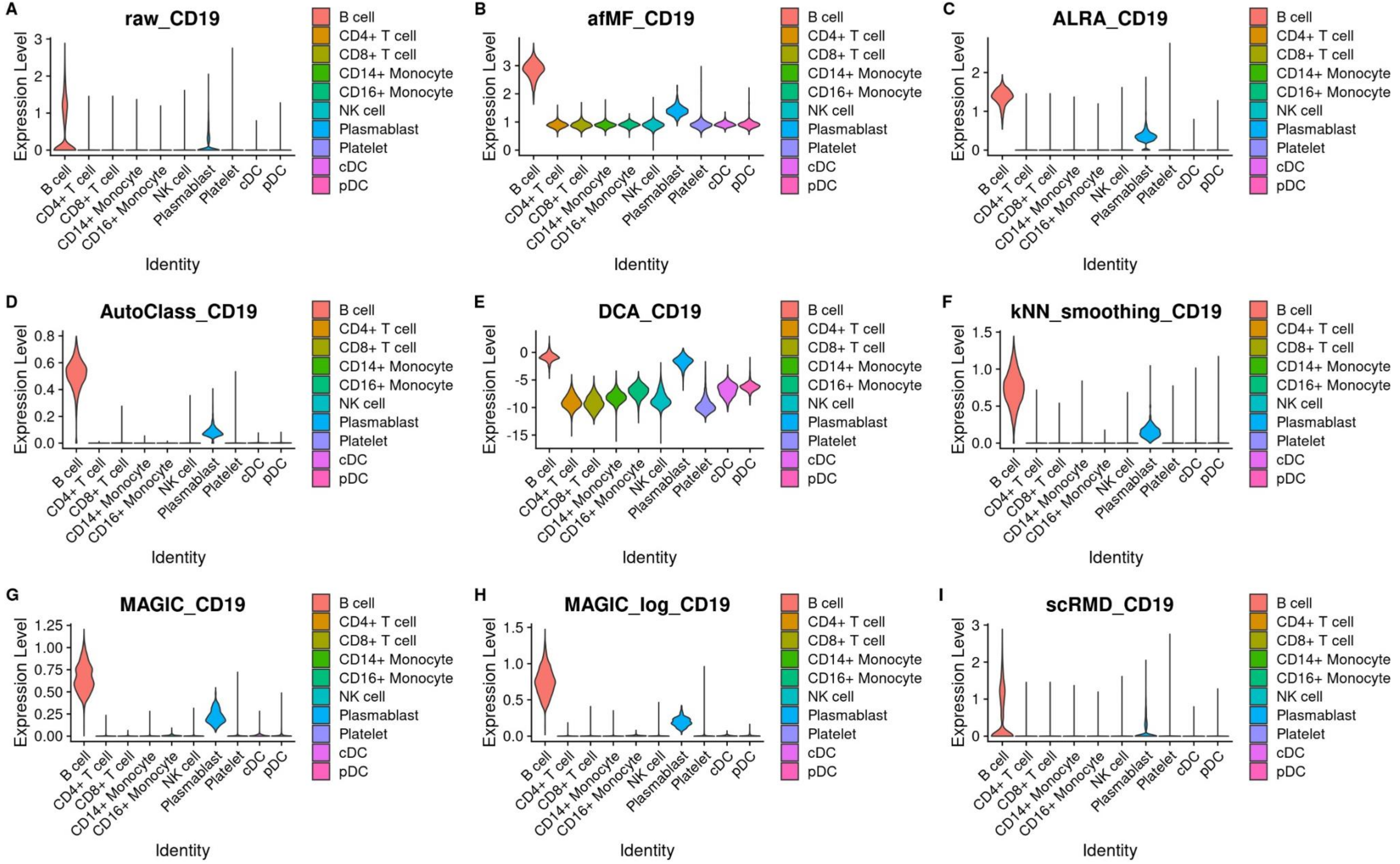

**Figure S5. 2-D PCA plots for simulated dataset Mock90**

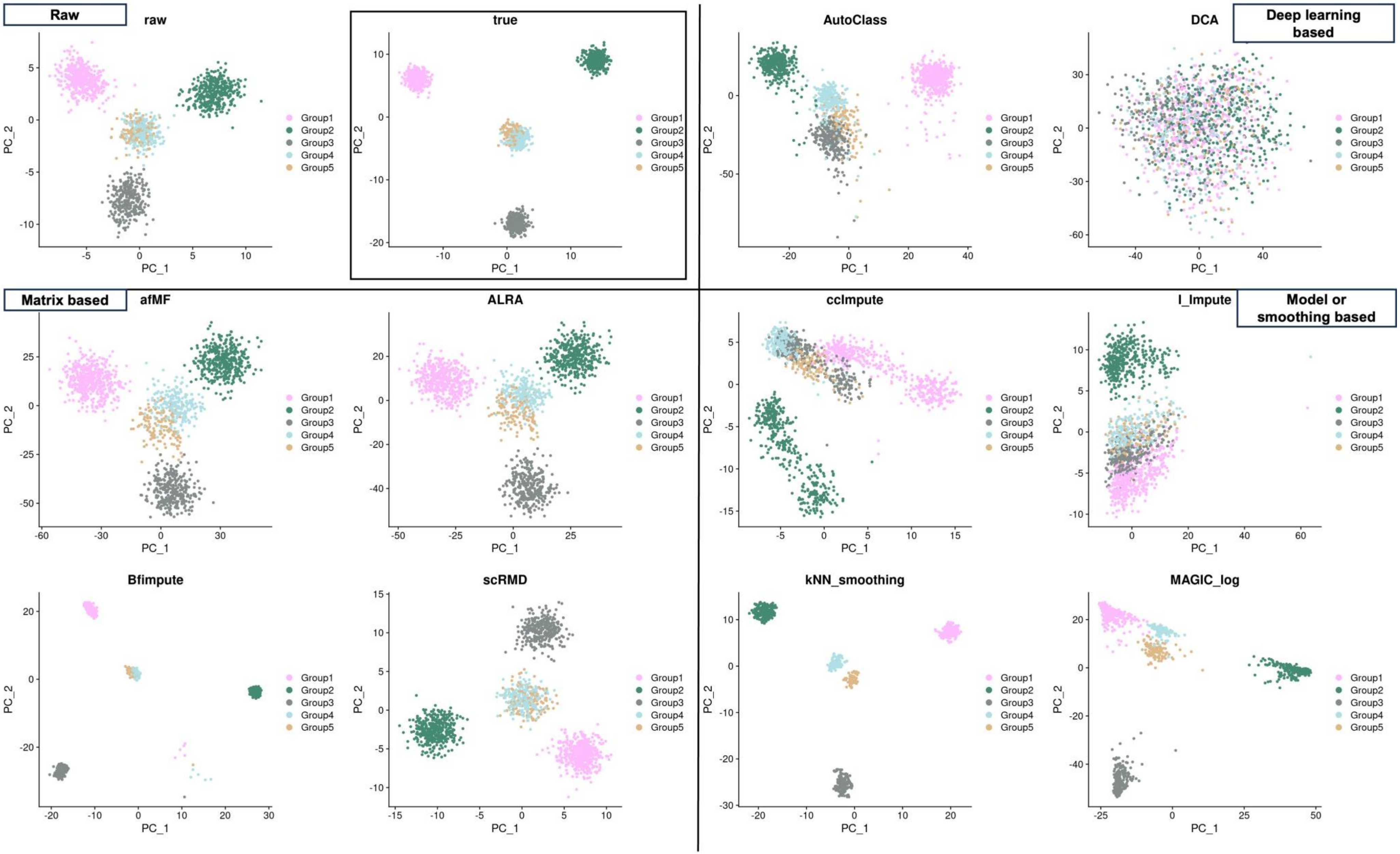

**Figure S6. 2-D PCA plots for simulated dataset SplatPop90**

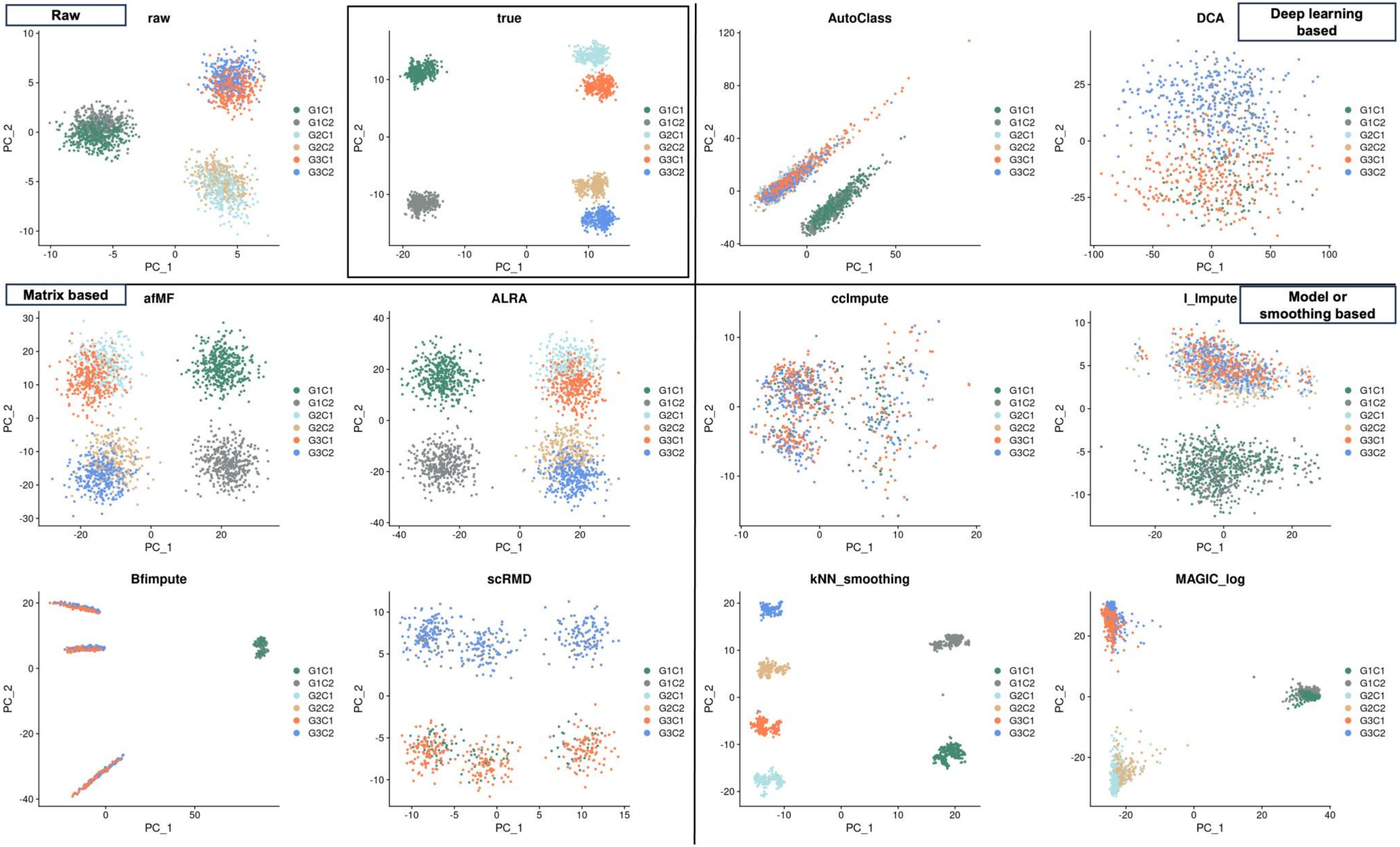

### Figure S7. Cell-Cell Correlation heatmaps in different imputations using CellBench-10X5CL

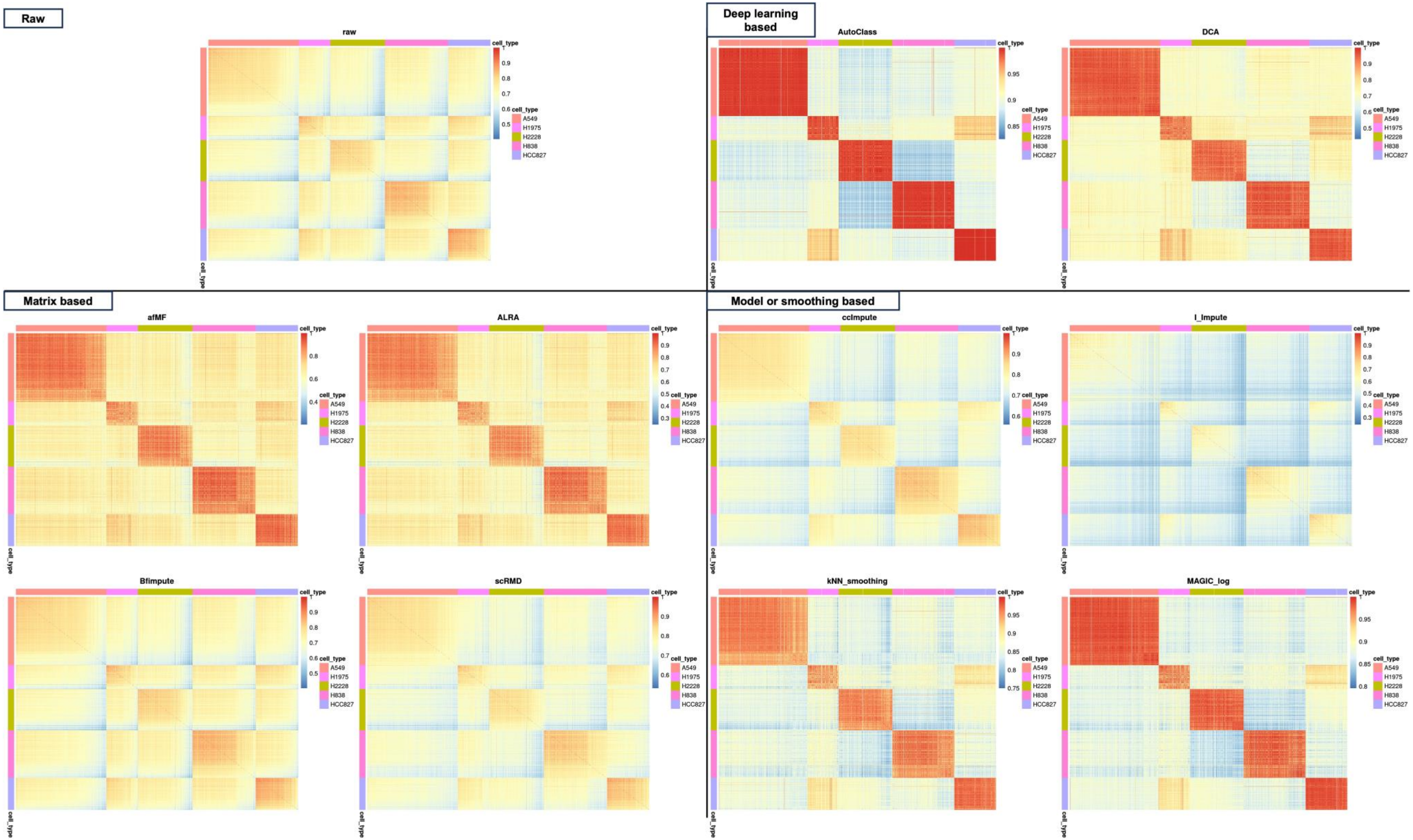
