## Additional file 3 for "Low-Rank Full Matrix Factorization for dropout imputation in single cell RNA-seq and benchmarking with imputation algorithms for downstream applications"

**Figure S8. Heatmaps showing each comparison in Differential Expression Analysis with MAST**

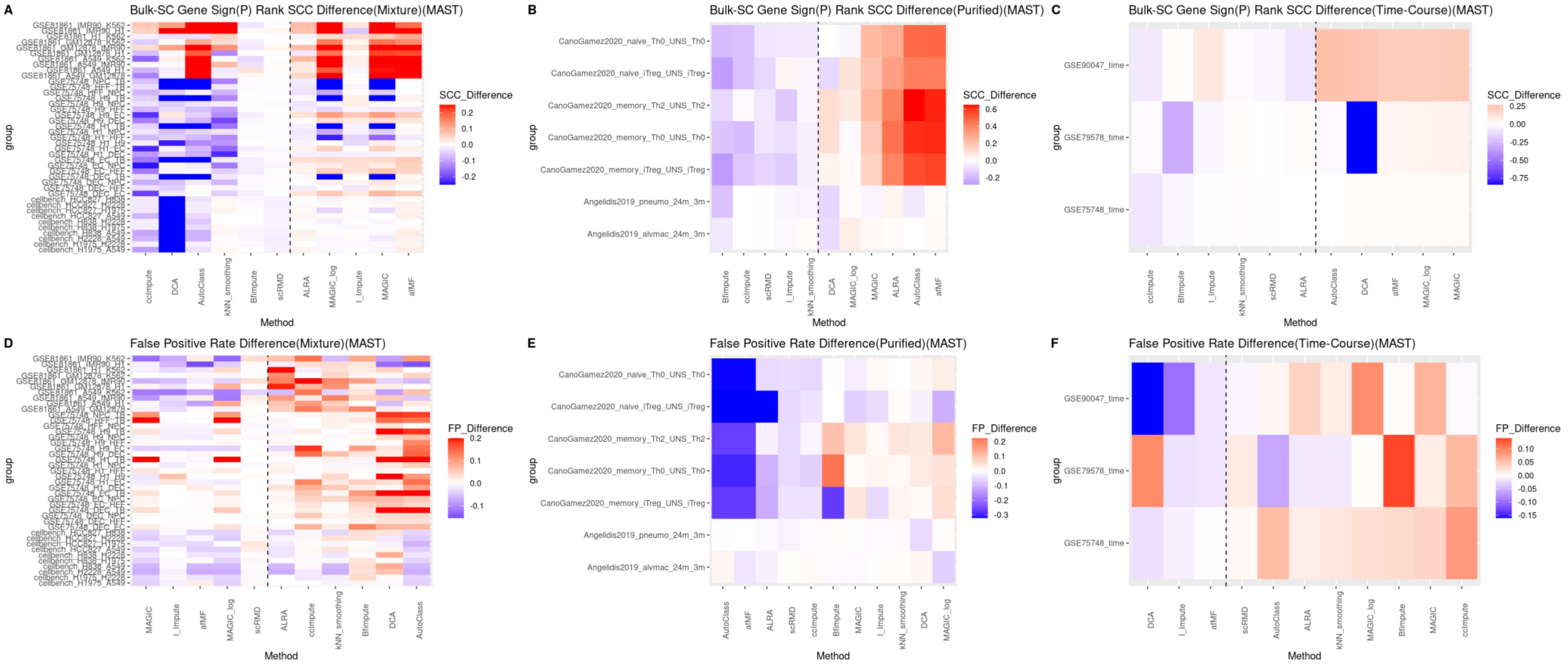

**Figure S9. Performance of imputations on Differential Expression Analysis with Wilcox Rank Sum test**

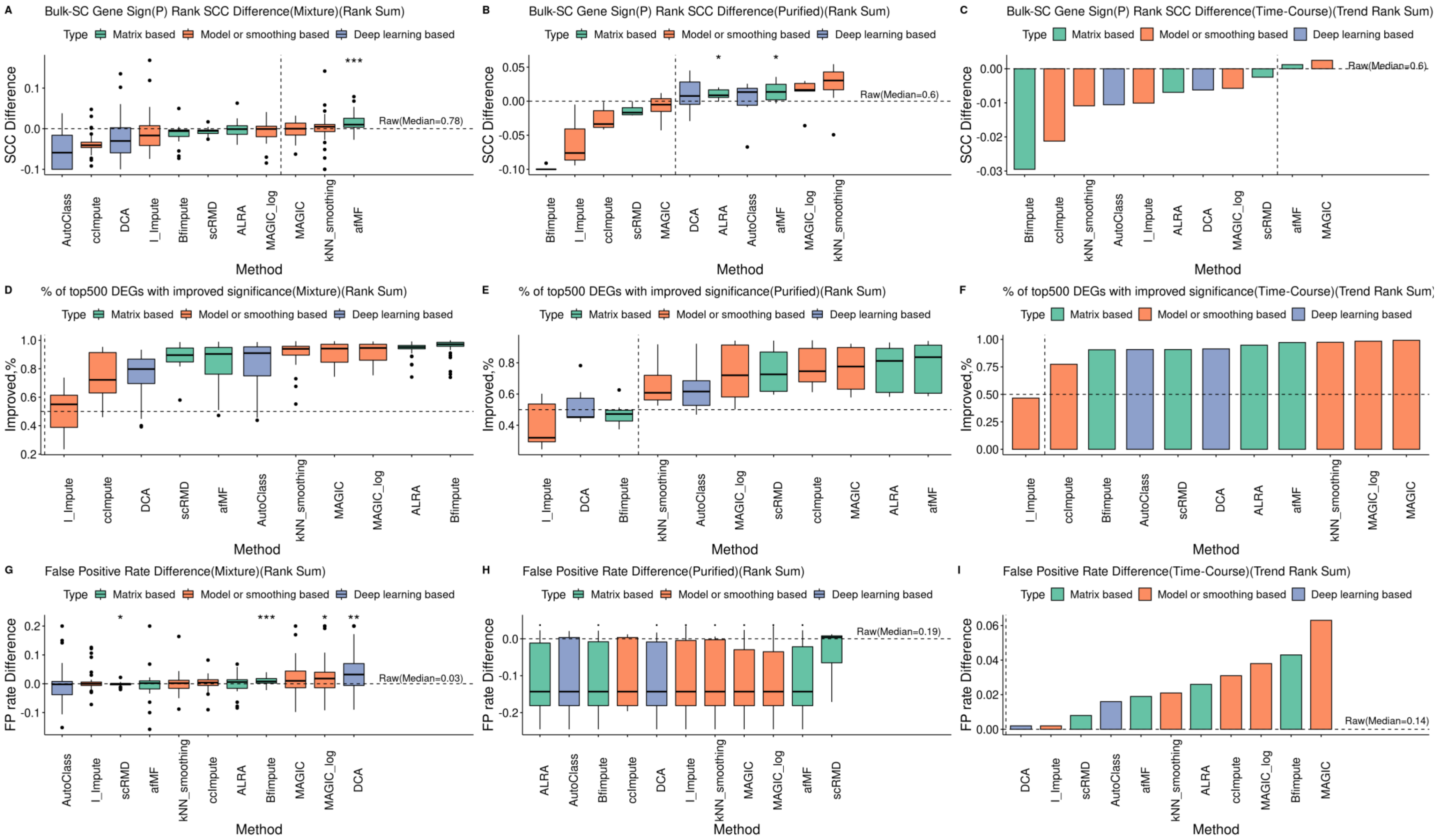

**Figure S10. Performance of imputations on Differential Expression Analysis with MAST & Wilcox Rank Sum test: SCC for Top 1000 DEGs**

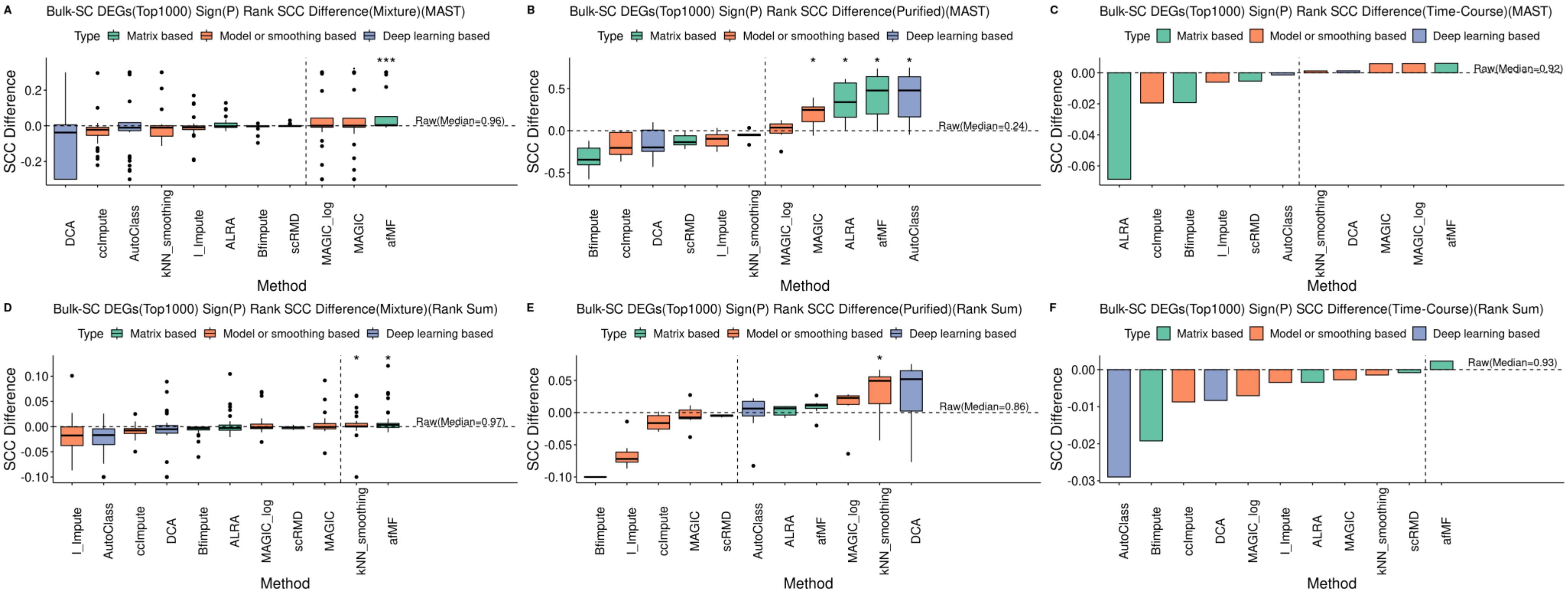

**Figure S11. Performance of imputations on Differential Expression Analysis with MAST & Wilcox Rank Sum test: false positive rates (percentile tail-ranked genes in bulk presented in top 500 single cell DEGs)**

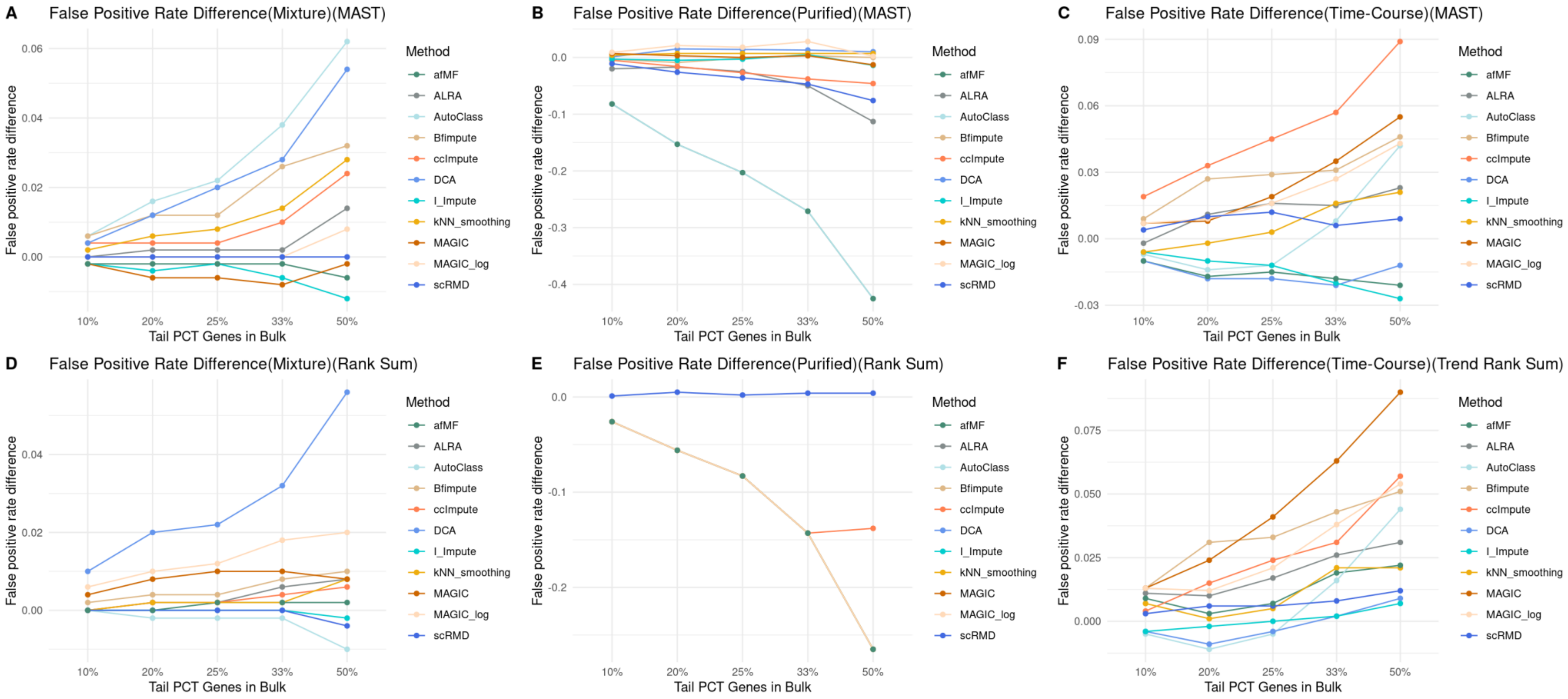

### Figure S12. Performance of imputations on Differential Expression Analysis with MAST: correlations of logFC

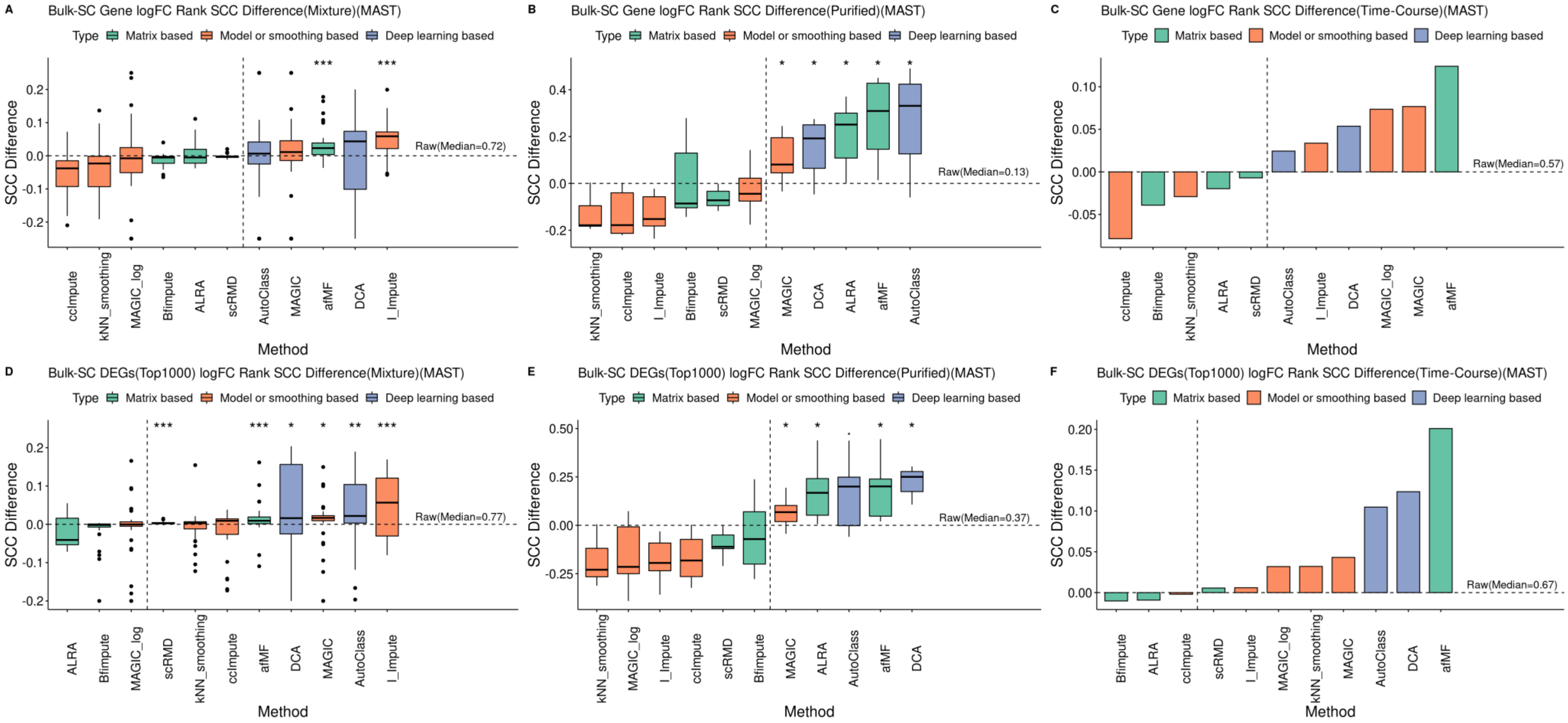

**Figure S13. Performance of imputations on Differential Expression Analysis with pseudobulk-limma-trend**

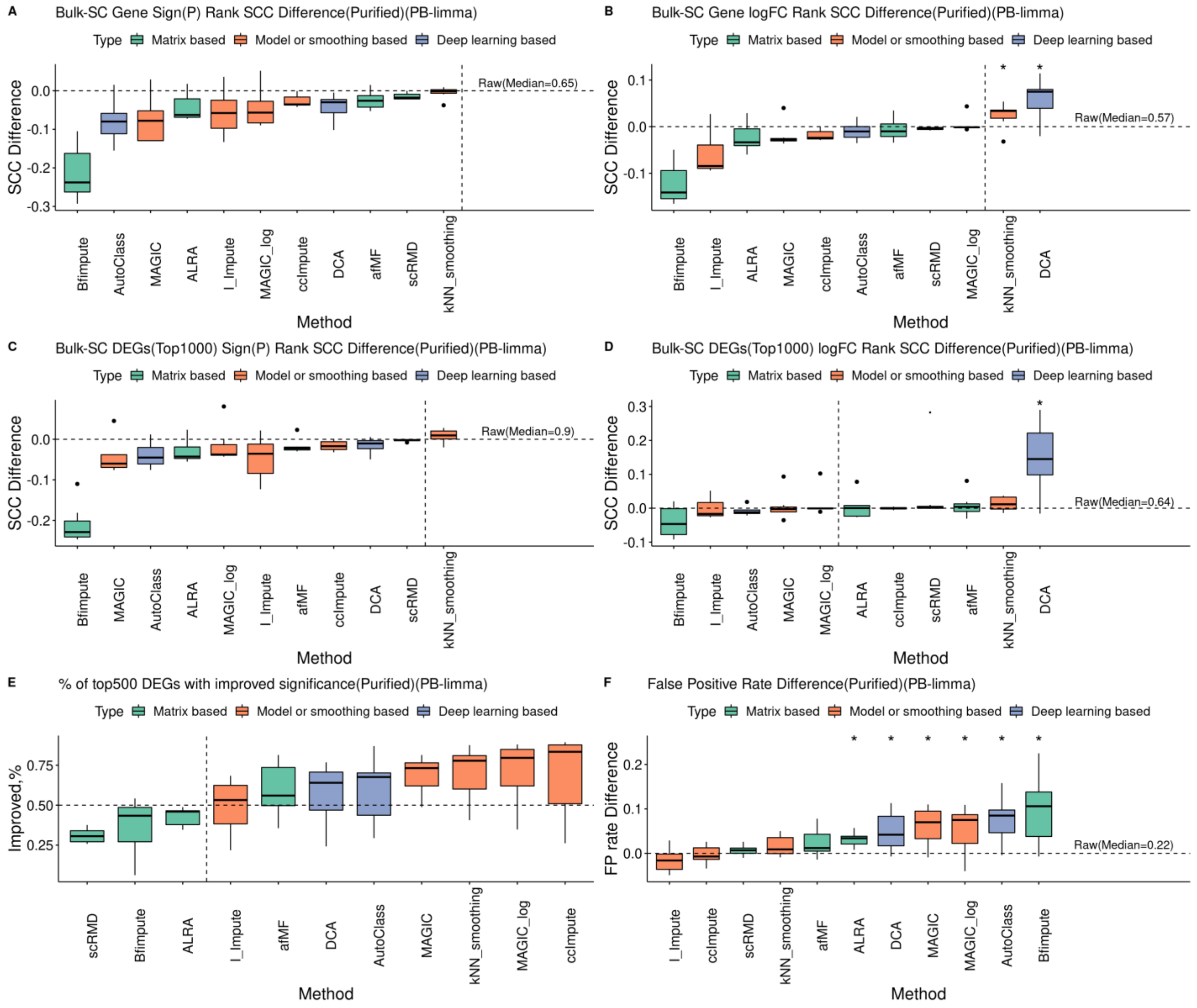

**Figure S14. Heatmaps showing each comparison in MAST-based GSEA**

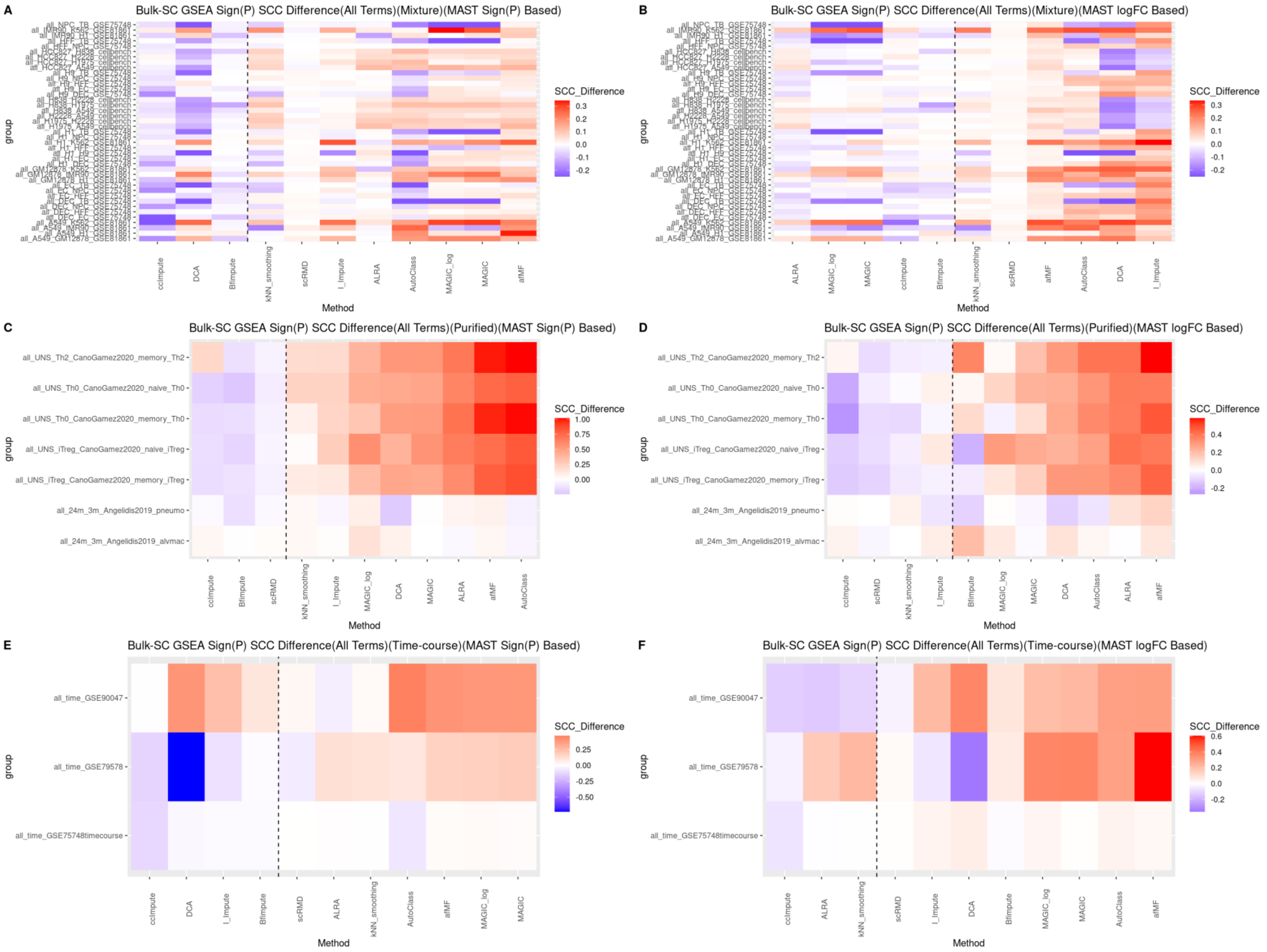

**Figure S15. Performance of imputations on MAST-based GSEA: correlations of P<0.05 GO terms (bulk)**

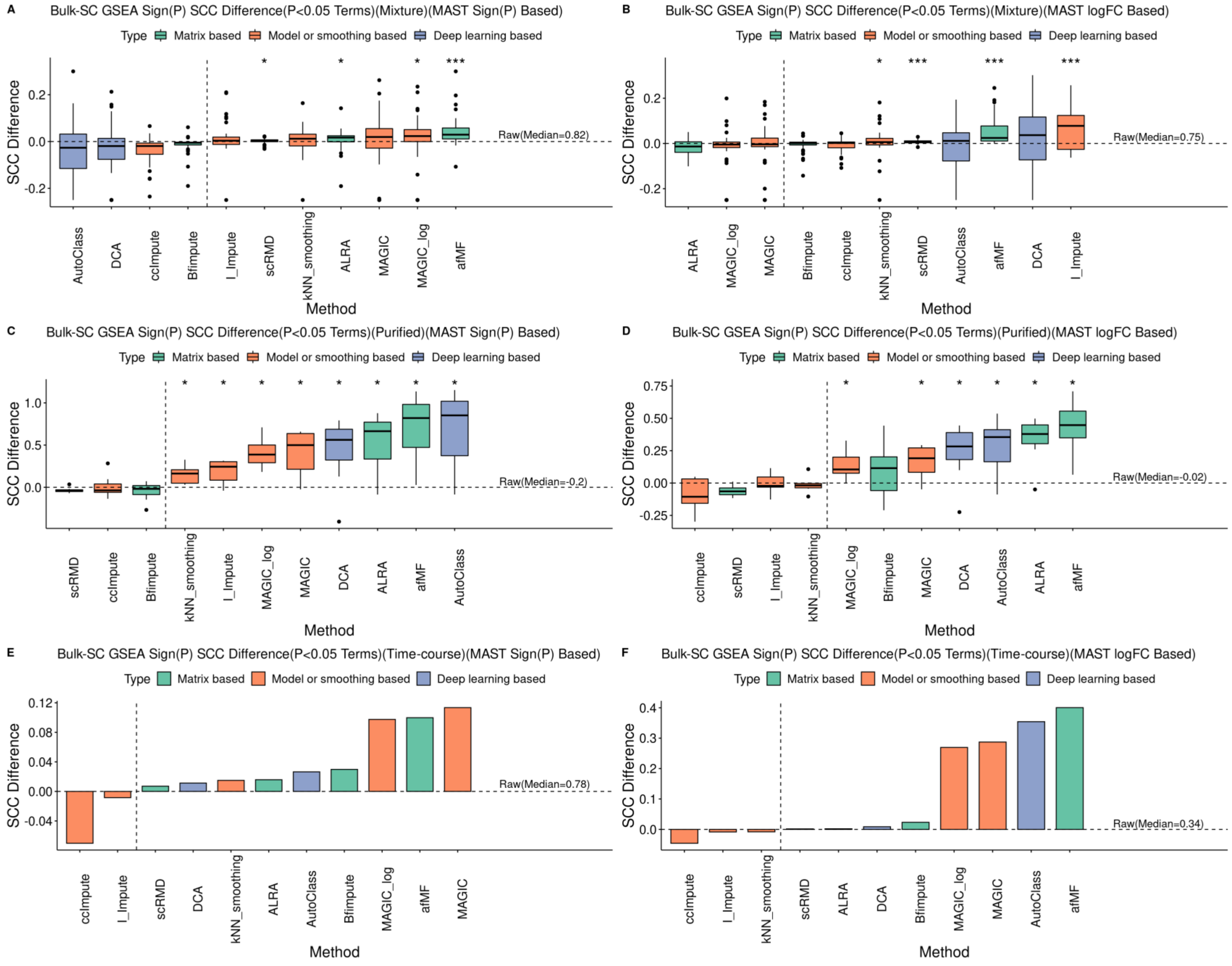

**Figure S16. Performance of imputations on Wilcox Rank Sum-based GSEA**

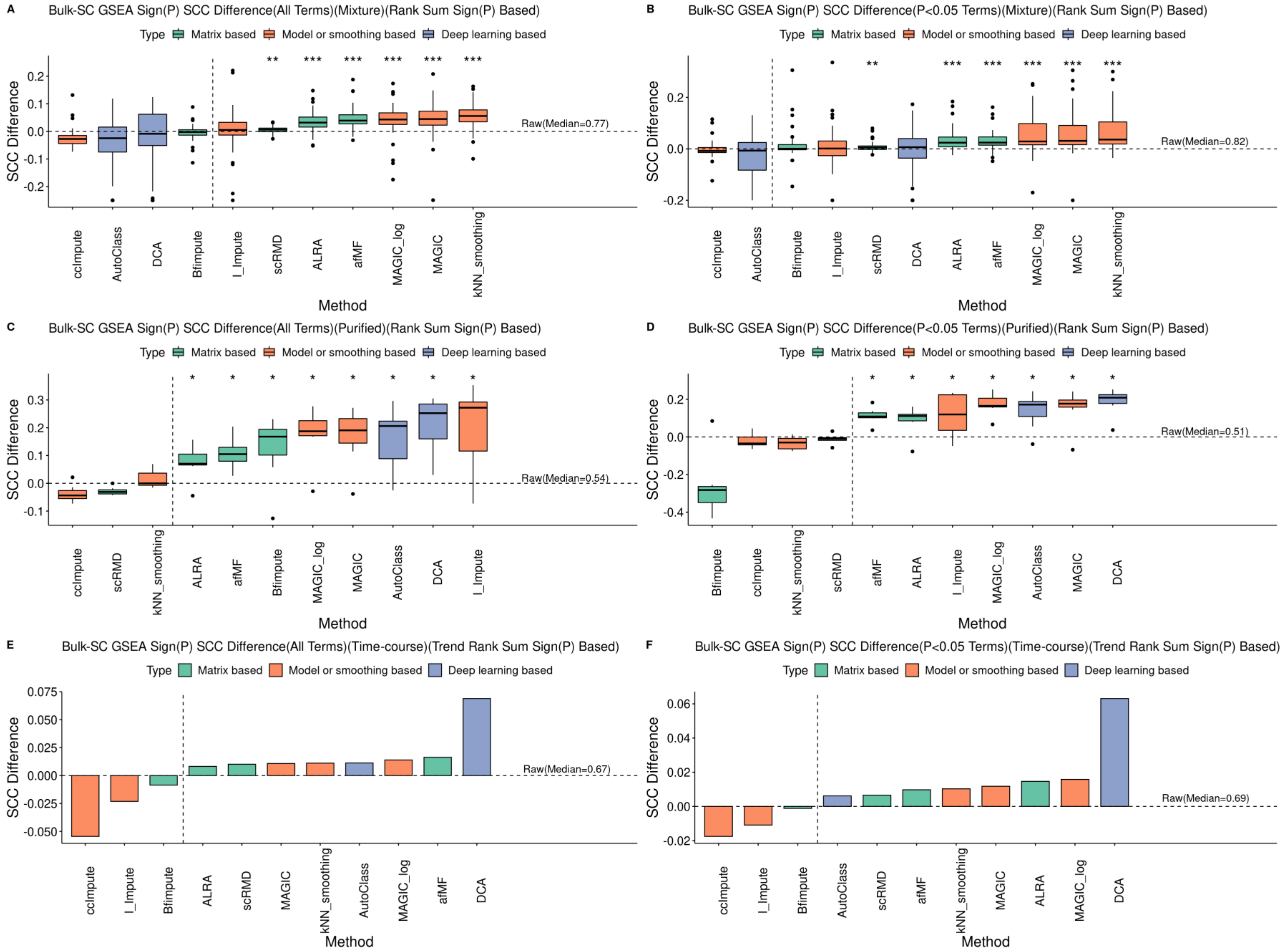

**Figure S17. Performance of imputations on pseudobulk-limma-trend-based GSEA**

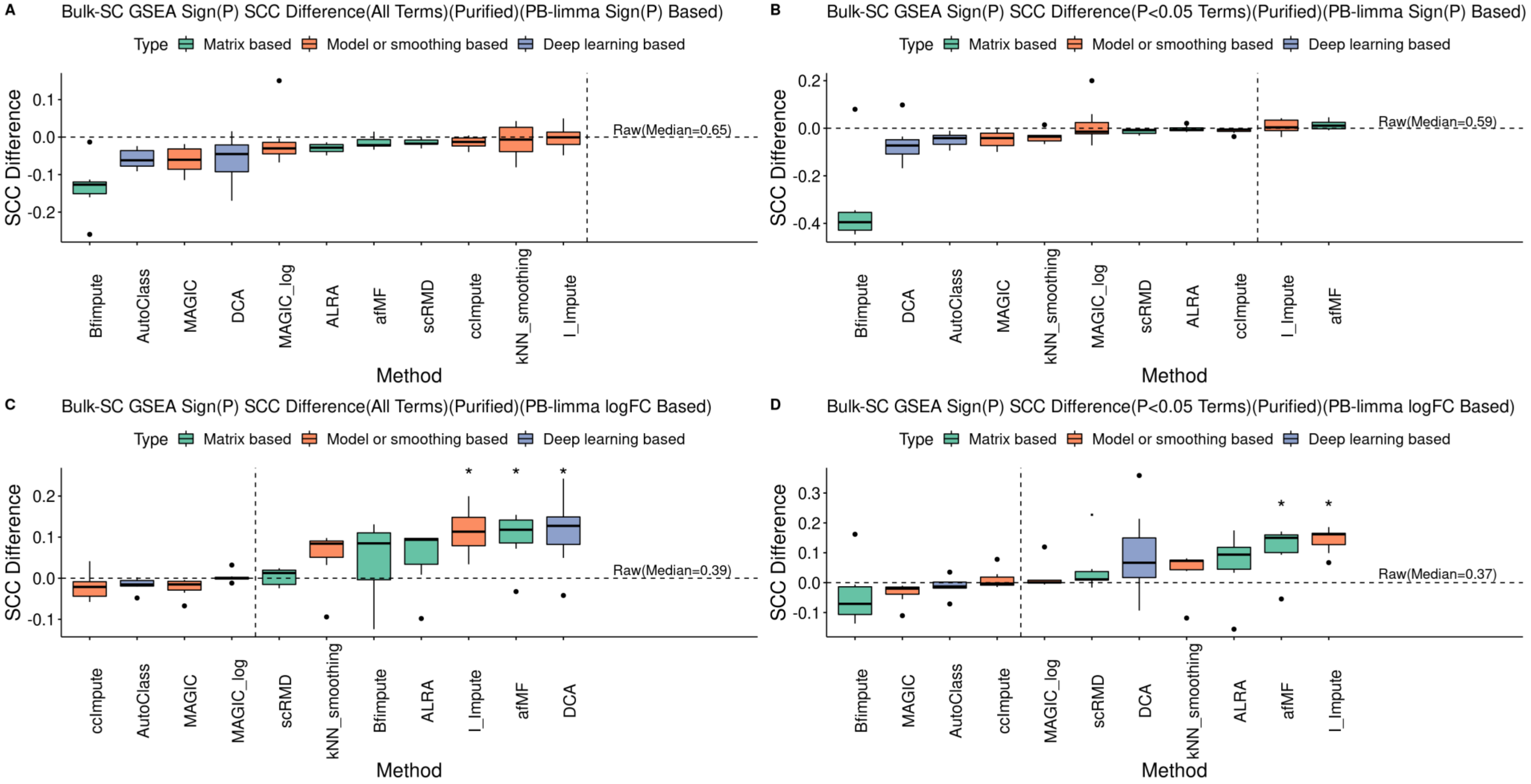
