## Additional file 4 for "Low-Rank Full Matrix Factorization for dropout imputation in single cell RNA-seq and benchmarking with imputation algorithms for downstream applications"

**Figure S18. Performance of imputations on Classification and Biomarker Predictions**

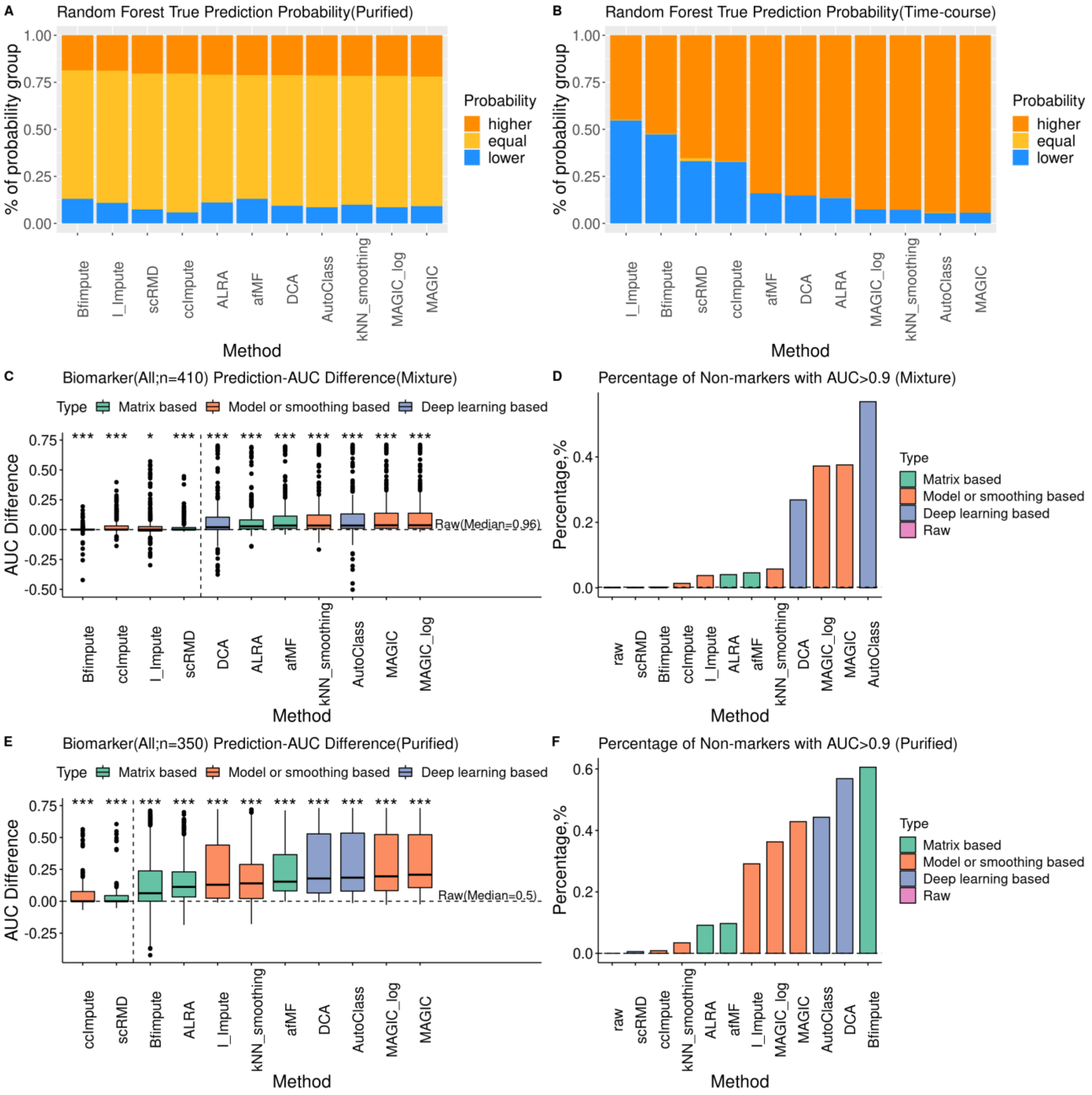

**Figure S19. Performance of imputations on Automatic Cell Type Annotation: SCINA and scType**

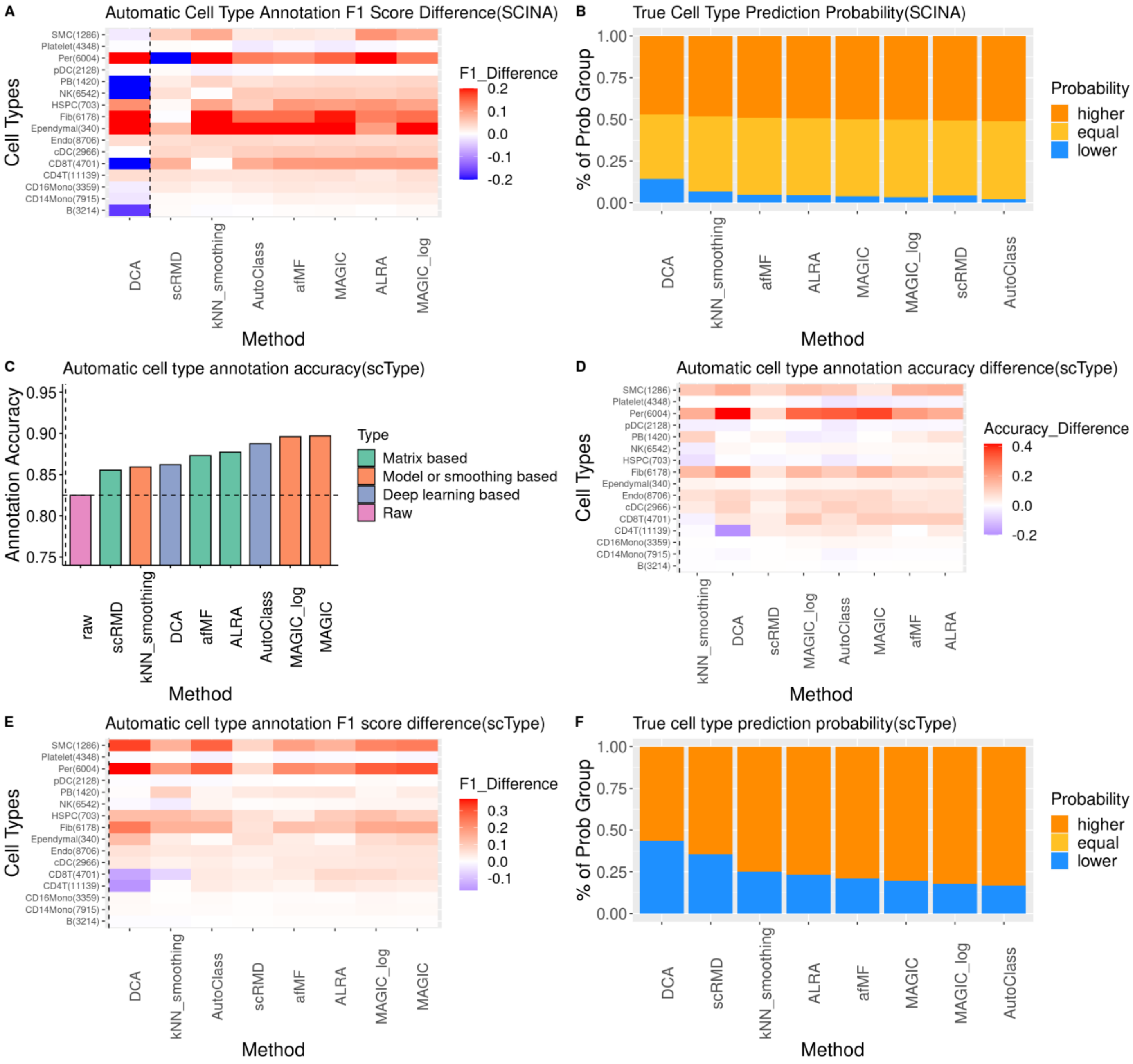
