## Additional file 5 for "Low-Rank Full Matrix Factorization for dropout imputation in single cell RNA-seq and benchmarking with imputation algorithms for downstream applications"

**Figure S20. UMAP plots for CellBench-10X5CL cell types**

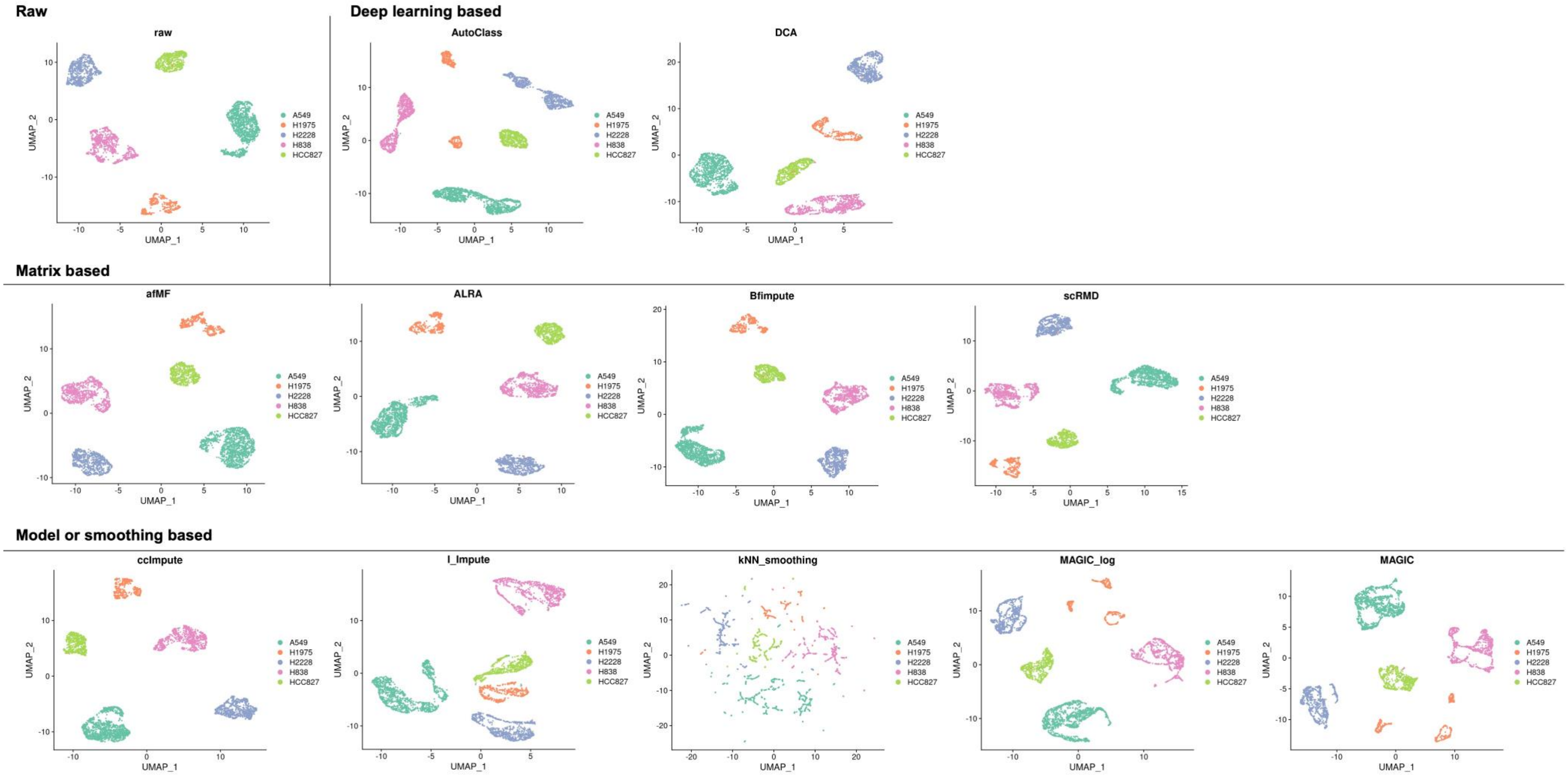

**Figure S21. UMAP plots for CellBench-10X5CL Louvain clusters**

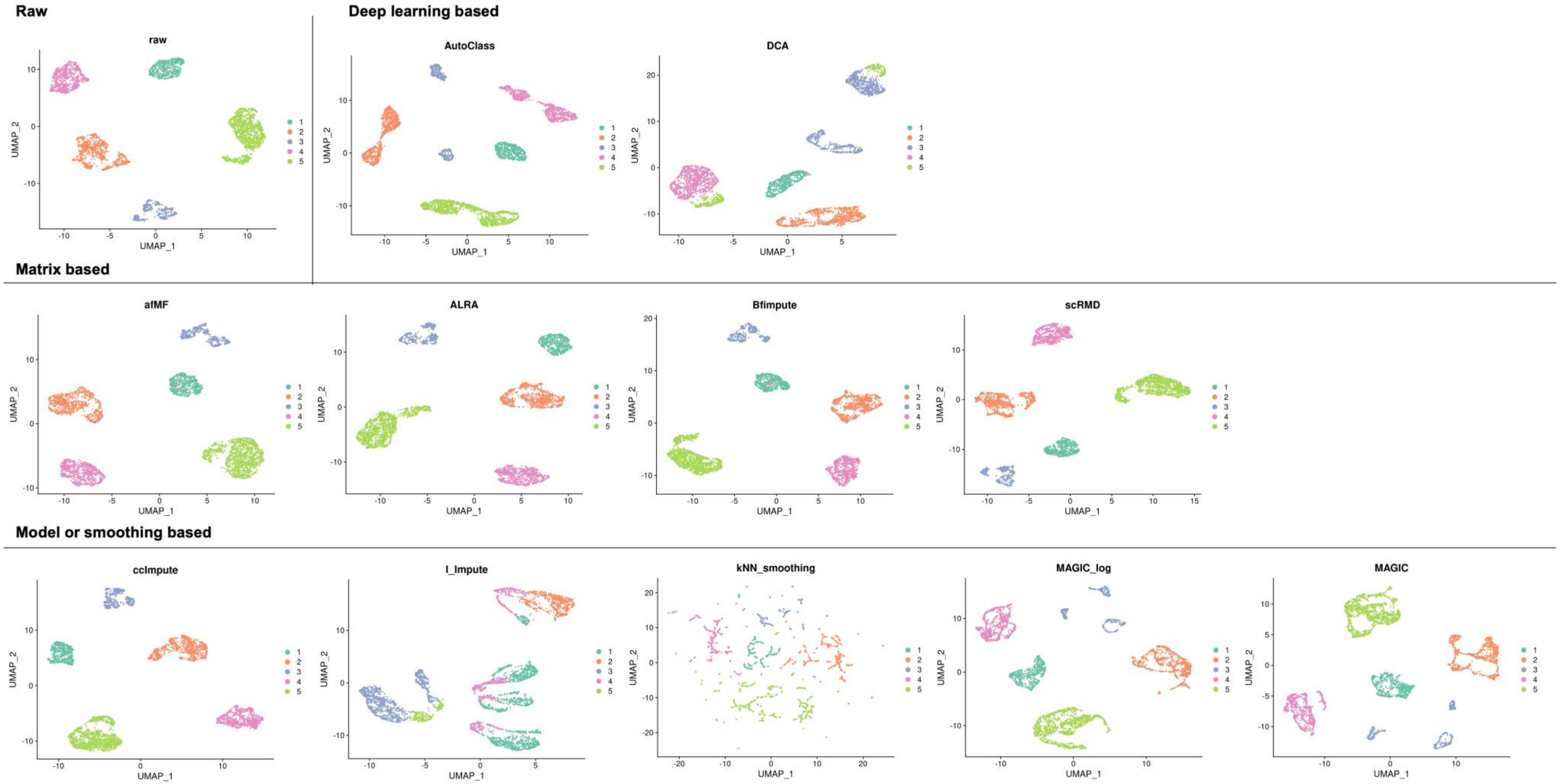

**Figure S22. Performance of imputations on Cell Cycle Dynamics: Seurat predicted cell cycle accuracy and F1 scores**

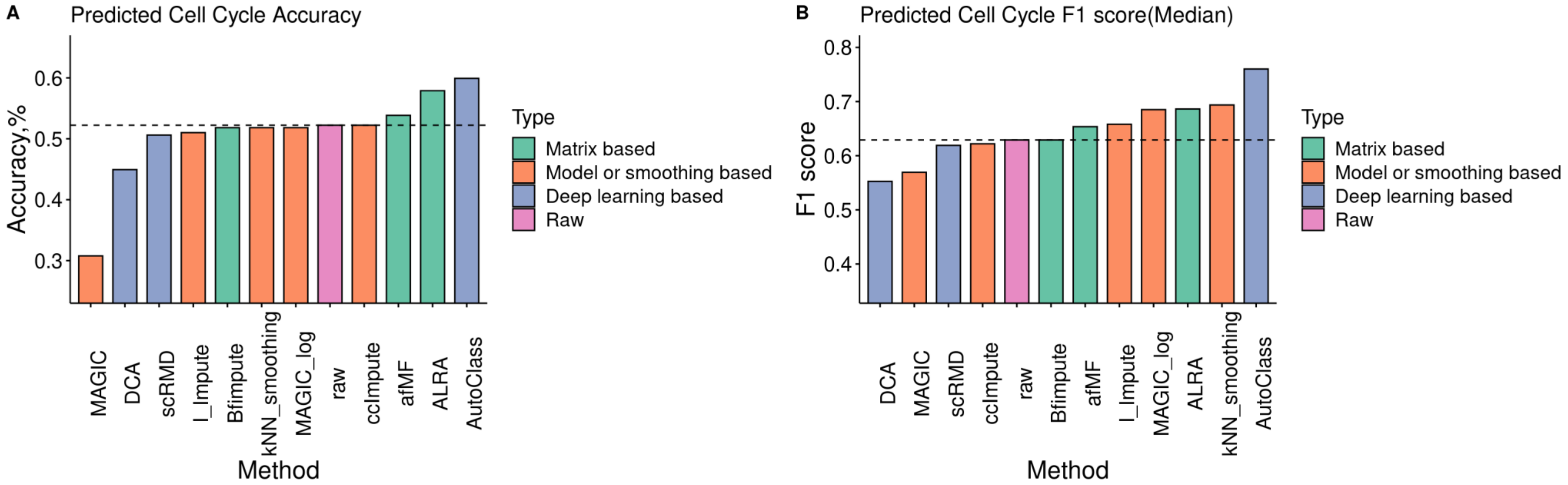
