## Additional file 6 for "Low-Rank Full Matrix Factorization for dropout imputation in single cell RNA-seq and benchmarking with imputation algorithms for downstream applications"

**Figure S23. Heatmaps showing each comparison in DPT SCC and POS**

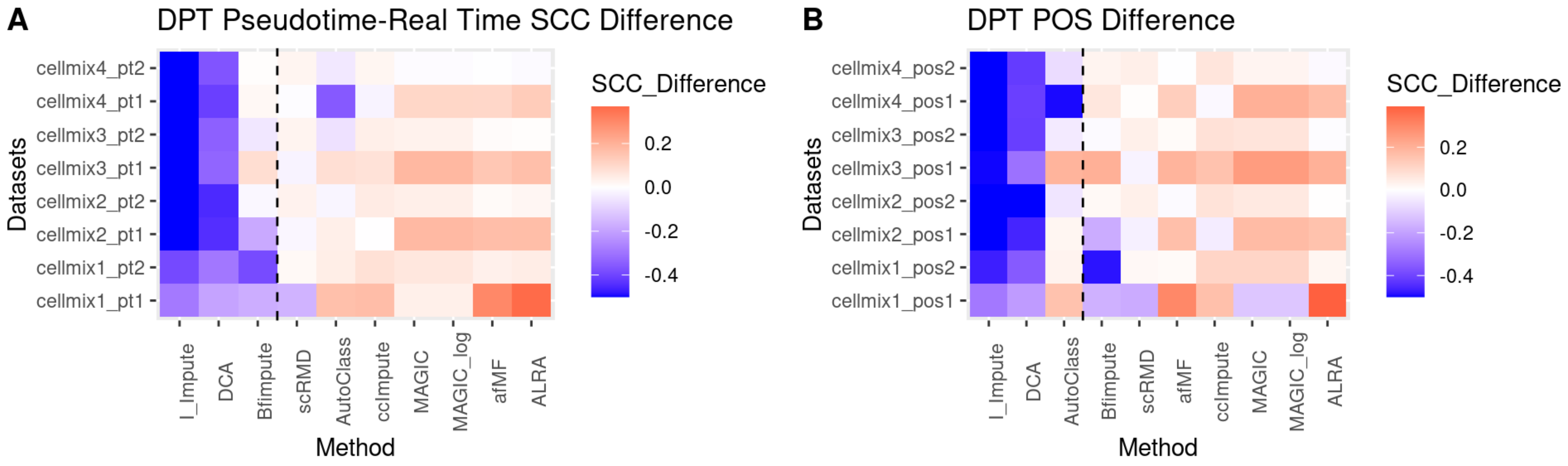

### Figure S24. Diffusion map plots using DPT

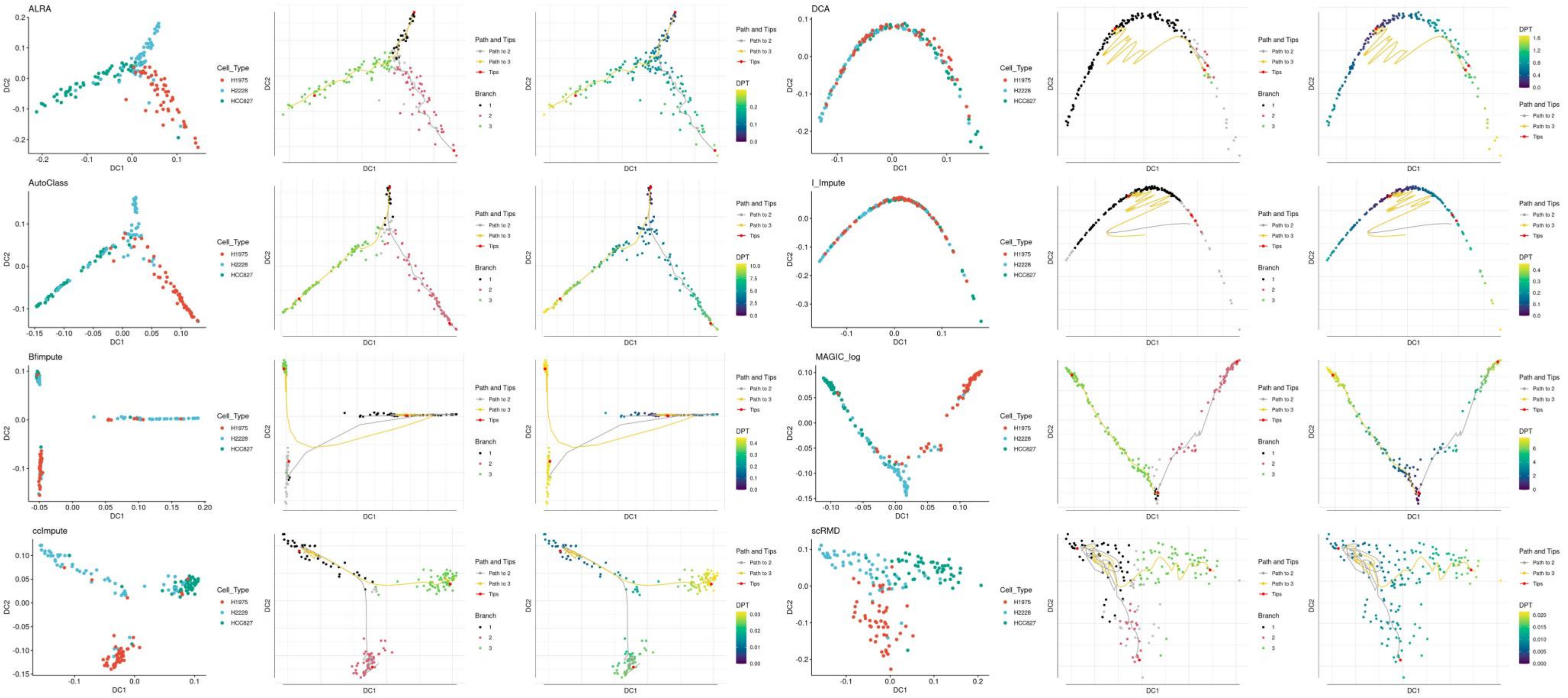

**Figure S25. Performance of imputations on Pseudotime Trajectory Analysis: Monocle3.**

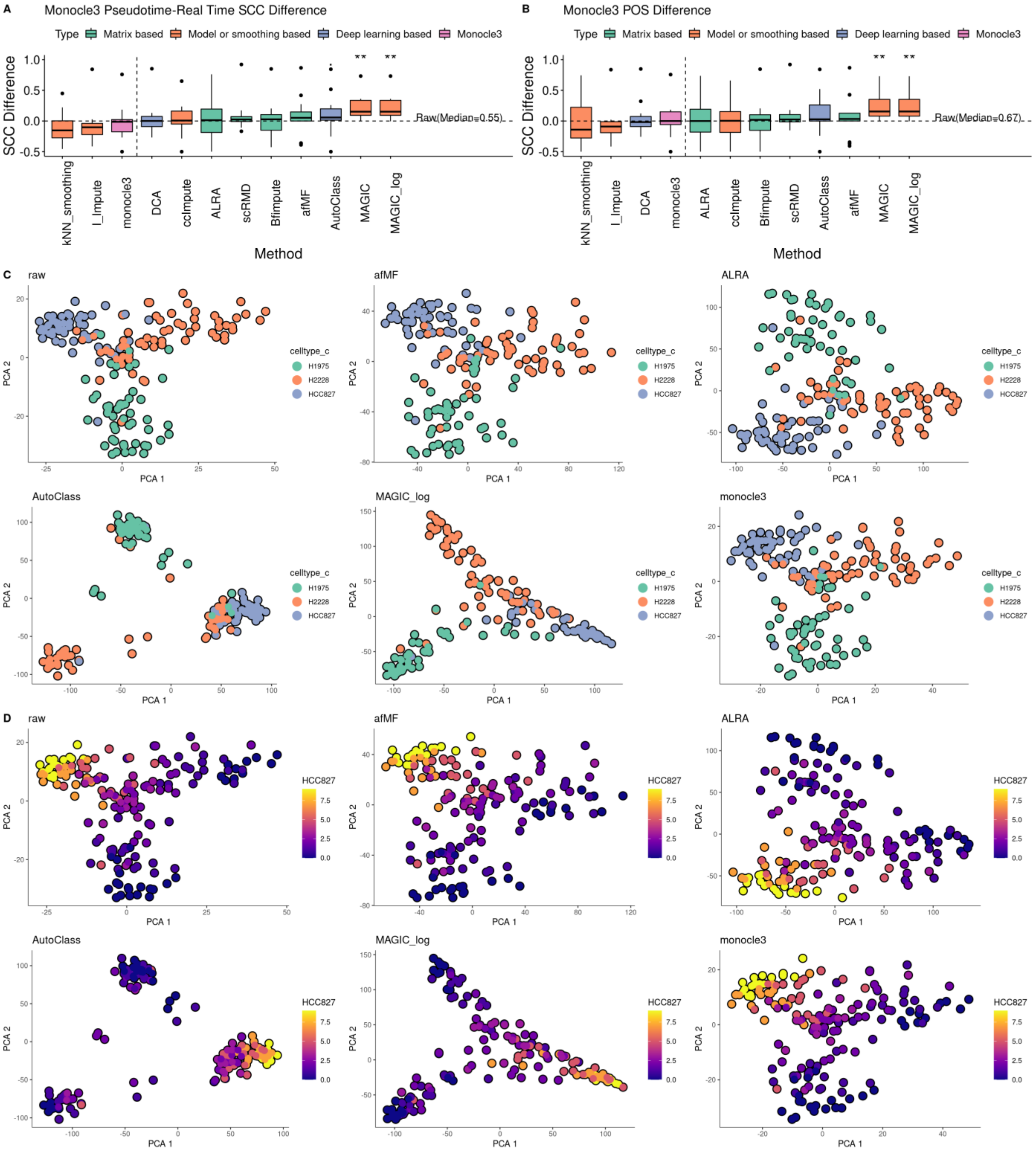

**Figure S26. Performance of imputation on Pseudotime Trajectory Analysis: Slingshot**

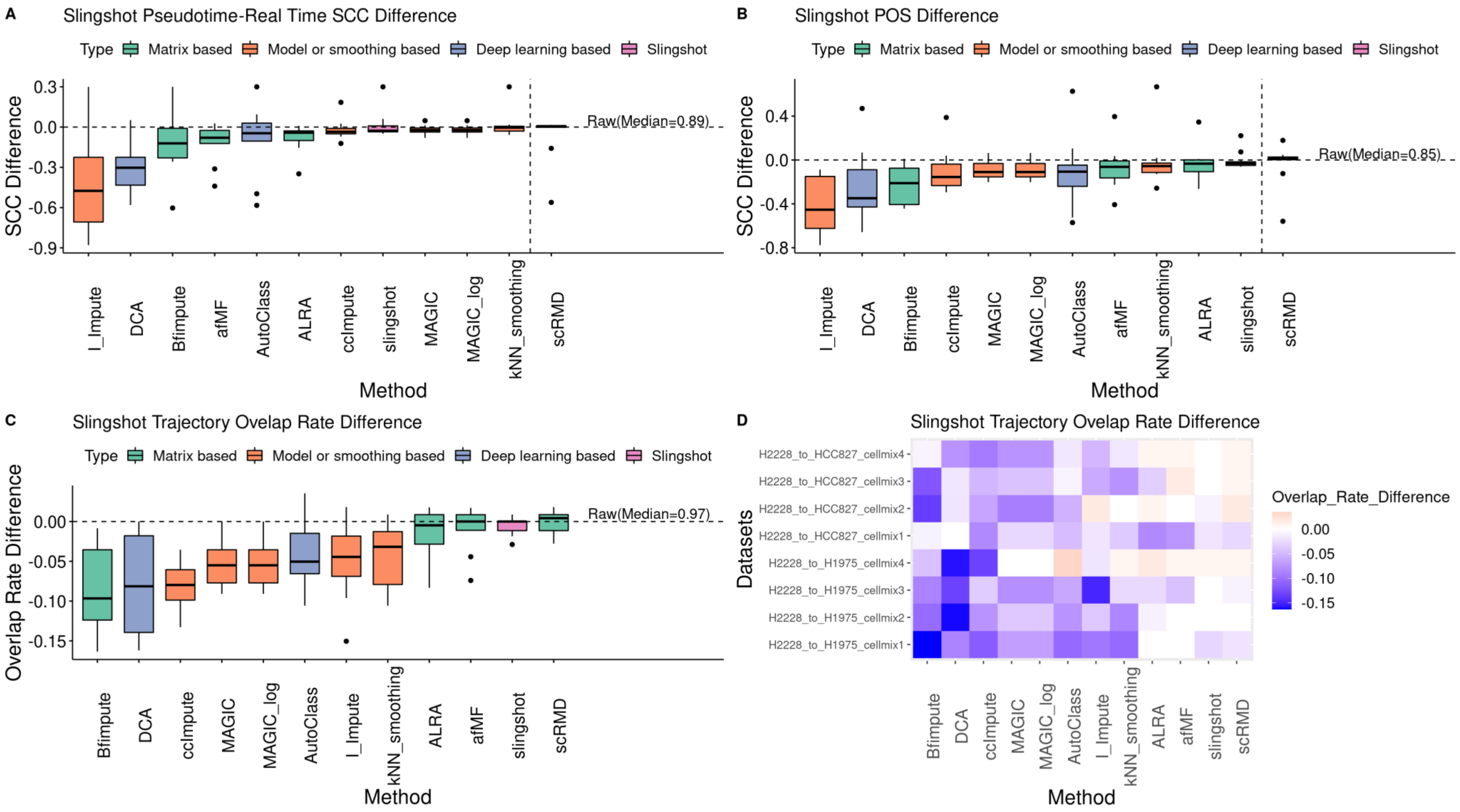
