## Additional file 7 for "Low-Rank Full Matrix Factorization for dropout imputation in single cell RNA-seq and benchmarking with imputation algorithms for downstream applications"

**Figure S27. Comparisons of all the imputation-identified regulons (Z-score>3) across all the cell types**

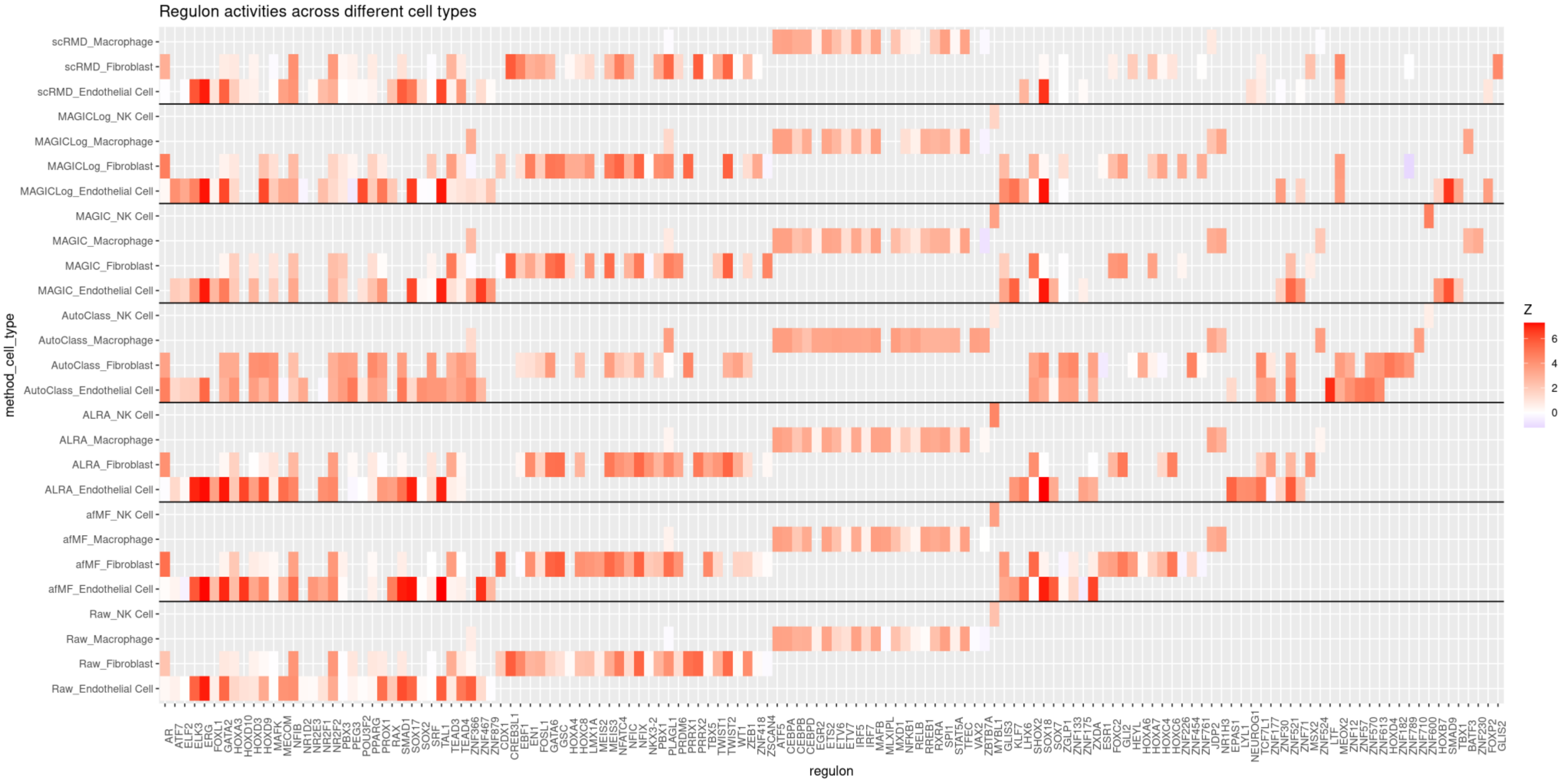

**Figure S28. Comparisons of ligand-receptor cell-cell communications across different imputations using CellPhoneDB**

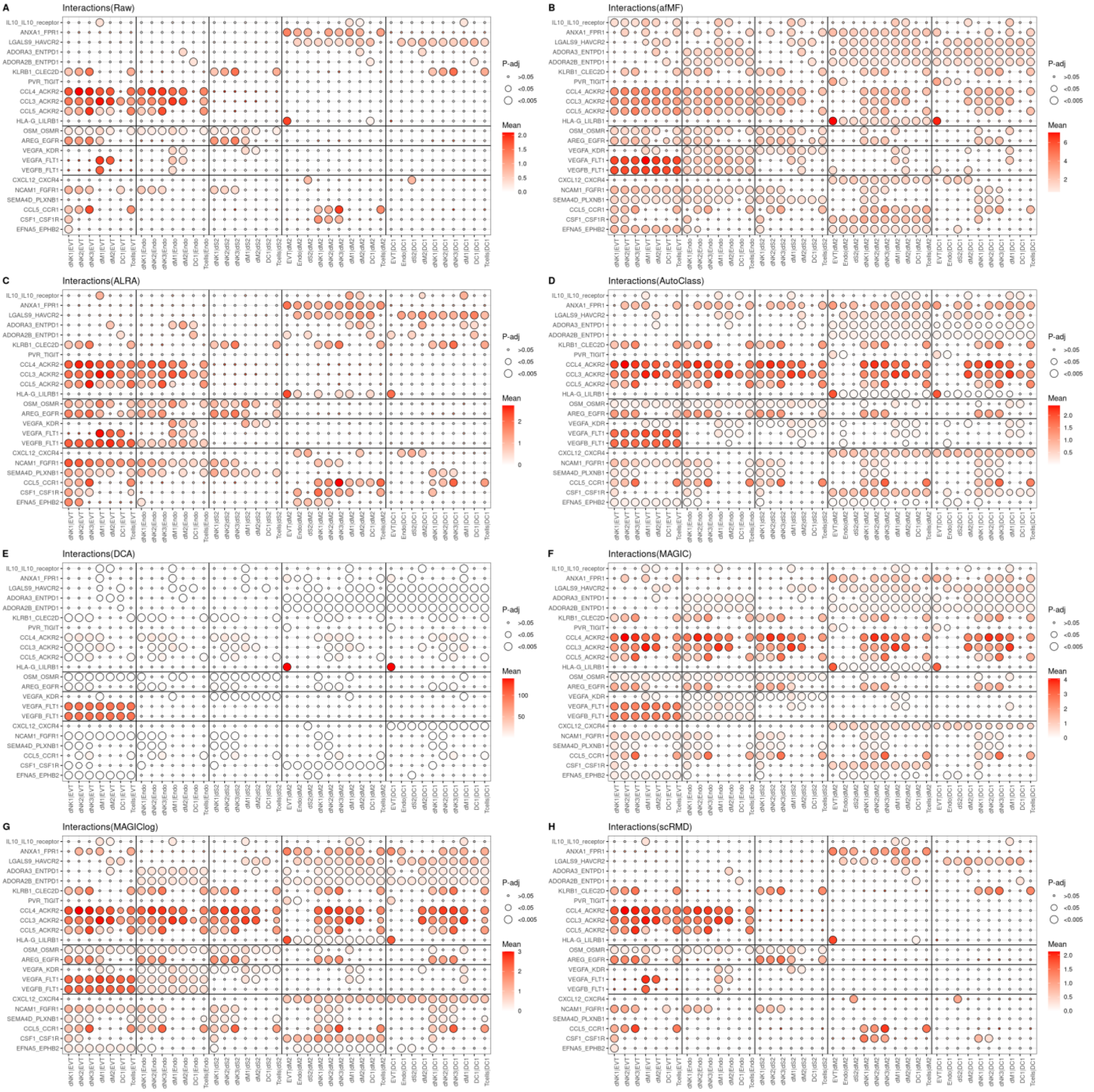

**Figure S29. Performance of imputations on cell-cell communications using CellPhoneDB**

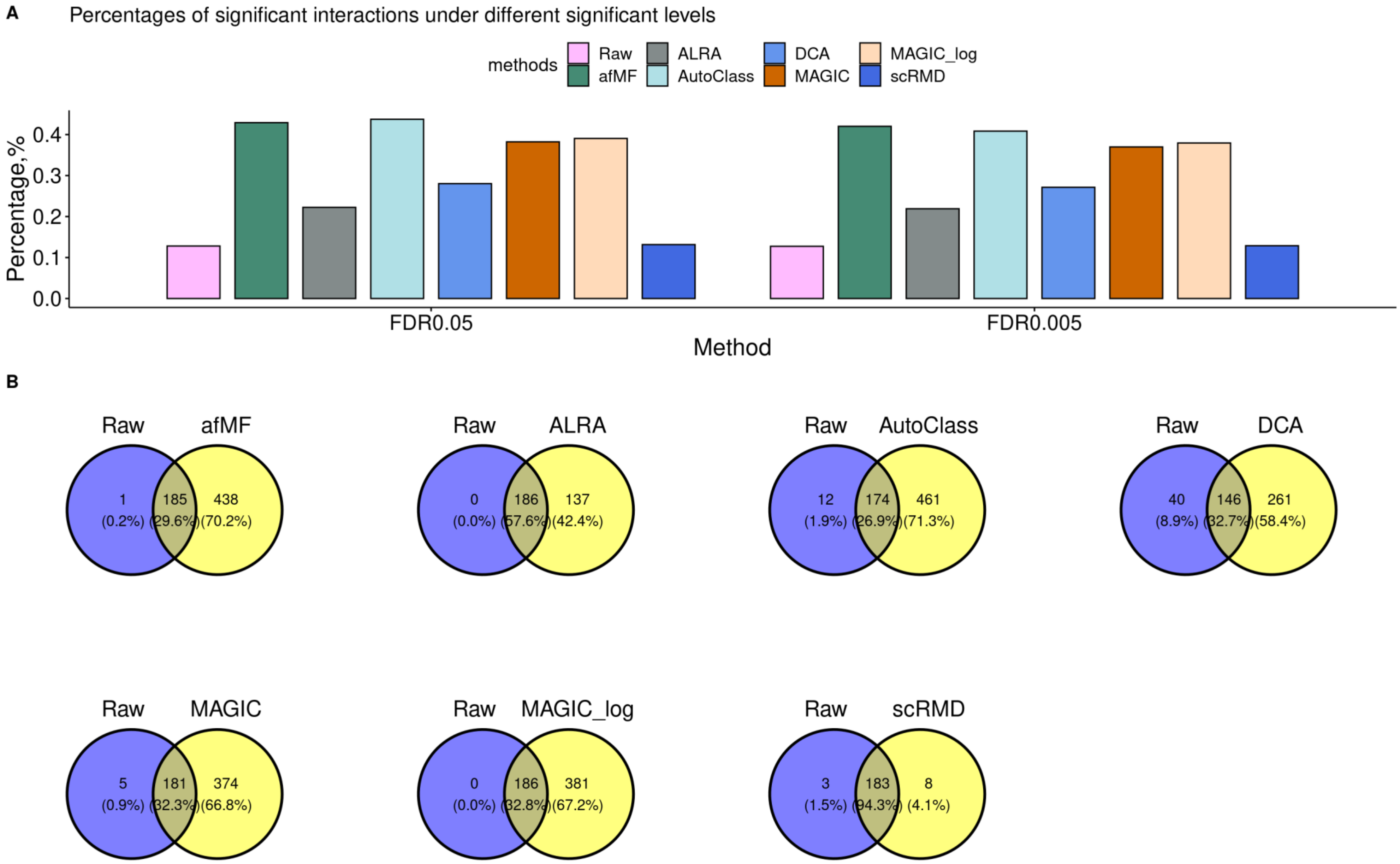

**Figure S30. Performance of imputations on cell-cell communications using CellChat**

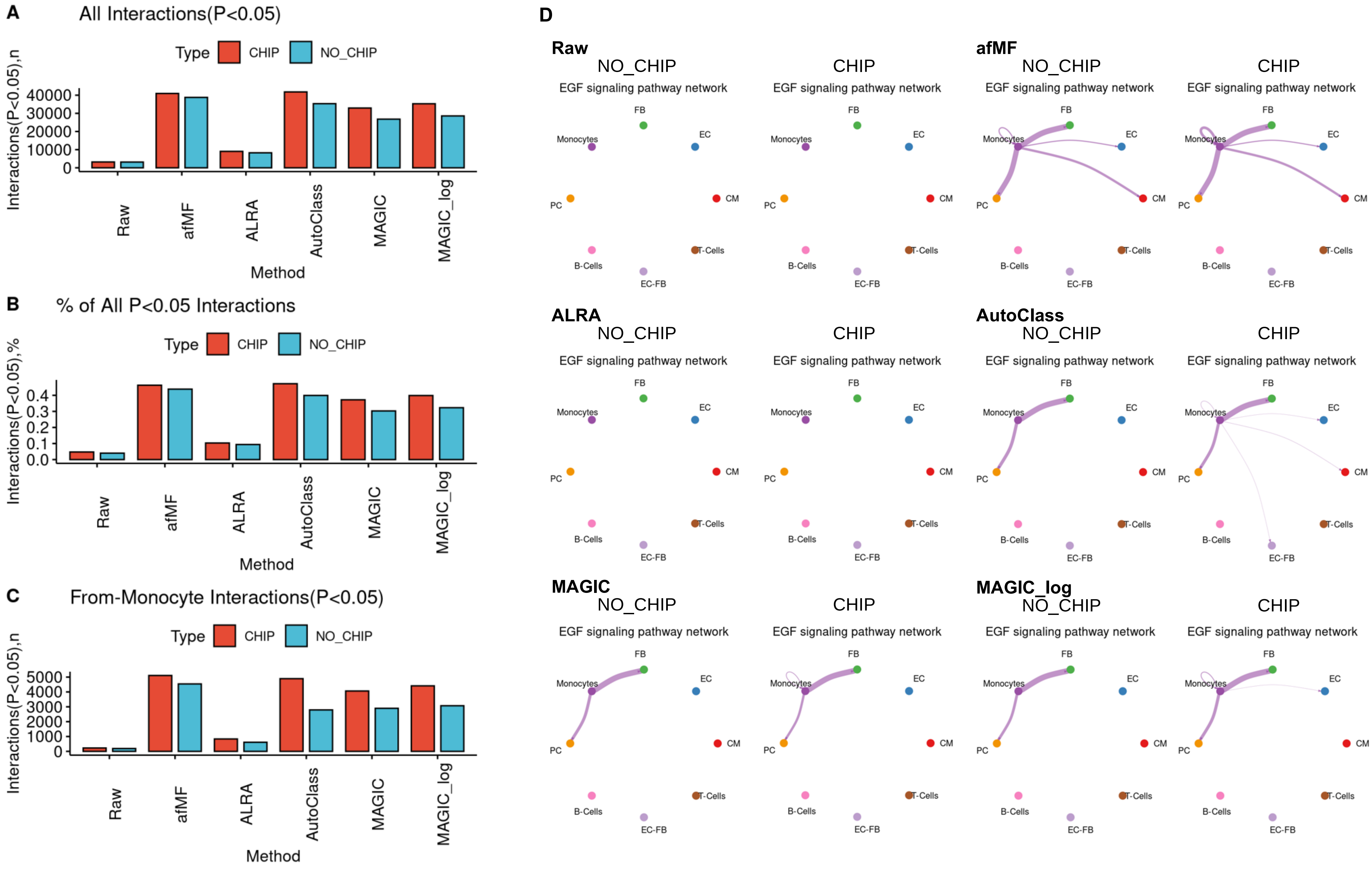
