## Additional file 8 for "Low-Rank Full Matrix Factorization for dropout imputation in single cell RNA-seq and benchmarking with imputation algorithms for downstream applications"

**Figure S31. Performance of imputations in various supporting analysis and heatmaps showing each comparison**

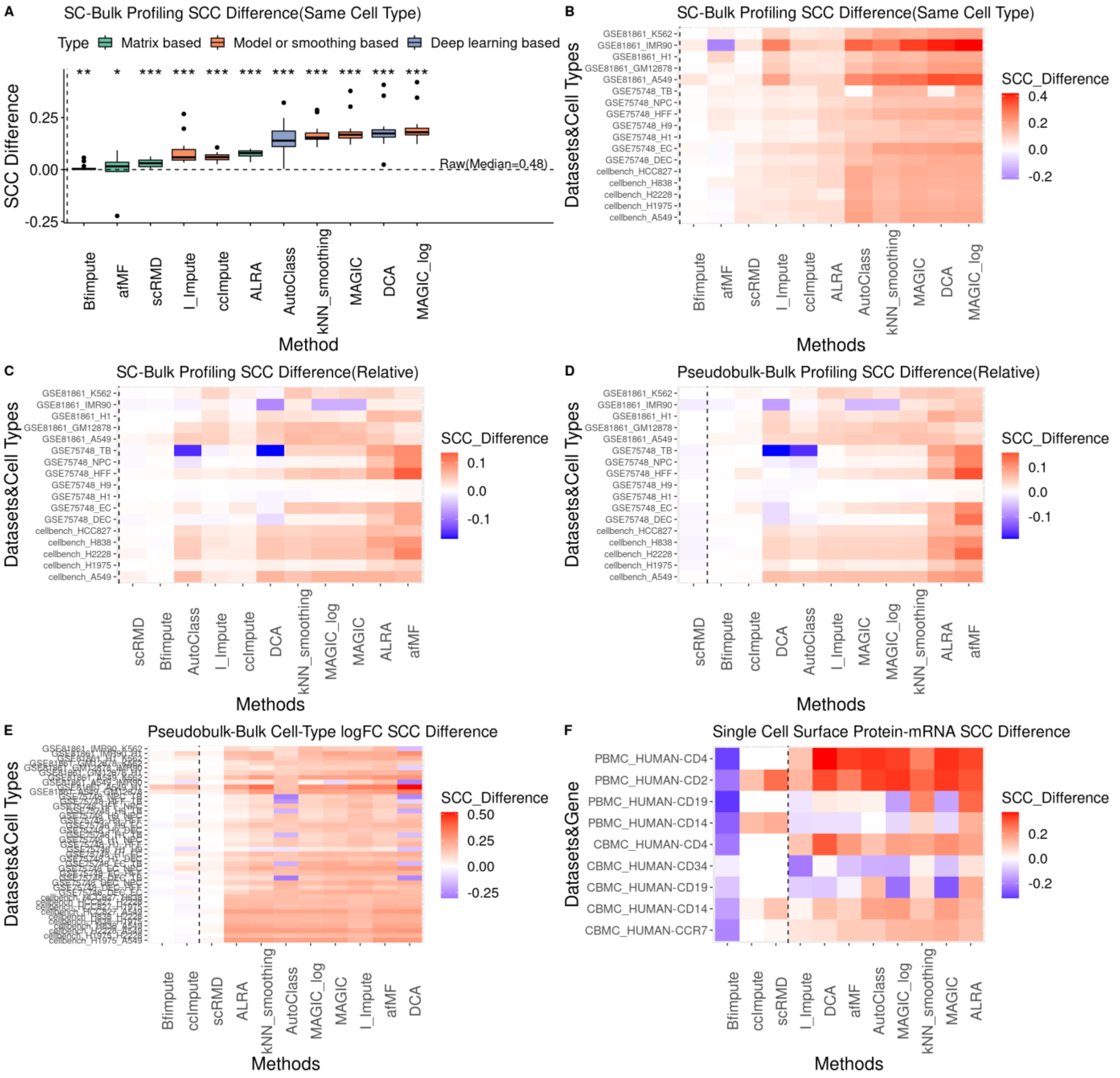
