## Additional file 9 for "Low-Rank Full Matrix Factorization for dropout imputation in single cell RNA-seq and benchmarking with imputation algorithms for downstream applications"

**Figure S32. Performance of imputations on simulated data with ground truth (main discovery)**

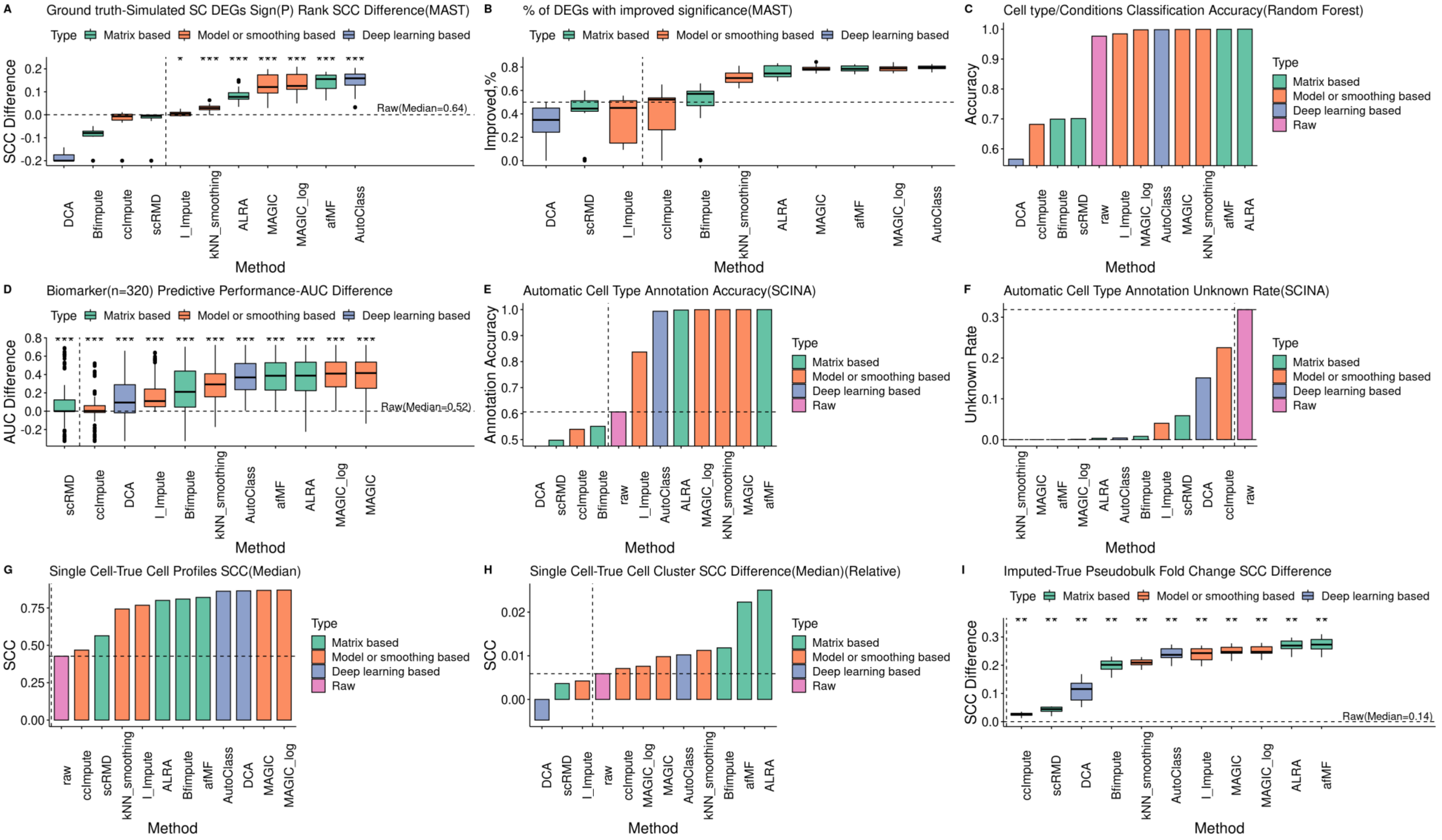


The simulated datasets with ground truth were generated using Splatter (Mock90) and SplatPop (SplatPop90). The ground truth (e.g., known DEGs, groups, matrix) instead of bulk data were used as gold standard in these evaluations. Some metrics were subtracted by the values of the raw results. Extreme values have been limited to a cutoff value for better visualization. The evaluations included: (**A-B**) Differential Expression Analysis; (**C**) Classification; (**D**) Biomarker Prediction, (**E-F**) Automatic Cell Type Annotation; Clustering (Supplementary) and (**G-I**)

Imputed SC-Ground Truth Similarity (Imputed Cell-True Cell; Imputed Cell-True Cell Cluster; Imputed Pseudobulk-True Pseudobulk).

**Figure S33. Performance of imputations on simulated data with ground truth (supplementary results with rank sum test and pseudobulk-limma-trend)**

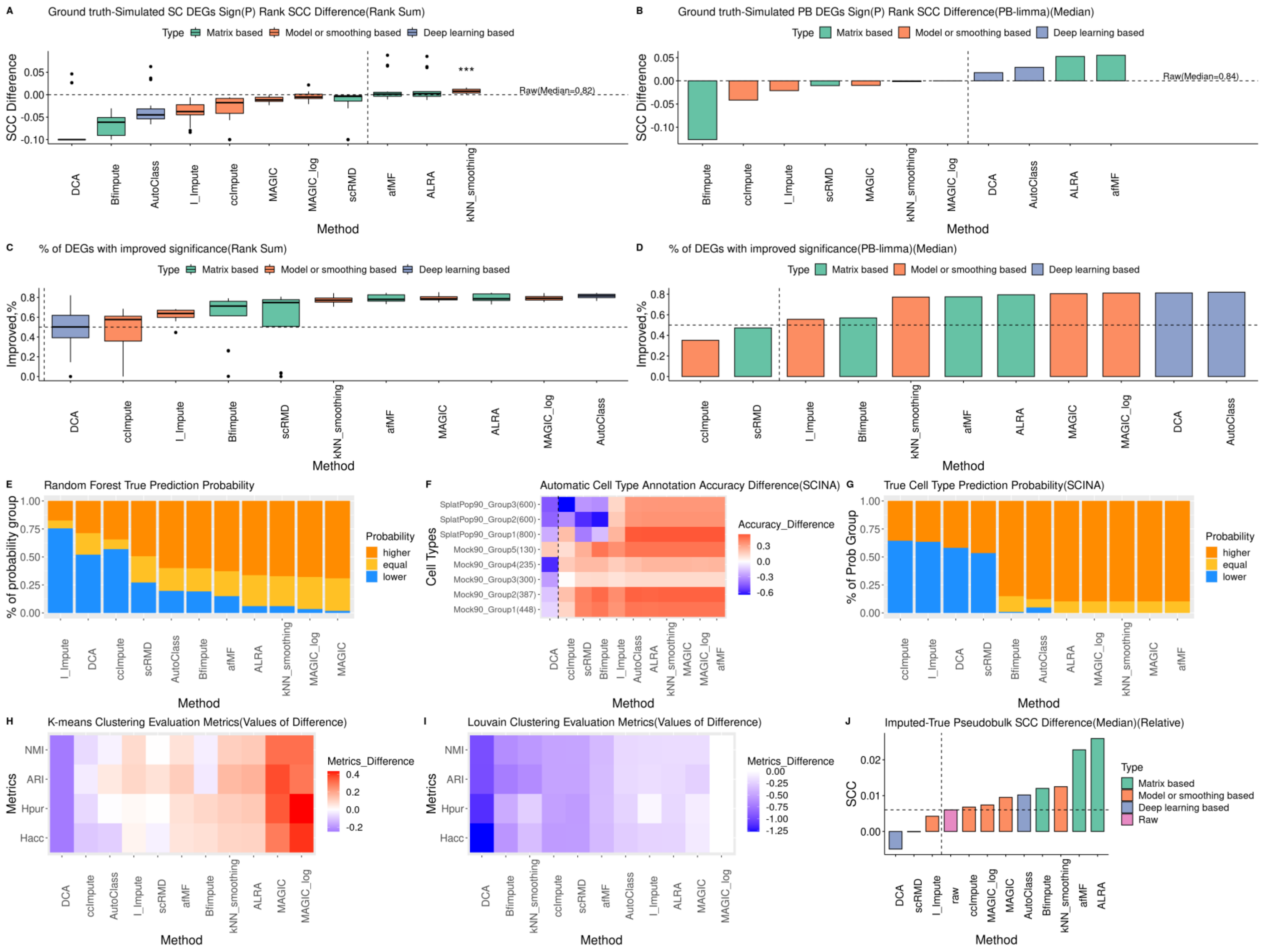

**Figure S34. Cell-Cell Correlation heatmaps in different imputations using simulated data (Mock90)**

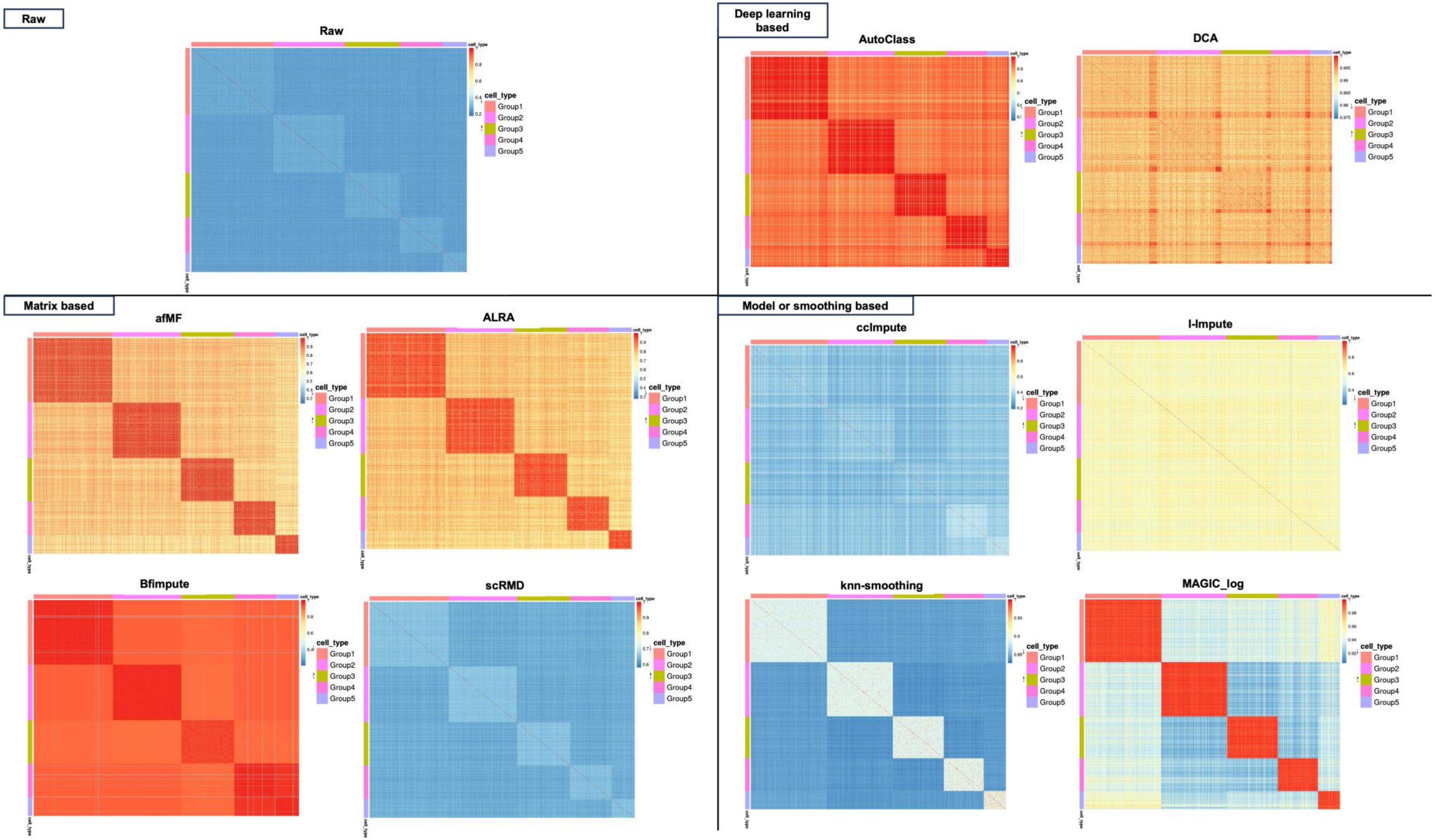
