## Additional file 10 for "Low-Rank Full Matrix Factorization for dropout imputation in single cell RNA-seq and benchmarking with imputation algorithms for downstream applications"

**Table S1. Summary of benefits and drawbacks of imputations with various applications**

| Methods | Matrix theory based | Model or smoothing based | Deep learning based |
|---|---|---|---|
| **Differential expression analysis** | | | |
| - Wilcox rank sum | -only afMF enhanced | -no enhancement or inferior | -no enhancement or inferior |
| - MAST | -only afMF enhanced | -MAGIC slightly enhanced | -no enhancement or inferior |
| - Pseudobulk-limma-trend | -no enhancement or inferior | -no enhancement or inferior | -no enhancement or inferior |
| **Gene set enrichment analysis (GSEA)** | | | |
| - -sign $\log_{10}P$ based | -only afMF enhanced | -only MAGIC enhanced | -no enhancement or inferior |
| - logFC based | -only afMF enhanced | -no enhancement or inferior | -only AutoClass enhanced |
| **Cell type classification and biomarker prediction** | -enhanced | -enhanced | -enhanced |
| **Automatic cell type annotation** | | | |
| - SCINA | -enhanced | -enhanced | -enhanced; except DCA for some cell types |
| - scType | -enhanced | -enhanced | -enhanced; except DCA for some cell types |
| **Cell clustering and cell cycle dynamics** | | | |
| - Louvain | -only afMF enhanced | -MAGIC_log & ccImpute enhanced | -AutoClass enhanced |
| - K-means | -afMF, ALRA & scRMD enhanced | -MAGIC enhanced | -no enhancement or inferior |
| **Dimension reduction (PCA & UMAP)** | -showed consistent patterns | -kNN_smoothing & I_impute generated artefact in UMAP | -DCA generated artefact in UMAP |
| **pseudotime trajectory analysis** | | | |
| - DPT | -afMF & ALRA enhanced | -all enhanced except I_impute | -no enhancement or inferior |
| - Monocle3 | -no enhancement | -only MAGIC enhanced | -no enhancement |
| - Slingshot | -no enhancement or inferior | -inferior to using raw data | -inferior to using raw data |
| **Advanced analysis: AUCell & SCENIC** | -afMF enhanced | -can generate false positives | -can generate false positives |
| **Advanced analysis: CellPhoneDB and CellChat** | -generated a lot of interactions, likely to be false positives | -generated a lot of interactions, likely to be false positives | -generated a lot of interactions, likely to be false positives |
| **Supporting analysis** | | | |
| - SC-Bulk profiling similarities | -afMF & ALRA enhanced | -all enhanced | -no enhancement |
| - Surface Protein-mRNA correlation | -afMF & ALRA enhanced | -knn_smoothing enhanced | -all enhanced |

**Table S2. Performance of the two matrix theory algorithms: ALRA and afMF**

| **Methods** | **ALRA** | **afMF** |
|---|---|---|
| **Dropout elimination for HK genes and cell type marker genes** | Incomplete | Complete |
| **DE analysis** | Only improved in purified datasets | Improved; better than ALRA |
| **GSEA** | Only improved in purified datasets | Improved; better than ALRA |
| **Classification and biomarker prediction** | Improved | Improved; generally better than ALRA |
| **Automatic cell type annotation** | Improved; the extent of improvement over raw data is similar | |
| **Cell Clustering and Dimension reduction** | Improved in K-means algorithm | Improved in both Louvain and K-means algorithms; better than ALRA |
| **Dimension reduction (PCA & UMAP)** | Decent patterns; separate cell types well; similar | |
| **Pseudotime trajectory analysis by DPT** | Improved; the extent of improvement over raw data is similar | |
| **Pseudotime trajectory analysis by Monocle3** | No improvement | |
| **Pseudotime trajectory analysis by Slingshot** | Incompatible; worse than raw log-normalization | |
| **Advanced analysis: AUCell and SCENIC** | Improved; the extent of improvement over raw data is similar | |
| **Advanced analysis: CellPhoneDB and CellChat** | Incompatible; produced false positive interactions | Incompatible; produced even more false positive interactions than ALRA |
| **SC-Bulk profiling similarities** | Improved | Improved; better than ALRA |
| **Surface Protein-mRNA correlation** | Improved | Improved; worse than ALRA |
| **Running time** | Very quick | More running time needed than ALRA but is acceptable (e.g., for 50,000 cells). |
| **Memory usage** | Acceptable | |
